# High-throughput functional profiling and evolutionary covariation analysis of entire riboswitch sequences

**DOI:** 10.1101/2025.11.18.689055

**Authors:** Laura M. Hertz, Anibal Arce, Elena Rivas, Julius B. Lucks

## Abstract

Riboswitches are useful models for revealing how some RNA molecules undergo dynamic rearrangements of their structures to perform cellular functions. A great deal is known about the structure of riboswitch ligand-binding aptamer domains through evolutionary sequence covariation analysis. However, covariation analysis has been more difficult to apply to riboswitch expression platforms given their large range in cellular functions, and their large sequence diversity. Here, we develop an approach to identify whole transcriptional riboswitch sequences starting from their conserved aptamer domains. We then generate covariation models for the entire riboswitch including the aptamer domain and the expression platform. The method consists of first bioinformatically extending identified aptamer domains to include downstream sequence that could contain an expression platform. Filtering is then performed using either a computational prediction algorithm to identify bacterial intrinsic terminator sequences in the expression platform, or a high-throughput functional assay that uses massive parallel oligo synthesis and next generation sequencing to characterize transcriptional termination of riboswitch candidates as a function of ligand. Filtered sequences are then used to develop full riboswitch sequence covariation models. We developed this approach in the context of the fluoride riboswitch, characterizing 1901 fluoride riboswitch sequences using our high-throughput assay. We find that the prediction filtering approach results in few false positives to identify novel, highly functional fluoride riboswitch variants. Finally, we employ the computational approach to develop covariation models of the ZTP, lysine, and TPP riboswitches and find covariation support for previously published rearrangement mechanisms. Overall, our method represents a new hybrid computational and high throughput experimental approach to characterize large numbers of riboswitch sequences and to generate new covariation models of complete riboswitch sequences, which should expand our understanding of riboswitch mechanism and the evolution of RNA structure dynamics.

## INTRODUCTION

Riboswitches, non-coding RNA structural elements that typically reside in the 5’ untranslated region (UTR) of bacterial genes, are important systems to study from fundamental principles of RNA folding (1–6) to anti-biotic targets (7–10). Riboswitches bind a ligand in the aptamer domain, which influences the adjacent structure of the expression platform (EP) that can regulate transcription (11), translation (12), or RNA degradation (13) in bacteria, and splicing in eukaryotes (14), depending on its sequence. While much is known about how riboswitches bind their ligands (15–20), there is still much to be learned about their structural switching mechanism (21,22).

Fundamental to aptamer discovery is sequence covariation analysis, which takes a sequence alignment as input and decodes the potential for two positions to co-vary through compensatory changes (i.e. an A and a U both changing to a G and a C) that would preserve base pairing (23–25). The result of covariation analysis is a secondary structure model of the RNA, called a covariation ‘motif’, that represents which base pairs are most likely given the input alignment. Aptamer domains are highly structured with the singular function of binding a ligand, making them highly structured and thus accessible for detection through covariation analysis (20,26–28). In contrast, expression platforms, due their sequence diversity have largely been not analyzed using covariation analysis.

To address this gap, we developed an approach to generate a covariation motif of a full riboswitch sequence containing both aptamer and expression platform, using high-throughput methods that filter for one specific gene regulatory function, namely intrinsic termination. Bacterial intrinsic terminators are comprised of a conserved sequence motif: a G-C rich hairpin structure followed by a poly-U track (Figure 1A) (29,30). Many riboswitch ligand classes regulate at the level of transcription, and some have had their cotranscriptional folding pathways well-characterized (6,31–33). Notably, past work generated full riboswitch covariation models for the purine (34) and SAM (35) transcriptional riboswitches.

**Figure 1.**
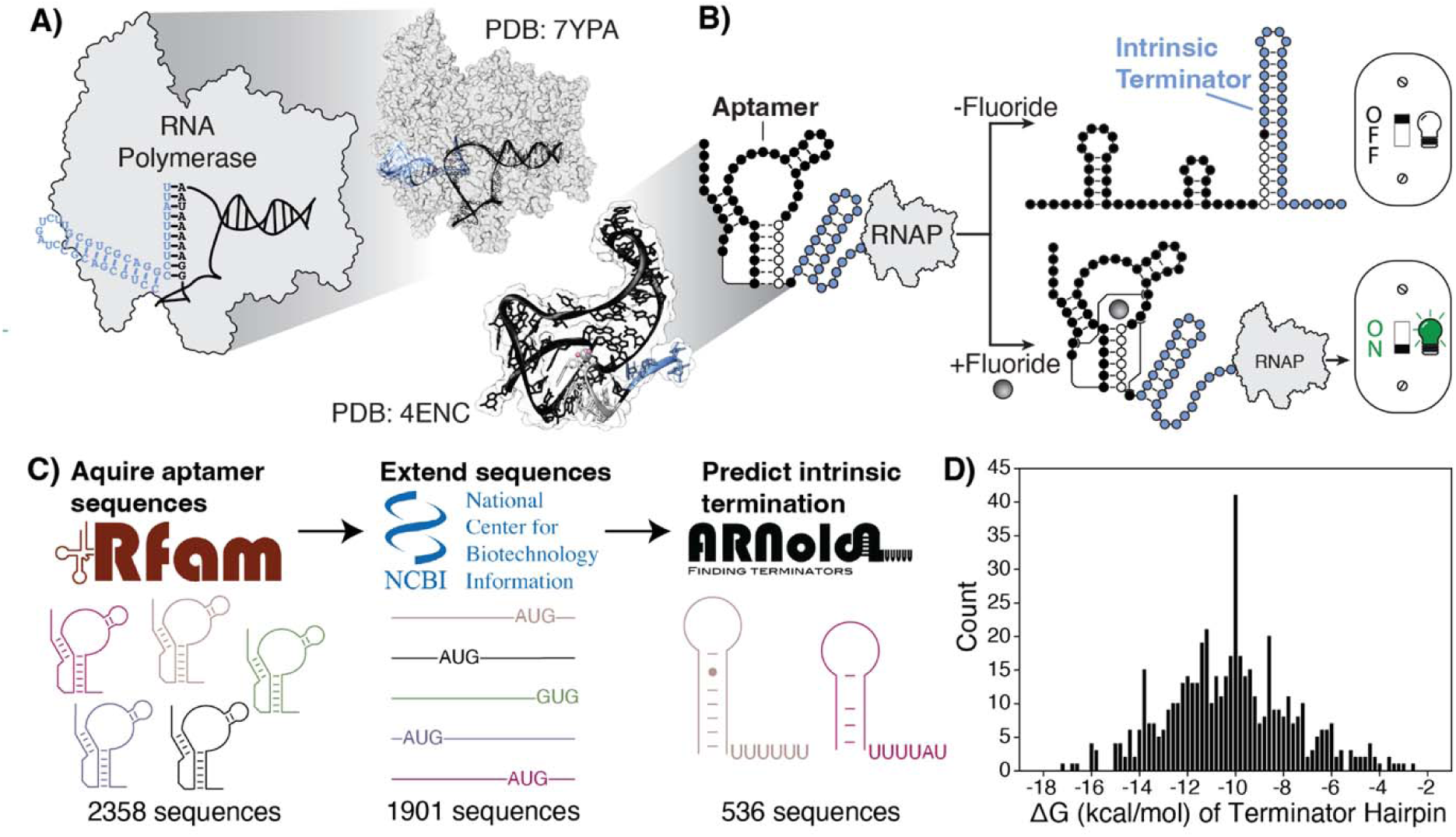
A bioinformatic pipeline to identify transcriptional fluoride riboswitches. **(A) Left:** Structure and schematic of a portion of an intrinsic terminator inside RNA polymerase (grey) exit channel (PDB: 7YPA) with the RNA G-C rich hairpin (blue) and polyU track at the RNA:DNA (blue:black) hybrid. The terminator hairpin secondary structure is modeled from minimum free energy folding predictions from RNAStructure. **Right:** Structure of the fluoride aptamer (PDB: 4ENC), not to scale with the RNAP. **(B)** Schematic mechanism of a transcriptional fluoride riboswitch. In the absence of fluoride, the aptamer (black and white circle nucleotides) undergoes a structural rearrangement to form an intrinsic terminator (white and blue, and singular black, circle nucleotides) to turn off gene expression. In the presence of fluoride, the aptamer is stabilized which prevents rearrangement and allows transcription to continue. RNAP = RNA Polymerase (grey). **(C)** A bioinformatic pipeline that acquires aptamer sequences, accession ID numbers and genomic positions from Rfam family (RF01734), uses genome coordinates to extend the aptamer sequences to capture the adjacent expression platform sequence, and predicts which extended sequences contain an intrinsic terminator motif using the ARNold webserver. **(D)** Histogram of the ΔG (kcal/mol) of the stem-loop terminator structure predicted by ARNold.

However, the approaches used to generate these models used either structural annotation in the alignment (34) or manual curation of the alignment (35) and thus are not widely accessible to other known riboswitch classes. We therefore sought to develop a more high-throughput approach using two complementary approaches, and validated it using the transcriptional fluoride riboswitch (Figure 1B) (4,15,36–40). The method consists of first bioinformatically extending identified aptamer domains to include downstream sequence that could contain an expression platform. Sequences are then filtered using either a computational method to predict the presence of intrinsic terminators, or a high-throughput transcriptional assay that leverages massive parallel oligo synthesis (41) to measure function. Filtered sequences are then used to build covariation models of the entire riboswitch sequence. Using this approach, we were able to build a full covariation model of the fluoride riboswitch that expands upon the known aptamer to cover the transcription terminator, which has conserved overlap with the aptamer. This full model was similar whether the transcriptional terminators were identified computationally or experimentally, with the experimental model having fewer significant covarying basepairs. We also modified our high-throughput transcription assay to explore the functional impact of the transcription elongation factor, GreB, which rescues stalled RNA Polymerases (42), to add to the growing body of work elucidating the role of transcription factors in riboswitch folding and function (39,43,44). Finally, we use the computational filtering approach to build covariation models of the ZTP, lysine, and TPP riboswitches, and show conserved folding mechanisms.

## METHODS AND MATERIALS

### Sequence acquisition and synthesis

Fluoride riboswitch sequences were generated *in silico* by extending known aptamer sequences with downstream genomic context, and concatenating with promoter and 3’ priming site sequences according to the following workflow. First, a FASTA file of all computationally identified sequences of the fluoride aptamer were downloaded from Rfam (45) (RF01734 downloaded 08 May 2023) (see **Data and code availability**). We next calculated the length of the aptamer sequences plus the length of the *E. coli* J23119 consensus promoter sequence (from the registry of standardize biological parts, https://parts.igem.org/Part:BBa_J23119), and a 3’ priming sequence used for oligo pool amplification (Supplementary Table 1). For each aptamer sequence, this total length was then subtracted from the Twist oligo pool length (300 nt), and the remaining number of nucleotides was retrieved from the genome entry for that aptamer sequence from NCBI database via Entrez (Entrez Programming Utilities Help, available at https://www.ncbi.nlm.nih.gov/books/NBK25501/), starting from the 3’ end of the aptamer sequence. The *E. coli* J23119 consensus promoter sequence was then prepended to each aptamer extended sequence, and the 3’ oligo priming site was appended to each sequence. We note that we removed the first five nucleotides from the Rfam sequence entry of the genomically encoded region as we observed that the Rfam database included the 10 nucleotides leading up to the fluoride aptamer (starting at P1) and leader sequences can impact riboswitch function (31,32). The final sequences were exported into a spreadsheet.

Entrez was used again to collect information about what protein the aptamer was potentially regulating and the location of the start codon. The information was added to the sequence information used for the Twist Bioscience sequence pool order (Supplemental_Document_A.xlsx, ‘Bioinformatics’ sheet).

Code was executed through a Jupyter Notebook running python (v. 3.10.9). The prepping_sequences.py script was used to generate the fluoride riboswitch sequences and oligo pool order. The NCBI_CDS.py script was used to identify the potential protein.

### Computationally predicting intrinsic termination

We next classified extended fluoride aptamers by whether they are predicted to undergo intrinsic termination using the online webserver, ANRold (46). To do this, extended sequences were converted into a FASTA file format. The FASTA file was then submitted to the ARNold webserver. The results were the processed to create a FASTA file of fluoride riboswitches post-aptamer sequences between 90 to 110 nts in length that are predicted to undergo intrinsic termination.

Code was executed through the script CDS_ARNold.py on a Jupyter Notebook running python (v. 3.10.9).

### Covariation analysis for computationally predicted fluoride riboswitches

Covariation analysis on fluoride riboswitch terminating variants (see Computationally predicting intrinsic termination) was done through analyzing a multiple sequence alignment with the R-scape (RNA Structural Covariation Above Phylogenetic Expectation) package (24) (v2.0.0.j). A multiple sequence alignment was first created for the aptamer region, another for the post-aptamer sequences, and then both alignments were merged for R-scape analysis.

To create the alignment for the aptamer region, the fluoride aptamer Stockholm alignment file was downloaded from Rfam (RF01734.sto). From the ARNold FASTA output of CDS_ARNold.py, the aptamer region was aligned to RF01734.sto using nhmmer (47), which created the output Predicted_RF01734_profile.sto

To create the alignment for the extended post-aptamer sequences, we took the output from CDS_ARNold.py and Predicted_RF01734_profile.sto to generate a FASTA file of the post-aptamer sequences. The post-aptamer sequences were aligned using Jalview (v. 2.11.3.3) and the accessory MUSCLE with default parameters (48).

The two separate alignments were then merged using the Shelley_Alignment.py script through a Jupyter Notebook running python (v. 3. 10.9). R-scape was used to assess significant covariation on the compiled alignment, and to build a consensus secondary structure informed by covariation using the CaCoFold algorithm (24).

Code was executed through a Jupyter Notebook running python (v. 3.10.9). The Shelley_Alignment.py script was used to assess aptamer alignment, generate the FASTA file for MUSCLE alignment, and merge the aptamer and post-aptamer alignments.

### Covariation analysis for experimentally validated transcriptional fluoride riboswitches

Covariation analysis on fluoride riboswitch variants that were validated to undergo intrinsic termination through next-generation sequencing (see Read analysis) were saved in a FASTA file (see Data and code availability). We then performed a similar alignment and R-scape analysis as in Covariation analysis for computationally predicted fluoride riboswitches with the exception that the post-fluoride aptamer sequences were limited to 26 nts in length to generate the MUSCLE alignment.

The Stockholm file output from running CaCoFold was used to generate a covariation model to align all terminators validated through next-generation sequencing. We used the Infernal package to build a covariation model with cmbuild, calibrated the model with cmcalibrate, and searched all validated terminators with cmsearch. We then generated the covariation model of all validated terminators by running cacofold from R-scape.

### Covariation analysis for additional riboswitch classes

We performed similar riboswitch covariation analysis for the purine (RFAM RF00167), SAM (RFAM RF00162), ZTP (RFAM RF01750), lysine (RFAM RF00168), TPP (RFAM RF00059), and glmS (RFAM RF00083) aptamer families. Sequences were extended following **Sequence acquisition and synthesis**, classified as containing an intrinsic terminator following **Computationally predicting intrinsic termination**, and then subject to analysis following **Covariation analysis for computationally predicted fluoride riboswitches**. During this process, the post-aptamer sequences were limited by length (post-purine aptamer sequences were limited to 45 – 65 nucleotides, post-SAM aptamer sequences to 70 – 80 nucleotides, post-ZTP aptamer sequences to 5 – 35 nucleotides, post-lysine aptamer sequences to 25 - 50, post-TPP aptamer sequences to 40 – 45 nucleotides, and post-glmS aptamer sequences to 30 – 100 nucleotides) to produce a clean alignment. The first pass analysis was restricted to the first 1000 sequences due to limits on the MUSCLE alignment webserver. Once CaCoFold was run on the combined alignment, a new covariation model was calibrated and searched through all extended sequences following “Covariation analysis for experimentally validated transcriptional fluoride riboswitches”, except that the models were searched through all downloaded Rfam sequences to expand the motif analysis to all potential expression platforms.

### Oligo pool library amplification

The ssDNA oligo pool from Twist (see **Sequence acquisition and synthesis**) was resuspended to 20 ng/µL. A 50 µL PCR reaction was set up to amplify 20 ng of DNA: 1 µL of ssDNA pool (20 ng/µL), dNTPS (NEB, N0447L) to final concentration of 200 µM each, Q5 Reaction Buffer (NEB, B9027S) to a final concentration of 1x, Primer A and primer B (Supplementary Table 2) to a final concentration of 300 nM each, 0.5 µL of Q5 polymerase (NEB, M0491L), and water to a final volume of 50 µL. In a thermocycler (BIO RAD, S1000TM), the reaction was heated to 95 °C for 3:00 min and then cycled 15 times through 98 °C for 0:20 s, 53 °C for 0:15 s, and 72 °C for 0:15 s. The reaction was held at 72 °C for an additional 2:00 min to complete polymerase extension. The dsDNA product was purified using Sera-Mag Select Beads at a ratio of 1.5x (cytiva, 29343052), following the manufacturer protocol. This consisted of vortexing beads with the 50 µL PCR reaction, incubating at room temperature for 5 min, placing on a magnetic stand, removing supernatant, washing beads twice with freshly prepared 85% ethanol, air drying beads, and eluting dsDNA with 20 µL of water. The dsDNA concentration was quantified using a High Sensitivity Qubit assay (Invitrogen, Q32854). From this PCR reaction, 100 ng of dsDNA went through a second round of amplification following the same protocol to produce enough dsDNA for in vitro transcription. Input library quality was assessed by electrophoresis using the DNA 1000 assay on the Bioanalyzer (Agilent, 2100 bioanalyzer).

We conducted a PCR to prepare the oligo pool for next-generation sequencing to assess the sequence representation within the oligo pool. The PCR reaction was downscaled to 25 µL reaction: 0.5 µL of ssDNA pool (20 ng/µL), 0.75 µL of 10 mM dNTPS, 5 µL of 5x Q5 Reaction Buffer, 0.75 µL of 10 µM of each Primer C and primer D (Supplementary Table 2), 0.25 µL of Q5 polymerase, and water to a final volume of 25 µL. The reaction was then thermocycled as follows: 95 °C for 3:00 min, then cycled 2 times through 98 °C for 0:20 s, 54 °C for 0:15 s, 72 °C for 0:15 s), then cycled 12 times through 98 °C for 0:20 s, 65 °C for 0:15 s, 72 °C for 0:15 s, and then 72 °C for 1:00 min, then cooled to 12 °C. The reaction was purified using the using Cytiva Sera-Mag Select Beads at a ratio of 1.0x. Notably, the sequencing was performed with a 2×150 cycle kit (Supplemental_Document_A.xlsx, ‘Pool validation’ sheet).

### GreB protein purification

The GreB transcription elongation factor was purified following previously published methods with minor modifications (49). The expression plasmid was generously provided by Irina Artsimovich. Briefly, *E. coli* BL21(DE3) cells were transformed with the pIA577 plasmid encoding GreB with a C-terminal 6xHis tag (49). A single colony was grown in 1 L of auto-induction terrific broth media (Boca scientific GCM19.0500) supplemented with Kanamycin (50 µg/L). Cultures were incubated at 37 °C with shaking at 240 rpm until the OD600 reached ∼0.6, after which the temperature was reduced to 20 °C for overnight protein expression. The next morning, cells were harvested by centrifugation at 5,000 x g for 10 minutes at 4 °C, and the pellets were resuspended in 40 mL of lysis buffer containing 1 M NaCl, 40 mM Tris pH 7.5, 5 mM TCEP, 5% glycerol.

Cell lysis was performed via sonication on ice using a 12 mm tip placed inside the 50 mL Falcon tube with the cell pellet and conditions set at 50% amplitude, pulse cycle: 10 sec ON and 20 sec OFF for 5 cycles in total, and the lysate was clarified by centrifugation at 15,000 x g for 30 minutes. The supernatant was filtered through a 0.45 µm syringe filter and incubated with 5 mL of Ni-NTA agarose beads (Goldbio H-350-25) that were previously washed and pre-equilibrated with lysis buffer, for 1 hour at 4 °C with rotation mixing. The resin was loaded into a gravity column and washed with 50 mL of wash buffers containing each 20 mM and 50 mM imidazole to remove non-specifically bound proteins. The target proteins were eluted with 500 mM imidazole and fractions of 5 ml were collected.

Elution fractions containing the target protein with minimal impurities were identified by SDS-PAGE, pooled, and dialyzed overnight at 4 °C against dialysis buffer (2 M NaCl, 80 mM Tris pH 7.5, 2 mM TCEP, 5% glycerol). The protein was subsequently concentrated and stored at −20°C in storage buffer (1 M NaCl, 80 mM Tris pH 7.5, 2 mM TCEP, 50% glycerol). Protein identity and mass were confirmed by mass spectrometry. Importantly, the His tag was not removed in this protocol.

### Single- and multiple-round in vitro transcription with fluorescent output

To evaluate the effect of GreB in single- and multiple-round (runoff) *in vitro* transcription reactions, we adapted a standard transcription protocol to enable real-time fluorescence measurements following open complex formation. Reactions were prepared in conditions with or without 1.2 µM GreB and with or without 10 mg/µL Rifampicin (50)(Sigma-Aldrich R3501-250MG) to distinguish single- and multiple-round transcription, respectively. Briefly, the DNA template used for in vitro transcription consisted of the constitutive *E. coli* pJ23119 consensus promoter (from the registry of standardize biological parts) driving the expression of the 3WJdB fluorogenic RNA aptamer (51). The 3WJdB template was generated using PCR with Primers E and F (Supplementary Table 2), followed by a PCR cleanup (Qiagen, 28506) and concentrated to 1.3 µM stock. The dye DFHBI-1T (Tocris, 5610) was dissolved in DMSO to 40 mM. To allow open complex formation, 2.5 pmol of template DNA was incubated with 2 units of *E. coli* RNAP (NEB, M0551S) in 20 µL of buffer containing 62 mM KCl, 1.25 mM DTT, 25 mM Tris pH 8, 6.25 mM MgCl_2_, 0.125 mM EDTA, 0.21 mM DFHBI-1T, and 25 ng of BSA. Where indicated, 2 µL of 15 µM purified GreB was added to the reaction or water for noGreb. The reaction was then incubated at 37 °C for 10 minutes to allow open complex formation. Transcription was initiated by adding 2.5 µL of a start solution containing 5 mM NTPs. For single-round reactions, this buffer also contained 100 µg/µL Rifampicin to inhibit reinitiation (single round start solution). In multiple-round reactions, rifampicin was omitted and substituted with water. Two technical replicates of 5 µL from each reaction were transferred to a well of a 384 well plate (Corning, 3544) sealed with optically clear adhesive film (Thermo, 232701) and incubated at 37 °C in a plate reader (BioTek Synergy H1, H1M). Fluorescence intensity (Ex: 485 nm; Em: 520 nm; gain = 60) was measured every 3 minutes for 7 hours. Each experiment was performed in triplicate. Data is in Supplemental Document B.

To convert fluorescence arbitrary units (AU) to µM FITC equivalents, a standard calibration curve was generated using a NIST-traceable FITC standard solution (ThermoFisher, F36915), yielding a linear fit with R^2^ = 0.9999 and a slope of 10,446. This calibration was used to convert raw fluorescence data to µM FITC equivalents (Figure S1, Supplemental Document B).

## RNA-SEQ FOR NEXT-GENERATION SEQUENCING

The following steps were used for sequencing based assay: Single-round *in vitro* transcription of oligo pool, DNase digestion, 3’ Linker ligation, Reverse transcription, 3’ cDNA Adapter ligation, Sequencing library preparation, and gel purification (see Figure 2A).

**Figure 2.**
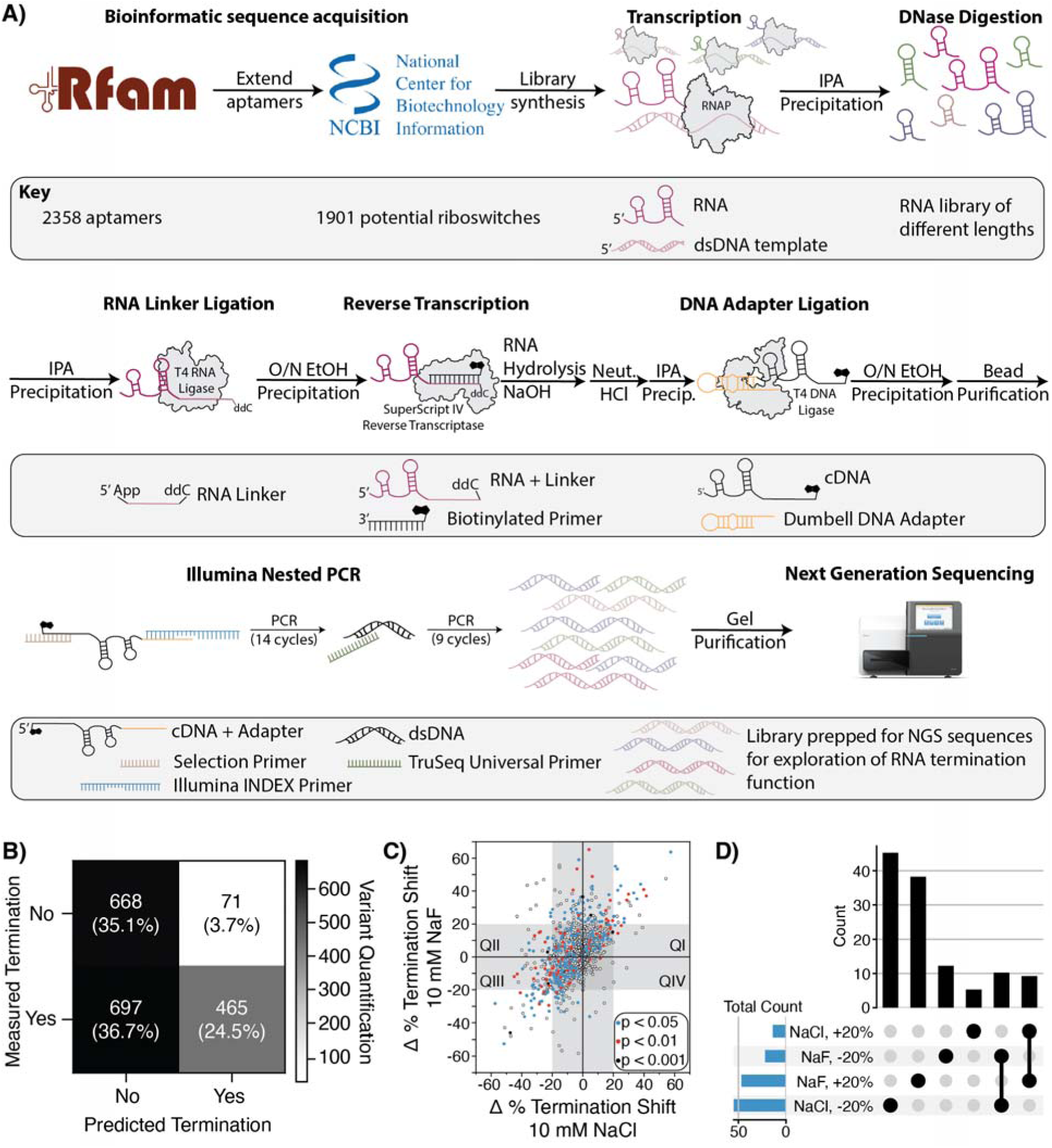
Experimental pipeline for high-throughput assessment of transcriptional riboswitch function. **(A)** Molecular schematic from acquiring aptamer sequences from Rfam to generating a Next Generation Sequencing library to characterize transcriptional read through. 2358 Fluoride aptamer (RF01734) sequences were collected from RFam and extended through NCBI genome records, resulting in 1901 riboswitch sequences. Promoter and priming site sequences were appended to create DNA templates, which were then synthesized and amplified. The DNA library was transcribed with the *E. coli* RNA Polymerase (PDB: 7YPA) in the presence of 10 mM NaCl or NaF and either 0 or 1.2 µM GreB. Cleanup was performed to isolate RNA constructs from proteins and DNA. To prepare the RNA library for next generation sequencing, the 3’ end was ligated (PDB: 1S68) with a known RNA sequence (light pink, Linker) to serve as the Reverse Transcriptase (PDB: 1MML) priming site. Then the 5’ end of the resulting cDNA (black) was ligated (PDB: 6DT1) with a known DNA sequence (orange, Dumbbell) and amplified using Illumina Primer sequences. The prepared DNA sequencing library underwent Next Generation Sequencing to identify reads as either terminated or anti-terminated, which were then counted for each variant and in each condition. **(B)** Confusion matrix tabulating which riboswitch sequences were predicted to be transcriptional regulators versus those measured to be transcriptional regulators through the NGS assay without GreB. The count and percent of variants for each category is written. The shading is for quantity comparison between the different conditions. **(C)** The difference of percent termination reads (1.2 µM – 0 µM GreB) for each variant in 10 mM NaCl (x-axis) and 10 mM NaF (y-axis). Variants in QI represent those that have increased OFF and ON signals in the presence of GreB, QII represent those that have reduced OFF and increased ON signals and thus higher dynamic range in the presence of GreB, QIII represent those that have reduced OFF and ON signals in the presence of GreB, QIV represent those that have increased OFF and decreased ON signals and thus lower dynamic range in the presence of GreB. A grey box encompasses −20%:20% for each salt condition to highlight the variants outside the box that have a change in termination by 20%. The Student’s Ttest was done for all comparisons with statistically significant variant changes being colored blue (p < 0.05), red (p < 0.01), and black (p < 0.001). **(D)** An UpSet plot to visualize the intersect of variants in the various conditions with possible outcomes of >|20%| change in termination rate with GreB that are statistically significant (p < 0.05) for NaCl or NaF. Total number of variants for each condition to the bar graph on the left (blue bars).

### Single-round *in vitro* transcription of oligo pool (high-throughput)

A 25 µL *in vitro* transcription reaction was prepared from the following: 2 µL of 1 M Tris pH 8.0, 0.02 µL of 0.5 M EDTA pH 8.0 (Invitrogen, 15575020), 1 µL of 0.1 M DTT (single-use frozen aliquot), 2.5 µL of 50 mM MgCl_2_ (NEB, B0510A), 0.25 µL of BSA (NEB, B9000S), 4.2 µL of 116 ng/µL amplified dsDNA template pool (final concentration of 100 nM), 2 µL of *E. coli* Holoenzyme (NEB, M0551S), either 2.5 µL of 100 mM of NaCl or NaF, and water (Invitrogen, 10977-015) up to 20.5 µL. Another 2 µL came from either water or GreB, which was first diluted in water to 15 µM and then added to the sample for a final transcription reaction concentration of 1.2 µM. The samples were incubated at 37 °C for 10 min. The transcription reaction was started through addition of 2.5 µL of single round start solution that included 5 mM NTPs for a final concentration of 500 µM (125 µM of each nucleotide) and Rifampicin (50) (Sigma-Aldrich R3501-250MG) to a final concentration of 0.01 mg/mL. Transcription was conducted for 5 min at 37 °C. The reaction was quenched by adding 75 µL of TRIzol (ambion, 15596018), followed by 20 µL of chloroform (Acros organics 423555000). The reactions were briefly vortexed and incubated at room temperature for 2 min. They were then centrifuged for 5 min at 4 °C at 12,000 g. The top 70 µL of the aqueous layer was placed into a new tube, followed by the addition 7 µL of 1 M NaCl, 1 µL of GlycoBlue (Invitrogen, ref. AM9515), and 50 µL isopropanol (Sigma-aldrich, 190764-1L). The reactions were inverted several times, incubated at room temperature for 10 min, and centrifuged for 10 min at 4 °C at 21130 g. The samples were then washed with 400 µL of 70% ethanol, centrifuged for 1 min at 4 °C at 21130 g, and air dried for 5 min.

### DNase digestion

Nucleic acid samples from single-round transcription of the oligo pool were resuspended in 50 µL containing: 43 µL of RNase-free water, 5 µL of 10x TURBO DNase buffer (invitrogen, 4022G), and 2 µL of TURBO DNase (invitrogen, AM2238), and incubated at 37 °C in a thermocycler for 60 min. The reaction was then quenched with 150 µL of TRIzol and 40 µL of chloroform, incubated at room temperature for 2 min, and centrifuged at 4 °C for 5 min. The top 140 µL of aqueous phase was extracted into a new tube. To this was added 14 µL of 1 M NaCl, 1 µL of GlycoBlue, and 100 µL of isopropanol, followed by inversion several times and incubation at room temperature for 10 min. This mixture was then centrifuged for 10 min at 4 °C at 21130 g. The samples were washed with 400 µL of 70% ethanol, centrifuged for 1 min at 4 °C at 21130 g, and air dried for 5 min. The RNA samples were resuspended in 9 µL of RNase-free water.

### 3’ Linker adenylation

A 3’ linker was then ligated to the post-DNase digested transcription products with the following procedure, which first adenylated the 5’ end of the linker. 5’ linker adenylation was performed with a reaction containing: 1.25 µL of 100 µM of oligo G (Supplementary Table 2), 2.5 µL 1 mM ATP (NEB, N0757A), 2.5 µL 10X NEB 5’ Adenylation Buffer (NEB, M2610S), and water to 22.5 µL. The reaction was scaled up to 100 µL and 22.5 µL was aliquoted into 0.2 mL PCR tubes. To each reaction, 2.5 µL of Mth RNA ligase (NEB, M2611AA) was added. Reactions were incubated at 65 °C for 1 hr. The reactions were combined and 300 µL of TRIzol was then added to the pooled reactions, which were then incubated at room temperature for 5 min. Eighty µL of chloroform was then added to the reactions, followed by incubation at room temperature for 2 min. The samples were then centrifuged at 21130 g for 15 min at 4 °C. The top 280 µL of the aqueous phase was extracted into a new tube. To this was added 1 µL of GlycoBlue, and 200 µL of isopropanol, followed by inversion several times and incubation at room temperature for 10 min. This mixture was then centrifuged for 10 min at 4 °C at 21130 g. The samples were washed with 400 µL of 70% ethanol, centrifuged for 1 min at 4 °C at 21130 g, and air dried for 5 min. The RNA linker pellet was dissolved in 20 µL of nuclease-free water and quantified by Qubit. The RNA molarity (MM: 6782.1 g/mol) was used to determine the final concentration for the linker ligation reaction.

### 3’ Linker ligation

The linker ligation was performed with a reaction containing: 6 µL of 50% PEG 8000 (NEB 1004S), 2 µL of 10x T4 RNA Ligase Buffer (NEB B0216S), and 2 µL of 200 nM 5’ App Linker prepared as above. The reaction mixture was vortex followed by the addition of 0.5 µL of SuperaseIn RNAse Inhibitor (Invitrogen, AM2696), 0.5 µL of T4 RNA Ligase, and 0.5 µL of Trunc KQ (NEB, M0373L) was added and mixed by pipetting up and down 10 times. A master mix according to the number of samples was made and then 11 µL of the master mix was distributed to each 9 µL sample. The final 20 µL reaction was incubated at 25 °C in a thermocycler for 2 hrs. Reactions were then ethanol precipitated by adding 130 µL of water, 15 µL of 3 M NaOAc (Invitrogen, AM9740), 1 µL of GlycoBlue, and 450 µL of ethanol. The samples were inverted several times and incubated at −20 °C overnight. The samples were then centrifuged for 30 min at 21130 g, washed with 70% ethanol, centrifuged for 1 min at 4 °C at 21130 g, and air dried for 5 min. The Linker-ligated RNA samples were then resuspended in 3 µL of RNase-free water.

### Reverse transcription

To perform reverse transcription, 3 µL of 0.1 µM Reverse Transcription (RT) primer H (Supplementary Table 2) was added to each linker-ligated sample and then incubated at 95 °C for 2 min, 65 °C for 5 min, and held at 25 °C to anneal the RT primer. During this time, an RT enzyme reaction was prepared containing: 7.5 µL water, 4 µL of 5x SSIV Buffer (invitrogen, 18090010), 1 µL of 10 mM dNTPs cNEB, N0447L), 1 µL of 0.1 M DTT (single-use frozen aliquot, invitrogen, 18090010), and 0.5 µL SSIV (invitrogen, 18090010). A master mix was made according to the number of samples. After the samples had been at 25 °C for at least 1 min, 14 µL of the master mix was added and mixed by pipetting. The complete reaction was incubated at 25 °C for 2 min, 50 °C for 10 min, 80 °C for 10 min, and held at 4 °C. After the reaction had been at 4 °C for at least 1 min, 1 µL of 4 M NaOH was added to hydrolyze the RNA, mixed by pipetting, briefly vortexed and spun down. The reactions were then heated at 95 °C for 5 min and then cooled to 4 °C. Two µL of 1 M HCl were added to each reaction to neutralize pH, followed by 17 µL of isopropanol. The samples were then incubated at room temperature for 10 min, centrifuged at 4 °C for 10 min at 21130 g, washed with 70% ethanol, centrifuged for 1 min at 4 °C at 21130 g, and airdried for 5 min. The resulting cDNA samples were resuspended in 7 µL of RNase-free water.

### 3’ cDNA Adapter ligation

Each cDNA reaction was then ligated on the 3’ end to a Dubmbell adapter following the StructureSeq 2.0 protocol (52). To each cDNA reaction was added 0.5 µL of 100 µM of SS2.0 Dumbbell (Oligo I, Supplementary Table 2). Samples were incubated on a thermocycler at 95 °C for 2 min and then cooled to 21 °C for 3 min. During this time, a ligation mix was prepared containing: 2.5 µL 10x T4 DNA Ligase Buffer (NEB B0202S), 2.5 µL of 5 M Betaine (Sigma, B0300-1Vl), 10 µL of 50% PEG8000 (NEB, B1004S), and 2.5 µL T4 DNA Ligase (NEB, B0202S). The reaction mix was scaled up according to the number of samples and then aliquoted to each cDNA sample. Reactions were then incubated at 30 °C for two hours, at 65 °C for 15 min to deactivate the enzyme, and cooled to 4 °C. To each reaction, 125 µL of RNase-free water, 15 µL of 3 M NaOAc, 1 µL GlycoBlue, and 450 µL of cold ethanol were added. The samples were left at −20 °C overnight. Samples were spun down for 30 min at 21130 g for 30 min at 4 °C, washed with 70% ethanol, centrifuged for 1 min at 4 °C at 21130 g, and airdried for 5 min. The ligated cDNA was then resuspended in 50 µL of nuclease free water. To remove excess Dumbbell and RT primer, bead purification was done with Agencourt AmpureXP (Beckman Coulter, A63881) at 1.8x ratio according to the manufacture’s protocol. The samples were resuspended in 20 µL of nuclease of free water and stored in −20 °C.

### Sequencing library preparation

Sequencing libraries were prepared through PCR reactions on the ligated cDNA to complete adapter sequences and add necessary indexes. PCR reactions were assembled by first combining 5 µL of each ligated cDNA sample, 0.25 µL of 100 µM Index primer (Oligo J, Supplementary Table 2, 3) in a PCR tube. To add sequence variability for the start of the sequencing reads, two different selection primers were used, primer K for NaCl conditions and primer L for NaF conditions (Supplementary Table 2). For each selection primer a master mix was made consisting of 21 µL of nuclease free water, 10 µL of 5X Q5 Rxn Buffer, 12.5 µL of 100 nM Primer K or L, 0.5 µL of 10 mM dNTPs, and 0.5 µL Q5 Polymerase. The master mix was aliquoted in 44.5 µL to the respective tube, i.e. primer K to NaCl conditions and primer L to NaF conditions. The following thermocycler protocol was then run: 98 °C for 0:30 s, then cycled 14 times through 98 °C for 0:10 s, 65 °C for 0:30 s, 72 °C for 0:30 s, 12 °C forever. At this point, 0.25 µL of 100 µM Illumina TruSeq Universal Adapter (Oligo M, Supplementary Table 2) was added to each reaction. The PCR reaction continued with 9 cycles through 98 °C for 0:10 s, 65 °C for 0:30 s, 72 °C for 0:30 s, and then 72 °C for 5:00 min, 4 °C for 3 min. At this point, 0.25 µL of ExoI (NEB, M0293S) was added to each reaction to remove excess primer. The reactions were then incubated at 37 °C for 30 min, followed by a 20 min enzyme deactivation step at 80 °C, and cooling to 4 °C. The samples were then transferred to a 1.7 mL tube for PCR clean up using 5x Buffer PB (Qiagen, 19066), applied to a silica spin column (NEB, T1017L), vacuumed, washed twice with 750 µL of Buffer PE (Qiagen, 19065), vacuumed, placed back into a collection column and dry spun for 1 min at 21130 g. The reaction was eluted in 15 µL of water and spun for 1 min at 21130 g.

### Gel purification of sequencing library

Sequencing libraries were loaded into an 8% denaturing gel (national diagnostics, EC-830, EC-835, EC-840) and run at 18 W for 2 hrs. The gel was stained with 3 µL of SybrGold (invitrogen, S11494) for 5 min in 0.5x TBE buffer (RPI, T60040-1000; Sigma, B7901; invitrogen, 15575020) while rolling and visualized with the SmartBlueTM Transillluminator (Accuris, E4000). Each sequencing library (between 200 – 404 bp) was then cut out and eluted overnight in 1 mL of 1x Buffer TE (IDT, 11-05-01-13) rotating at 4 °C in a 2 mL tube. Due to tube size limitation, the 1 mL elution reactions were split into 500 µL into 1.7 mL tubes for ethanol precipitation. To each tube 60 µL of 3 M NaOAc, 1 µL GlycoBlue, and 1250 µL of cold ethanol was added. Reactions were incubated at −70 °C for 30 min and centrifuged at 21130 g for 30 min at 4 °C, washed in 70% ethanol, centrifuged for 1 min at 4 °C at 21130 g, and air dried for 5 min. The sequencing libraries were resuspended in 11 µL of nuclease-free water and concentrations measured using the High Sensitivity Qubit assay (invitrogen, Q32854).

### Next Generation Sequencing

To assess the success of the gel purification prior to NGS, the purified libraries were run on an 8% denaturing gel for 2 hrs at 18 W (Figure S2A). The libraries were sent to NUSeq Core Facility for Paired End, 150-cycle sequencing on the Element AVITI (System: 880-00001). The core performed additional quality control with a 2100 bioanalyzer (Figure S2B) and KAPA qPCR quantification prior to loading.

### Read analysis

FASTQ Paired End files were combined using PEAR (v0.9.6) (53) using the following command line: pear -f [library_#_R1].fastq.gz -r [library_#_R2].fastq.gz -o library_#/ library_#_PEAR -y 4G -j 1, where the # symbol refers to the sample number (Supplemental Document A). The reads were aligned to their accession number and genome positions using STAR (v2.7.9a) (54). A synthetic genome was made from the pool of sequences (Supplemental Document A, ‘Bioinformatics’ sheet) by removing the J23119 promoter sequence and formatting the sequences into a FASTA format with their accession IDs. The FASTA file (see **Data and Code Availability**) was turned into a genome assembly using megahit (v1.0.6.1) and calibrated with the genomeGenerate mode from STAR.

To align the reads to the reference genome, STAR was run with the following specified parameters: --alignIntronMax 1 --alignEndsType Local --alignSoftClipAtReferenceEnds Yes --outFilterScoreMinOverLread 0.5 --outFilterMatchNminOverLread 0.5 --outSAMmultNmax 1 (Figure S3). The generated output SAM file was used to quantify reads through the Read_Classification_SUBMIT.py script on a Jupyter Notebook running python (v. 3.10.9). Reads were filtered out if the start position of the alignment was not within the beginning of the riboswitch sequence. The length of the transcript was calculated and polyU sites were searched using the regular expression: “’TTTTT|T[AGC]TTTT|TT[AGC]TTT|TTT[AGC]TT’”. Reads were classified as “terminated” if they ended from 2 nts before the pattern through 6 nts after the pattern. Results from this analysis are within the file Supplemental Document A, ‘Rep1_output’ and ‘Rep2_output’ sheets. The Pearson correlation coefficient (r) was used to assess the read count similarity between technical replicates (Supplementary Table 5) in Excel using the command: =PEARSON([array1],[array2]).

Data plotting was done with DataGraph (v4.5.1) and python 3 (v. 3. 10.9, confusion_matrix, ConfusionMatrixDisplay, matplotlib).

### Calculation of percent Anti-Termination (%AT)

Percent anti-termination (%AT) was calculated by the formula 1-“terminated” reads/total reads (Supplemental Document A, Figure S4). The %AT were averaged between the two replicates and the change (Δ) of %AT between conditions was calculated (%AT NaF - %AT NaCl). Then the reads were classified as "Transcriptional" or "Translational", where variants were marked as "Translational" if their %AT was greater than 95%. If "Transcriptional" the variants are further categorized into "Off" if Δ%AT < −20%, "Non-functional" if −20% < Δ%AT < 20%, "Low" if 20% < Δ%AT < 40%, "Mid" if 40% < Δ%AT < 50%, and "High" if 50% < Δ%AT. Then, the Δ%AT change between 0 and 1.2 µM of GreB was assessed. If GreB altered the Δ%AT by at least 20%, then GreB was noted as impacting riboswitch function.

### Single dsDNA template preparation for individual in vitro transcription

Individual riboswitch sequences were ordered as gBlocks (IDT) that matched their sequence in the oligo pool and resuspended to 10 ng/µL. A 50 µL PCR mix was made of: 5 µL 10X Taq Buffer (NEB, B9014S), 1 µL 10 mM dNTP mix (NEB, N0447L), 0.25 µL of 100 µM Primer A (Supplementary Table 2), 0.25 µL of 100 µM Primer B (Supplementary Table 2), 0.25 µL of 10 ng/µL gBlock, and 0.25 µL Taq DNA Polymerase (NEB, M0273X), and water to 50 µL. The reaction was then heated to 95 °C for 3 min, then cycled 25 times through 95 °C for 0:45 s, 53 °C for 0:30, and 68 °C for 1 min. A final 68 °C extension occurred for 5 min and then the reaction was cooled to 12 °C. The PCR reaction was cleaned up by adding 5x Buffer PB (Qiagen, 19066) and applying the solution to a spin column (Epoch life science, 1910-250). The sample was washed twice with 750 µL of Buffer PE (Qiagen, 19065), spun for 1:00 min to remove excess liquid, and eluted in 40 µL of RNase-free water (invitrogen, 10977-015). DNA template concentrations were measured using the Broad Rang Qubit assay (invitrogen, Q32853).

### Single-round *in vitro* transcription of single riboswitches (low-throughput)

To validate the next-generation sequencing results, the single-round *in vitro* transcription with GreB was repeated with dsDNA templates of individual sequences (Supplemental Document A, ‘Low-Throughput Validation’ sheet). After DNase digestion, the samples were resuspended in 10 µL of RNase-free water. 10 µL of 2x loading buffer (8 M urea with bromophenol blue dye (Sigma-Aldrich, B0126-25G) and run on a 10% denaturing gel (national diagnostics, EC-830, EC-835, EC-840) at 18 W for 90 min with 0.5 µL of 1% ssRNA Century Plus Ladder (invitrogen, AM7145). The gel was stained for 5 min with 3 µL of SybrGold gel (invitrogen, S11494) in 40 mL 0.5x TBE gel (RPI, T60040-1000; Sigma, B7901; invitrogen, 15575020) buffer while rolling. The gel was imaged on the ChemiDocTM Touch Imaging System (BIO RAD, 1708370). Gel analysis was done using Image Lab (v6.1) to assess band intensity. Percent anti-termination (% AT) was determined by subtracting the annotated terminated band from 100%.

## RESULTS

### A bioinformatic pipeline to discover transcriptional fluoride riboswitch variants

To begin this study, we first developed a bioinformatic pipeline to find fluoride riboswitch variants that are predicted to regulate at the transcriptional level. Most bioinformatic analysis performed on riboswitch sequences is done on the highly conserved aptamer domains, which perform a single function of binding a specific ligand. RNA structure databases, such as Rfam (45), only contain information on the aptamer sequence and structure and do not contain information about the downstream expression platforms (EPs), which can vary in function, sequence and structure. To analyze transcriptional expression platforms of riboswitch variants, we first created a bioinformatic pipeline to collect genomic sequences downstream of the aptamers that could contain expression platforms. We then computationally predicted whether these expression platforms could contain an intrinsic terminator.

Our pipeline consisted of several steps (Figure 1C, Methods and Materials): (1) collect aptamer sequences from the fluoride riboswitch Rfam family (RF01734); (2) extend aptamer sequences to include the downstream EP sequence using aptamer genomic coordinates; and (3) computationally predict whether the extended EPs could contain intrinsic terminators. Aptamer sequences were acquired from a representative pool of fluoride aptamers from the RNA structure family database, Rfam (45), which results in a collection of 2358 aptamer sequences (Figure 1C). For each aptamer, we next extended its sequence up to a length that fit within the 300 nt limit for DNA pool synthesis. This was done using the NCBI accession ID and aptamer genomic coordinates within the Rfam FASTA file of aptamer sequences. After extending the sequences, we were left with a collection of 1901 potential riboswitch variants since non-NCBI entries meant there was no genomic context from which to acquire downstream sequence (e.g. URS0000BC4686_32630 from Patel, et. al.(15))(Figure 1C). To check that the extended sequences capture all the downstream EP, we bioinformatically searched for the location of the start codon of the first open reading frame (ORF) downstream of each aptamer (Figure S5). This analysis showed that 1566/1901 (82%) riboswitch sequences had start codons within the DNA pool synthesis limits, suggesting that our pipeline captures the complete expression platform for most sequences.

To identify which riboswitch variants potentially regulate gene expression using intrinsic termination, we next analyzed extended EP sequences with the ARNold webserver (46). The ARNold webserver predicts intrinsic termination through two algorithms: Erpin, which is trained on terminator sequences from *Bacillus subtilis* and *E. coli* (55), and RNAmotif, which is a descriptor with a score cutoff to identify terminator hairpins (56). The ARNold webserver predicted 536 (28%) of the fluoride aptamers in the pool were coupled to an expression platform that specifically folds into an intrinsic terminator (Supplemental Document A). Of these, 128 terminators were predicted by both algorithms, 207 predicted by Erpin, and 201 predicted by RNAmotif. These terminators had a predicted free energy range from −17.1 to −2.5 kcal/mol, with an average of −10.17 kcal/mol (Figure 1D), which falls within expected values (57,58). Furthermore, the full riboswitch sequences, including aptamer and extended EP, ranged between 49 – 208 nts in length (98 nt average) (Figure S6A). The terminator start positions range from within the aptamer to over 100 nts away, with an average terminator start within the aptamer (∼57 nts) (Figure S6B, Supplemental_Document_A.xlsx). Predicted total terminator lengths were between 19 – 53 nts (36 nt average) (Figure S6C), with hairpin stems from 6 – 10 base pairs (10 nts average) (Figure S6D) and loop lengths from 1 – 32 nts (15 nts average) (Figure S6E). While ARNold did not identify terminators for the majority of riboswitch variants, our pipeline successfully identified a large pool of transcriptional fluoride riboswitch candidates that we experimentally assessed for regulatory function.

### A high-throughput next-generation sequencing pipeline to measure intrinsic termination of 1901 riboswitch variants

To experimentally assess the function of each fluoride riboswitch variant, we converted the in silico generated riboswitch sequences into a pool of DNA transcription templates using massively parallel oligonucleotide synthesis (41). To each riboswitch sequence, we added a 60 nt sequence to the 5’ end that contains the J23119 consensus *E. coli* promoter sequence. We also added a 20 nt 3’ primer binding site for oligo pool amplification. After synthesis, we sequenced the oligo pool to characterize sequence variant coverage, which showed that each member was represented in the pool, with several sequences overrepresented likely due to their shorter length (Figure S7, Supplemental Document A).

We next amplified the oligo pool into a dsDNA template library and performed *in vitro* transcription in either 10 mM NaCl or 10 mM NaF. The resulting RNA pool was then processed in a series of preparation steps for next-generation sequencing (Figure 2A). Sequencing reads were processed by aligning them to riboswitch variant IDs, mapping the end of the sequencing read to the genomic position, and using this genomic position to classify the read as either terminated or anti-terminated (Figure S8, Methods). If a polyU was identified, then the reads that ended within that region were classified as terminated (Methods). A plot of the normalized read counts for each variant across replicates showed strong correlations with R^2^ of 0.97 (10 mM NaCl) and 0.99 (10 mM NaF) (Figure S9A, S9B). We found 1138 variants to have terminated classified reads, with 1111 overlapping between the two replicates (Figure S10A).

Given recent work highlighting the dynamic relationship between transcription factors and riboswitches (39,43,44), we decided to test a previously unexplored transcription elongation factor to identify novel crosstalk between riboswitches and transcription. We supplemented the in vitro transcription reaction with 1.2 µM of the transcription elongation factor, GreB, which rescues stalled RNAPs that have backtracked by cleaving the 3’ end of the RNA transcript in the RNAP active site (42). To confirm that the addition of GreB alters our transcription reaction, we performed IVT of a three-way junction dimeric broccoli construct (51), which showed that GreB increases transcriptional output (Figure S11). We next repeated transcription, processing, sequencing and analysis of the DNA template pool with 1.2 µM GreB. Again, we found replicate experiments with GreB highly reproducible with R^2^ of 0.98 (10 mM NaCl) and 0.98 (10 mM NaF) (Figure S9C, S9D).

To identify intrinsic termination events, we wrote a Python script that classifies reads as “terminated” if they ended at a polyU site, and “anti-terminated” if they ended beyond the identified polyU site for that variant (Methods and Materials). If no polyU site was identified in the variant, then all reads are classified as “Anti-Terminated”. To reduce noise, the number of “Terminated” reads had to be over 5%.

After this analysis, 1165 variants were identified as transcriptional with 1111 variants classified as “Terminated” in both technical replicates (Figure S10A). For the replicates with the transcription factor, GreB, there were 1162 variants identified with “Terminated Reads” with 1049 variants shared between replicates (Figure S10B). A confusion matrix comparing the average number of variants with measured termination without GreB (1162 variants after averaging the replicates) versus those predicted to contain terminators by ARNold showed that 59.6% of predicted terminators were correctly identified while 40.4% were incorrectly identified (Figure 2B). The major failure mode of the prediction was the 36.7% of variants predicted to not terminate while they were measured to undergo some termination (Figure 2B). We used RNAStructure (59) to assess terminator structures of a random selection of riboswitches that were measured to terminate. Those variants that were successfully predicted by ARNold had a perfect hairpin whereas the unsuccessful predicted variants had a mismatch within the terminator hairpin stem (Figure S12). When ARNold successfully predicted a terminator, we found it accurately predicted the complete riboswitch (R^2^ = 0.95) (Figure S13).

For terminating variants, riboswitch lengths were calculated by taking the median read length within the polyU site (Supplemental Document A, ‘Termination Analysis sheet’). The shortest variant is 46 nts, the longest is 219 nts, and the median terminating riboswitch length is 90 nts (Figure S14A), which is similar to the range and average predicted by ARNold (Figure S6A). We then subtracted the aptamer length from the riboswitch length to approximate the terminator lengths. The shortest terminator was calculated to be −94 nts, with the negative number due to extremely long aptamers sequences deposited on Rfam, e.g. MGTH01000159.1/17659-17500 with a deposited aptamer length of 161 nts. The longest terminator length was calculated to be 157 nts, and the median terminator length to be 31 nts (Figure S14B). The median terminator length is similar to the ARNold prediction while the shortest and longest lengths vary greatly (Figure S6B). The majority of riboswitches had downstream start codons after the identified polyU site, with a median position of 40 nts (Figure S14C), further supporting that these variants are putative riboswitches. Together, these demonstrate the large range of terminating variants our assay can detect.

As GreB acts on arrested RNAP complexes (42), we sought to assess how transcription termination frequency changed between the without (0 µM) and with (1.2 µM) GreB reactions (Supplemental Document A, ‘Termination Analysis sheet’). When plotting the change in termination, highlighting changes greater than 20%, between the NaCl or NaF conditions, we observe that variants predominately cluster as either overall increased signal or decreased signal (QI and QIII in Figure 2C), suggesting an overall positive correlation that is indifferent to salt composition. We also tested for significant changes using Student’s t-test (see Material and Methods) and counted which variants had a p-value < 0.05 and greater than 20% change in termination (Figure 2D). Notably, only 19 variants significantly change in both conditions.

Overall, our NGS pipeline was able to identify a larger number of transcriptional fluoride riboswitch variants than the number predicted by ARNold that we could then validate for regulatory function.

### Comparison of high-throughput NGS characterization of riboswitch function with IVT

We next sought to validate our NGS pipeline by comparing the quantification of percent anti-termination (%AT) with and without NaF with that quantified by low-throughput *in vitro* transcription (IVT) experiments. For a particular riboswitch variant in the NGS assay, we calculated a %AT value by classifying sequence reads as “Terminated” or “Anti-Terminated” based on the read alignment end position, and calculated %AT by dividing the number of terminated reads by the sum of all the categorized reads and subtracting from 1 in both NaCl and NaF conditions (Figure 3A). This NGS quantification was then compared to IVT experiments by running IVT outputs on a denaturing PAGE gel, classifying bands as “Terminated” and “Anti-Terminated” based on length, and calculating %AT based on band intensities (Figure 3B). We then calculated a change in %AT (Δ %AT) by subtracting the +NaCl condition from the +NaF condition.

**Figure 3.**
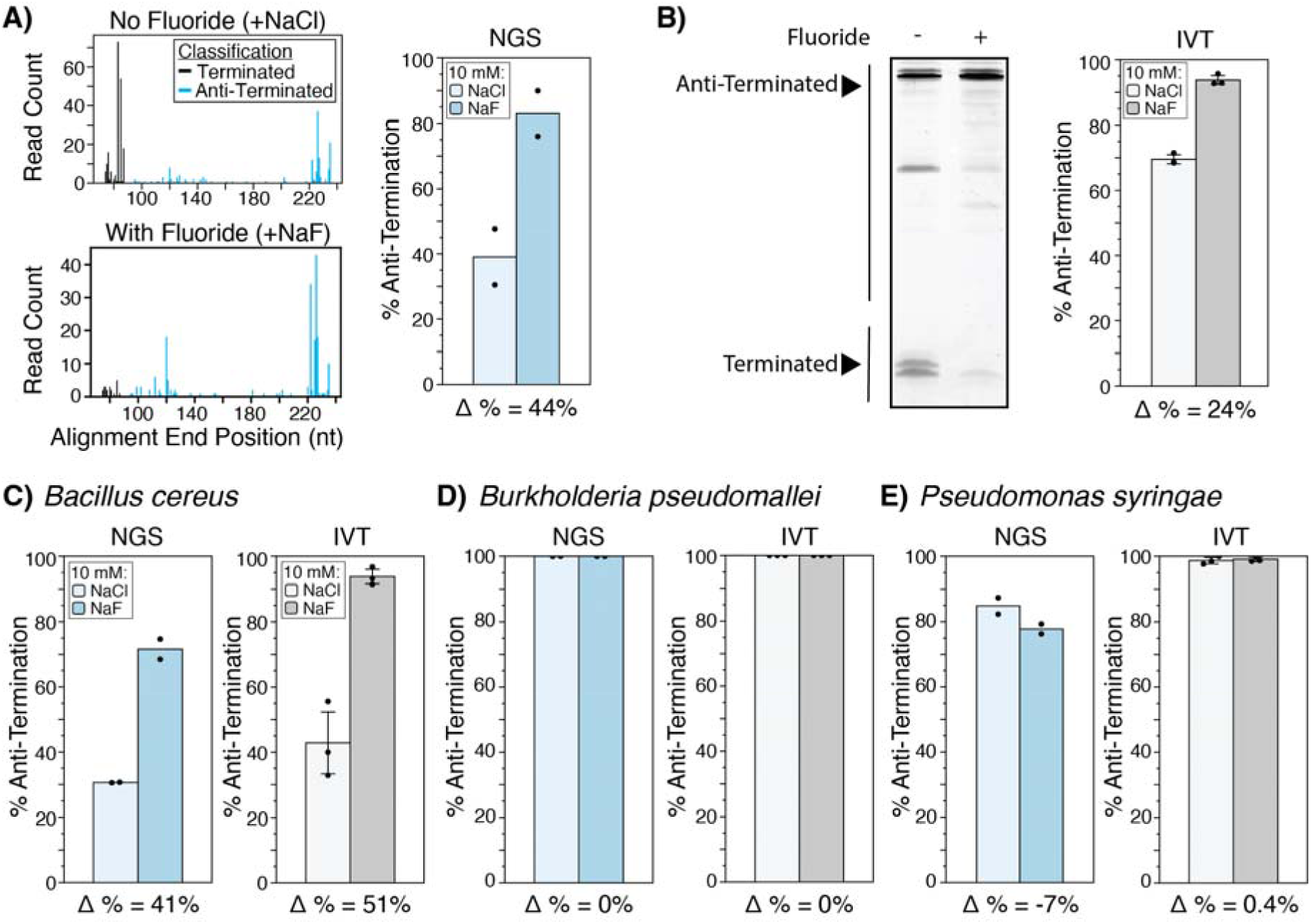
Validation high-throughput versus IVT characterization of published fluoride riboswitch function. **(A)** NGS analysis of the *Thermotoga petrophila* (accession ID: CP000702.1/1794817-1794880) riboswitch. Plots represent read alignments classified as terminated or anti-terminated in 10 mM NaCl (left, top) or NaF (left, bottom) in the presence of 1.2 µM GreB. The percent Anti-Termination was calculated as 1 - (Terminated count)/(Anti-Terminated count + Terminated count) and plotted in a bar graph with two replicates (right). The change (Δ) of percent (%) Anti-Termination is annotated on the plot. **(B)** IVT quantification (with 1.2 µM GreB) of the *Thermotoga petrophila* variant. The percent Anti-Termination was calculated by quantifying the band intensities of the regions annotated as Anti-Terminated and Terminated (left) using the formula from (A) and plotted in a bar graph (right). **(C, D, E)** NGS- and IVT-quantified percent Anti-Termination for **(C)** *Bacillus cereus* (accession ID: CP000227.1/4763720-4763779), a transcriptional control, **(D)** *Burkholderia pseudomallei* (accession ID: BX571966.1/2539005-2538939), a translational control, and **(E)** *Pseudomonas syringae* (accession ID: AE016853.1/5215709-5215637), a translational control. Bars represent average % anti-termination with error bars representing ± standard deviation (N = 2 for the high-throughput NGS assay, N = 3 for the low-throughput IVT assay). Structures for aptamer and expression platforms and high-throughput data without GreB are in Figure S15. Annotated gel images: Figure S19.

We used this method to validate the NGS approach using two transcriptional fluoride riboswitch variants (60), *Thermotoga petrophila* (CP000702.1/1794817-1794880) (*T. pe*, Figure 3A,B, S15A) and *Bacillius cereus* (CP000227.1/4763720-4763779) (*B. ce*, Figure 3C, S15B). For *T. pe* we observed a Δ %AT of 44% in the NGS assay but 24% in the IVT assay. A closer analysis revealed that this difference was mostly due to a lower %AT value in +NaCl condition in the NGS (39%) vs. the IVT (70%) assay. The comparison was much closer for *B. ce*, where we observed a Δ %AT of 41% in the NGS assay and a 51% in the IVT assay, with overall higher %AT in both NaCl and NaF conditions in the IVT assays.

We next validated our approach using two translational fluoride riboswitch variants (61), *Burkholderia pseudomallei* (BX571966.1/2539005-2538939) (*B. ps*, Figure 3D, S15C) and *Pseudomonas syringae* (AE016853.1/5215709-5215637) (*P. sy*, Figure 3E, S15D). As expected for *B. ps* we observed a Δ %AT of 0% in both the NGS and IVT assays. For *P. sy*, we observed a Δ %AT of −7% (85% AT in the NaCl condition and 78% in NaF condition) in the NGS assay but 1% (90% AT in the NaCl condition and 100% in NaF condition) in the IVT assay.

Overall these results show that the NGS assay is able to accurately classify riboswitches as using intrinsic termination, though quantitative values of %AT differ when compared to more traditional IVT assays.

### Discovering highly functional transcriptional riboswitches with NGS

We next sought to evaluate the function of the identified transcriptional fluoride riboswitches, i.e. the difference of Anti-Termination (AT) between the +NaCl and +NaF conditions. For each variant, we calculated the change of %AT (“Δ%AT”) by subtracting the %AT in the NaCl condition from the %AT in the NaF condition. We then used the calculated Δ%AT to classify each variant in both 0 µM and 1.2 µM GreB conditions as “Off” (−40 to −20%), “Non-functional” (−20%-20%), “Low functional” (20-40%), “Mid functional” (40-50%), and “High functional” (50-100%). Using this classification, we identified several variants in each category with a majority of riboswitch variants in the “Non-functional” category (Figure 4A), which could be due to incompatibilities with riboswitch folding or terminator function and the *E. coli* RNAP used in this experiment.

**Figure 4.**
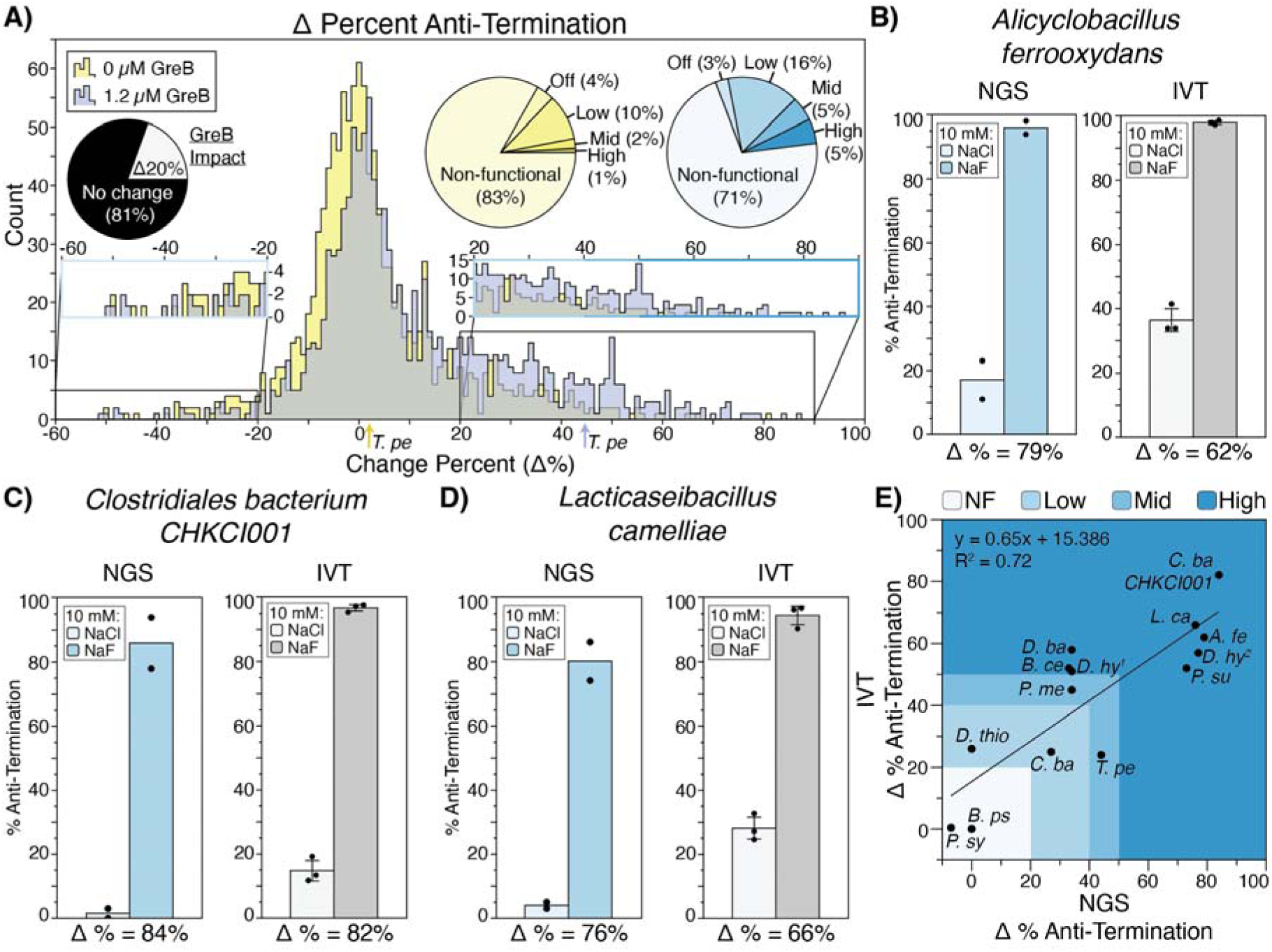
Estimating fluoride riboswitch function from high throughput sequencing assay. **(A)** Distribution of change (Δ) in the percent anti-termination measured by the NGS assay for variants with detected intrinsic termination. Distributions are shown from assays without GreB (yellow), and with 1.2 µM GreB (blue) (overlap = grey). The dynamic range in response to fluoride, Δ%AT (change in percent anti-termination), was determined by averaging the % Anti-Termination between each condition replicate and then calculating NaF % Anti-Termination - NaCl % Anti-Termination. Variants were classified according to calculated Δ ranges: “Off” (−40 to −20%), “Non-functional” (−20%-20%), “Low functional” (20-40%), “Mid functional” (40-50%), and “High functional” (50-100%). Pie charts show classification percentages for each GreB condition. Arrows labeled “*T. pe*” designate the *Thermotoga petrophila* switching function in 0 µM (blue) and 1.2 µM (yellow) GreB. **(B, C, D)** IVT validation of NGS assay results for *Alicyclobacillus ferrooxydans* (*A. fe*; LJCO01000051.1/58159-58221, **(B)**), *Clostridiales bacterium CHKCI001* (*C. ba CHKCI001*; FCNS01000019.1/7032-7129, **(C)**), and *Lacticaseibacillus camelliae* (*L. ca*, AYZJ01000062.1/6126-6187, **(D)**). Data shown on the blue bar graphs are from the two biological replicates (N = 2) of the high-throughput sequencing assay with 1.2 µM of GreB. Data shown on the grey bar graphs are from three (N = 3) replicates of single-round *in vitro* transcription reactions with 1.2 µM GreB. Both assays are done in the presence of either 10 mM NaCl or NaF. Data is presented as percentage anti-termination with ± standard deviation. Under each graph is the change (Δ) of percent (%) Anti-Termination between the NaF and NaCl conditions for the dataset (Supplemental Document A). Structures for aptamer and expression platforms and high-throughput data without GreB are in Figure S16. **(E)** Correlation of the average Δ % Anti-Termination in the NGS (x-axis) vs IVT (y-axis) measurements. Points are plotted for 10 riboswitches and labeled corresponding to their species (Supplementary Table 6, Figures S15-17). Colors show riboswitch functional classification. The line of best fit is drawn with a slope of 0.65 and a y-intercept of 15.386% and an extracted R^2^ = 0.72. Graphing calculations done in the DataGraph (v4.5.1) software. Gel imagines are shown in Figure S18-S21.

As we observed that GreB impacts transcription termination in this assay (Figure 2C, 2D), we next compared how the addition of GreB affects riboswitch function in this assay. Interestingly, when GreB was present, 24% of variants had an increase in Δ %AT of at least 20% (Figure 4A), suggesting GreB is important for this subset of variants. For example, the *T. pe* riboswitch Δ %AT changed from 1% without GreB to 44% with GreB (Figure 4A, Figure S15), which mostly comes from a decrease in the %AT in the NaCl condition (Figure S15). In aggregate, 14% of riboswitch variants went from “non-functional” or “off” to functional, and 13% improved their function, showing that GreB has an impact on riboswitch function.

We next sought to establish how individual variants characterized in the high-throughput NGS assay perform when individually isolated in the low-throughput IVT assay. We selected several riboswitch variants in each functional category, and not currently characterized in the literature to our knowledge (Figure 4B-D, Figure S15-S17), as well as several variants with a greater dynamic range than the current fluoride riboswitch model variant, *B. ce* (Figure 3C) (4,37–39,60,62). Correlating the Δ %AT values measured by the NGS and IVT assays (Figure 4E), shows a strong correlation (R^2^ = 0.72), but with a slope of 0.65, showing that the NGS assay correctly categorizes each variant, but overestimates the quantitative value of Δ %AT compared to IVT. This could be due to a function of the different assay conditions in NGS, with all riboswitches being transcribed simultaneously, or due to steps in the NGS library preparation that could introduce bias, for example by favoring shorter RNA fragments during amplification and flow cell clustering (63)(Illumina Knowledge Article #3874).

Ultimately, these results demonstrate that the high-throughput NGS assay can accurately categorize nearly two-thousand riboswitch variants into functional categories in a single experiment, and that it can be used to investigate how transcription conditions (i.e. the addition of GreB) affects the performance of these systems. This leads to rapid identification of highly functional switches, such as the *Clostridiales bacterium* fluoride riboswitch which we found to have Δ %AT = 84%, which is the highest we have observed.

### Using experimentally-guided bioinformatics to develop a covariation motif for a complete transcriptional riboswitch

Covariation has been instrumental to riboswitch discovery (20,26) as aptamers have highly conserved structures. Covariation models work by taking an input sequence alignment and searching for positions that co-vary (i.e. an A position changing to a G, and a downstream U position changing to a C), indicating that these changes occur during evolution to preserve base pairing (23,24). However, covariation analysis has been difficult to apply to riboswitch expression platforms given their wide sequence and functional variability (20). To address this limitation, we developed an approach to filter expression platforms that function as transcription terminators through either computational (Figure 1) or experimental (Figure 2) means. We hypothesized that this functional filtering would constrain the expression platform sequences enough to be able to develop a covariation motif for the entire transcriptional fluoride riboswitch sequence including both the aptamer and terminator.

From the functionally filtered sequence, we perform two alignments: the aptamer portion using the nhmmer command in R-scape (RNA Structural Covariation Above Phylogenetic Expectation) (24,47) and the sequence after the aptamer to the polyU termination site using MUSCLE (48). We then devised the Shelley method to produce “Frankenstein” alignments for the aptamer and terminator to create a combined aptamer-terminator alignment. Covariation was assessed using R-scape (24) (RNA Structural Covariation Above Phylogenetic Expectation) (Figure 5A). The holo structure, when ligand is bound, was the automatic output of CaCoFold; whereas the apo structure, when ligand is unbound, was drawn with the alternative covarying pattern (Figure S22).

**Figure 5.**
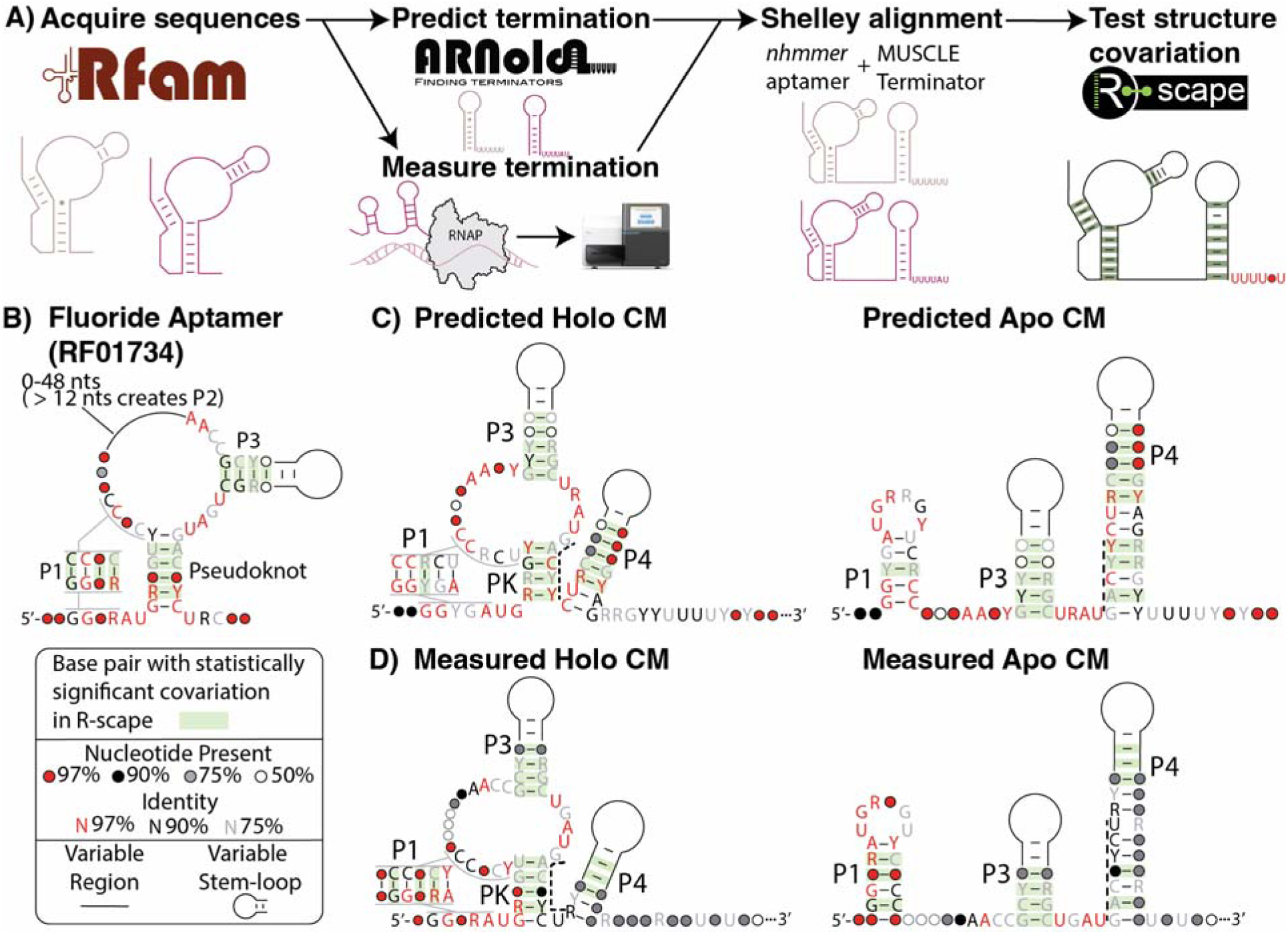
Deriving an experimentally-informed covariation model of the fluoride riboswitch aptamer and expression platform. **(A)** Computational pipeline to establish a covariation model. Fluoride aptamer sequences were acquired from Rfam family RF01734, and extended by approximately 150 nts (Figure 1C). Expression platform sequences were filtered to preserve those that contain an intrinsic terminator using either computational prediction by ARNold (Figure 1), or through experimental validation (Figure 2). A Shelly alignment was used to create an alignment for the aptamer and the terminator, which was then used in R-scape to generate a covariation model. **(B)** Covariation motif of the fluoride aptamer from Rfam (RF01734). Green highlighting on base pairs denotes evolutionarily significance in base pair covariation, and nucleotides are denoted as circle or specific letters with colors to signify conservation: red = > 97%, black = 90-97%, grey = 75-90%, and white = 50-75%; R = purines: A or G. Y = pyrimidines: U or C; P1 = Pairing Element. **(C)** Covariation motif of transcriptional fluoride riboswitches predicted to undergo intrinsic termination in the Holo (ligand-bound, **left**) or Apo (ligand-free, **right**). The two alternative stems are depicted as PK (pseudoknot) and P4 (terminating stem). The structures have been derived from Figure S22A showing the full R-scape output. **(D)** Holo (ligand-bound, **left**) or Apo (ligand-free, **right**) configurations for variants measured to terminate with the NGS assay. The two alternative stems are depicted as PK (pseudoknot) and P4 (terminating stem). The structures have been derived from Figure S22C showing the full R-scape output. The dashed line in C and D highlights the expression platform sequence known to be directly involved in the riboswitch structural switching mechanism. Covariation motifs adapted from raw output in Figure S22.

The covariation motif of the fluoride aptamer previously identified (RF01734) (45) contains three helixes covering at least three significant covarying base paired regions: P1, pseudoknot, P3 (Figure 5B). In comparison, the covariation model using the transcriptional riboswitch variants predicted by ARNold largely recreated the aptamer model by identifying P1, the pseudoknot, and P3 (Figure 5C). There are slight changes in the length of the base paired (P) regions and more overall sequence conservation within the aptamer (Figure 5C). The expression platform model did result in an intrinsic terminator motif that in the apo state includes a hairpin and a downstream polyU motif (Figure 5C). Notably, purines are preferred on the 3’ end of the terminator, adjacent to the polyU, as similarly observed for the purine (34) and SAM (35) riboswitch covariation motifs. We also identified sequence covariation along the terminator involving nucleotides that comprise the pseudoknot in the holo state (Figure 5C, dashed line), in accordance with several studies on the riboswitch structural rearrangement mechanism (4,38,39,60). A model utilizing sequences measured through NGS to be transcriptional riboswitch variants resulted in a similar model, but with overall less covariation (Figure 5D), potentially due to the increased number of sequences.

Overall, our bioinformatic approach demonstrates a high-throughput and accessible construction method of a covariation model for a complete riboswitch by filtering for a specific expression platform function. The fact that this model contains elements consistent with the known switching mechanism of the fluoride riboswitch suggests this could be a powerful approach to uncovering the mechanism of other riboswitch classes.

### Covariation models of full riboswitch aptamer and expression platforms for other classes reveal common riboswitch aptamer and downstream expression platform motifs

The similar riboswitch covariation motifs we observed between the predicted and measured pools of terminators supports the use of ARNold for rapid and high-throughput generation of complete covariation motif for other riboswitch classes. We therefore applied our approach to generate the complete motifs for several additional riboswitch classes: purine (34) and SAM (35) for their prior covariation characterization; ZTP, which has been extensively studied in a cotranscriptional context (6,64); Lysine, which regulates transcription termination (65–67), translational sequestering and RNase E cleavage (68); and TPP classes, which is the most common riboswitch and occurs throughout all three kingdoms of life (20).

To expand our analysis beyond transcriptional riboswitches, we further refined the method by calibrating the ARNold-generated covariation motif of the fluoride riboswitch, consisting of the aptamer and the downstream terminator hairpin, to increase sensitivity and speed in motif searching through sequences by determining the expected value scores and generating filter score cutoffs (cmcalibrate, see Methods and Materials). We then searched all the extended sequences in the riboswitch class for the aptamer+hairpin motif (Figure S23A). For the fluoride riboswitch, searching all variants did not produce novel information from the previous results, although the polyU track was reduced to a string of five pyrimidines (Figure 5C, 5D, Figure S23B) suggesting that this covariation motif analysis is able to fit a more diverse set of sequences. We interpreted this as the motif capturing all instances of aptamer + hairpin that included both transcriptional terminator expression platforms and translational expression platforms that consist of sequestering hairpins.

We next performed the covariation motif analysis for the purine (34) and SAM (35) riboswitches. For the purine riboswitch, we initially used 101 sequences identified by ARNold to generate a detective model and then used this model to search all 2611 sequences from Rfam (RF00167). The resulting model recreated the aptamer and revealed a downstream hairpin with 19 out of 21 base pairs covarying (Figure S24A). Notably, the previously published purine covariation motif generated through manually curating an alignment of likely transcriptional variants has higher sequence specificity in the expression platform, likely due to filtering down to 37 variants compared to the 2616 variants in our model (34). For the SAM riboswitch, we created an initial covariation model with 132 sequences identified from ARNold and then searched the 6114 pulled from Rfam (RF00162) (Figure S24B). Compared to the previously published SAM riboswitch motif (35), we observed the stretch of pyrimidine-purine base pairing pattern in the terminator ahead the polyU but did not observe significant covariation between the aptamer and downstream expression platform. These results demonstrate our method’s high tolerance for sequence diversity when detecting expression platforms.

We next generated novel riboswitch covariation motif for additional classes: ZTP riboswitch (Figure 6A), lysine riboswitch (Figure 6B), and TPP riboswitch (Figure 6C). For the ZTP riboswitch, we created an initial covariation motif from 187 sequences identified from ARNold (Figure S25A) and then used it to search the 3458 sequences pulled from Rfam (RF01750) (Figure 6A). The resulting model showed a rearrangement of the aptamer by the expression platform, which is supported from previous structural studies (6,64). For the lysine riboswitch, the covariation motif was generated from 40 sequences identified from ARNold (Figure S25B) and then used to search the 3458 sequences pulled from Rfam (RF00168) (Figure 6B). Interestingly, the aptamer-expression platform covariation overlap was found to include more of the aptamer than present within the *B. subtilis* lysC transcriptional model system (65–67) but matches the *E. coli* lysC translational variant (68), potentially providing insights into the more common regulatory mechanism. Thirdly, we generated a motif for the highly occurring TPP riboswitch, which similar to ZTP has been used to characterize RNA structural rearrangements outside a cotranscriptional regime context (5). We created a covariation motif of the TPP riboswitch from 48 sequences identified from ARNold (Figure S25C) and then searched the 16107 sequences pulled from Rfam (RF00059) (Figure 6C). The resulting model had only two nucleotides shared between the aptamer and expression platform (Figure 6C), which support the previous rearrangement mechanism (5).

**Figure 6.**
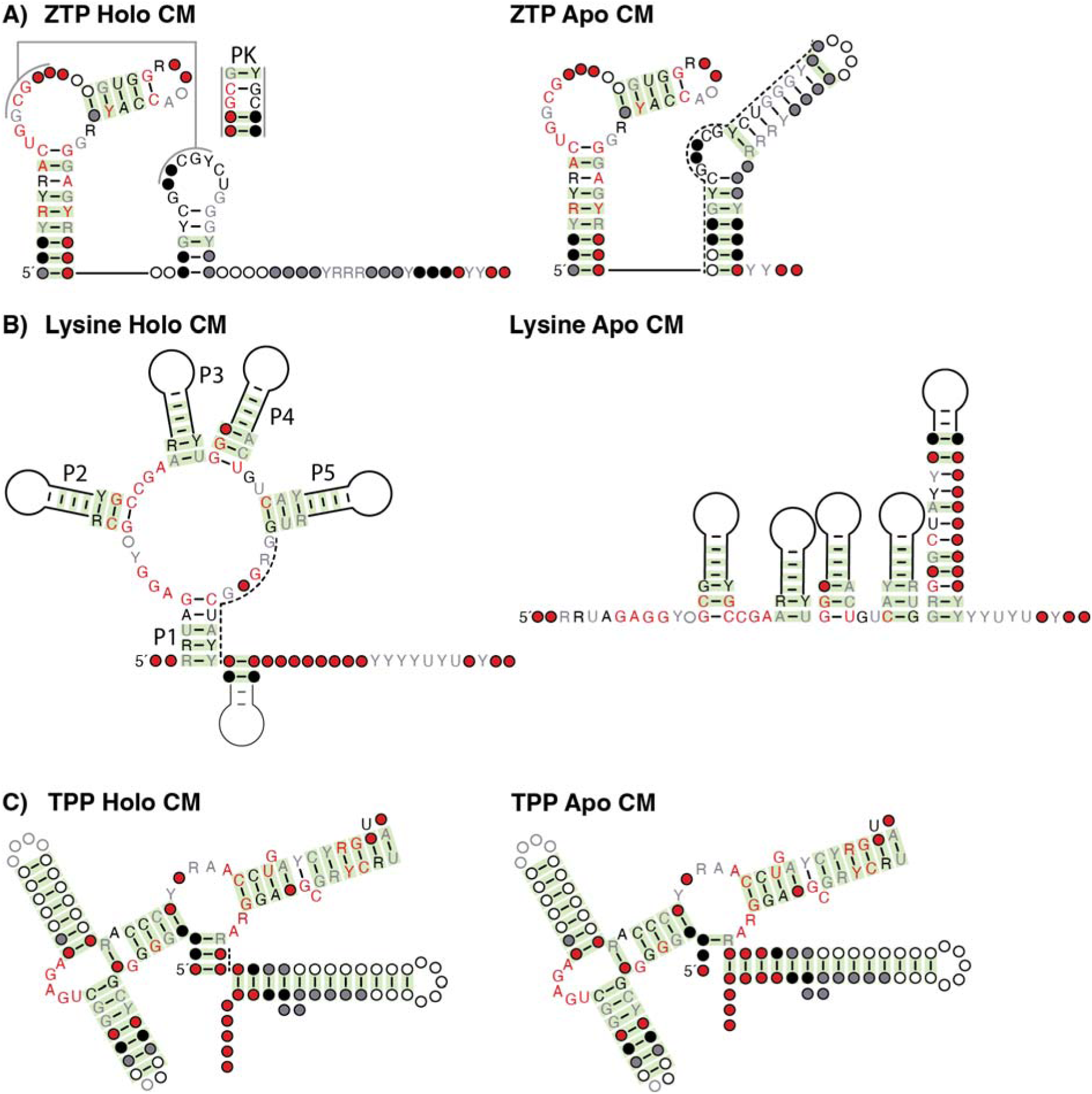
Covariation models of aptamer and expression platform sequences for ZTP, lysine, and TPP transcriptional riboswitches. Covariation models were generated as in Figure 5 using the ARNold computational prediction to filter for transcriptional sequence variants. The models were then calibrated and searched for through all downloaded sequences of each riboswitch class (Figure S23). **(A)** The ZTP (RF01750) riboswitch in the holo (ligand-bound, **left**) or apo (ligand-free, **right**) state. **(B)** The Lysine (RF00168) riboswitch in the holo (ligand-bound, **left**) or apo (ligand-free, **right**) state. **(C)** The TPP (RF00059) riboswitch in the holo (ligand-bound, **left**) or apo (ligand-free, **right**) state. Green highlighting on base pairs denotes evolutionarily significance in base pair covariation, and nucleotides are denoted as circle or specific letters with colors to signify conservation: red = > 97%, black = 90-97%, grey = 75-90%, and white = 50-75%; R = purines: A or G. Y = pyrimidines: U or C; P1 = Pairing Element. The initial R-scape/CaCoFold outputs from the ARNold sequences are in Supplemental Figure S25. The raw output from CaCoFold is in Supplemental Document C.

We then generated the covariation motif for the glmS riboswitch (Figure S26), which uniquely regulates gene expression through transcript cleavage rather than an expression platform (69). We sought to observe if there may be glmS riboswitch variants with an expression platform. While ARNold does identify terminators after the glmS aptamer, there is no covarying overlap between the aptamer or expression platform (Figure S26), suggesting that an expression platform method of gene regulation is not commonly used by the glmS aptamer.

Ultimately, this demonstrates that our method can be used to generate covariation models of riboswitches that include both aptamer and expression platform, and that these models can contain mechanistic insights into how riboswitches structurally rearrange between the aptamer and expression platform.

## DISCUSSION

In this work, we developed high-throughput methods to generate covariation models of complete riboswitches to investigate the evolutionary conservation of structural switching mechanisms. The core method consisted of a bioinformatic pipeline to acquire full riboswitch sequences predicted to undergo transcription termination (Figure 1); high-throughput characterization of riboswitch function using NGS readout (Figure 2); and use of identified transcriptional variants to build covariation models of riboswitch sequences (Figure 5). Application of this NGS pipeline to the fluoride riboswitch class identified 323/1901 (17%) fluoride riboswitch sequences tested were functional in our transcriptional assay. This shows that there are large pools of candidate riboswitch sequences that can be used to study mechanism beyond the handful of examples typically studied (4,38,39,60). Furthermore, the covariation model of the complete fluoride riboswitch sequence showed conservation patterns in the expression platform that directly overlap with those of the aptamer, consistent with the previously uncovered mechanism of the fluoride riboswitch entailing switching between mutually exclusive aptamer/expression platform folds (60).

One limitation of our approach is that we utilized *in vitro* transcription with the *E. coli* RNAP. As polymerases from other species can respond differently to transcriptional pause signals (70), and have different requirements for intrinsic transcriptional termination (71,72), the measured function for variants in our assay would likely change if different transcription conditions, or even cellular assays, were used. Although, it is highly notable even with these considerations that the Actinomycetota phylum in the Bacillati kingdom is nearly all translational with the other Bacillati phylum subclasses being mostly transcriptional (Figure S27). However, the use of a non-native polymerase limits the application of this pipeline to assessing non-*E. coli* anti-biotic targeting (Figure S28) (7,8). Secondly, the native transcript start site for most of the variants analyzed is unknown and the 5’ sequence ahead of riboswitches can impact their function (31,32,73). Thus, variant functionality may change with differences in the pre-aptamer leader sequence. Ultimately, our method is specifically suited to identifying functional riboswitch variants in an in vitro context rather than predicting *in cellulo* function.

We also found that riboswitch function measured by NGS was in-line with manual IVT validation but not in complete agreement. We hypothesize that this is due to biases in NGS library prep that allow shorter sequence products, i.e. terminated products, to be amplified and bind to the flow cell overcompensating the pool (Figure S7, Supplemental Document A) (63). This may explain the discrepancy in dynamic range between the high- and low-throughput comparisons (Figure 4E). Recently an approach called TECDisplay was developed (74) that uses an alternative approach to address this bias.

One benefit of our approach is that it can help uncover patterns that deviate from the established understanding of riboswitch mechanisms. During transcription, nascent RNAs can undergo dynamic structure rearrangements, which riboswitches have been an ideal model system to study. For the fluoride riboswitch, this dynamic rearrangement occurred through strand exchange of the six nucleotides in the pseudoknotted aptamer into the base of the terminator for the *B. ce* and *T. pe* variants (4,60,75). This mechanism is supported through covariation analysis (Figure 5C, D). However, our high-throughput analysis also revealed several functional variants of the fluoride riboswitch that extend through the aptamer auxiliary helix, P3 (Figure S16A, S16B, S17D, S17E, S17G), rather than just the aptamer pseudoknot as observed for *B.ce* and *T. pe*. Such evolution context and exception may be useful in understanding how folding pathways change when an aptamer switches between controlling transcription vs translation (76,77).

Our high-throughput approach also allowed us to investigate the effect of transcription factors such as GreB on the function of many riboswitch variants all at once. There is a growing body of work studying the dynamic relationship between riboswitches and transcription factors (39,43,44) such as GreB, which rescues backtracked *E. coli* RNAP through cleaving the 3’ end of the RNA (42). In our assay, we found that the presence of GreB can have selected impact on a subset of fluoride riboswitch sequences. Notably, GreB engages during class II pauses (78), which differs for the class I pause typically observed for intrinsic termination (29), suggesting that our method may reveal novel principles of molecular biology that are unseen at the single-model case study. Towards the single-model case study, the nucleotide-amino acid interactions between a riboswitch and RNAP exit channel was mapped (79) but notably the nucleotides that interact with the RNAP amino acids are not in highly conserved positions of the aptamer. This highlights the sequence-specific, rather than generalizable, ways nascent transcripts fold and influence transcription progression.

The ease of the computational approach allowed us to apply the method to other riboswitch classes. In total, we generated six of the current 51 riboswitches on Rfam. Our full covariation motifs validated similar mechanisms seen previously (5,6,35,64,68). Thus, such an approach could potentially be used to discover mechanisms for other riboswitches.

Central to this work was the employment of massive parallel oligo synthesis (MPOS). MPOS was previously used to test the structural tolerance of the Twister ribozyme and covariation model searching was employed to identify an active ribozyme in mammals (80). While we have not similarly searched through other genomes for the fluoride riboswitch, we do note that there are fluoride aptamer sequences deposited on Rfam from *Drosophila ficusphila* (KB456680.1), *Spinacia oleracea* (AYZV02264506.1), wheat (*Triticum Urartu*, AOTI010596493.1), and rice (*Oryza sativa Indica Group*, CH399130.1). Our NGS assay shows functional variants from Spinacia and wheat, however validation in the organism systems is needed given how transcription machinery changes throughout evolution (70,81). Furthermore, this MPOS and NGS assay may accelerate addressing the orphan riboswitch problem (20) by screening a large number of riboswitches during candidate ligand searches.

Ultimately, this work progresses covariation analysis for rapid detection of dynamic RNA confirmation states. These pipelines will aid our search for new riboswitch mechanisms, studies for antibiotic targeting, and developing riboswitches for novel biotechnologies.

## Supporting information

Output from bioinformatics, python read analysis, read calculations, and gel quantification.

Output for the in vitro fluorescence assay.

Unedited output from R2R with Rscape.

Unedited phylogenetic trees.

## DATA AND CODE AVAILABILITY

The scripts to run the analysis is available at GitHub: github.com/LucksLab/Hertz_HighTroughput_Riboswitch_Discovery_2025. Code was written with the assistance of ChatGTP (v. 3 and v. 4) and manually validated and edited.

Accession numbers for all sequences are in Supplemental_Document_A.xlsx. The next generation sequencing data is deposited on the Sequence Read Archive (SRA) managed by the NCBI under BioProject: PRJNA1359411.

## SUPPLMENTAL DOCUMENTATION

Supplemental Documentation and Data are available online.

Supplementary Information – Supplementary Figures and Tables.

Supplementary_Document_A.xlsx – Output from bioinformatics, python read analysis, read calculations, and gel quantification.

Supplemental_Document_B.xlsx – Output for the in vitro fluorescence assay..

Supplemental_Document_C.pdf – Unedited output from R2R with Rscape.

Supplemental_Document_D.pdf – Unedited phylogenetic trees.

## AUTHOR CONTRIBUTIONS

Conceptualization: L.M.H., J.B.L. Experiments and data analysis: L.M.H. GreB purification and florescence assay: A.A.M. Covariation analysis: L.M.H., E. R. Writing (original draft): L.M.H. Editing: all authors.

## ACKNOWELEDGMENTS

We thank: Irina Artsimovich for providing the RNAPT7-driven expression plasmids of C-terminal his-tagged GreB (pAI577) and advice for protein purification. Edric Choi and Tina Fu for discussions about Next Generation Sequencing preparation. Reese Richardson and Edric Choi for coding assistance. Xavier Bower for discussions on analyzing the NGS data for GreB mechanism. Herma Demissie for manuscript feedback. The Northwestern University IT department for the BYOD groups and one-on-one consultations as well as the Writing Place for the writing feedback group.

## FUNDING

Research reported in this publication was further supported by NIGMS of the National Institutes of Health under award number R01GM130901 to J.B.L and Biotechnology Training Program via NIH training grant T32GM008449 to L.M.H. This work was also supported by the National Science Foundation’s MRSEC program (DMR-2308691) at the Materials Research Center of Northwestern University. The content is solely the responsibility of the authors and does not necessarily represent the official views of the National Science Foundation or the National Institutes of Health. L.M.H. gratefully acknowledges support from the Ryan Fellowship and the International Institute for Nanotechnology at Northwestern University. This work was partially supported by NIH grant R01GM144423 to E.R. Pew Latin American Fellowship 2022 partially supported A.A.M. in this work.

## CONFLICTS OF INTEREST

None declared.

## Notes

### Competing Interest Statement

The authors have declared no competing interest.

