## Supplementary Figures and Tables for "High-throughput functional profiling and evolutionary covariation analysis of entire riboswitch sequences"

**Table 3: Index list.** 34
**Table 4: ARNold averages for the terminator results.** 34
**Table 5: Pearson correlation coefficient (r)** 34
**Table 6: Key to Figure 4E** 35
*RAW GEL IMAGES* *36*
**Pre-sequencing gel image, Figure S2A** 36
***In vitro* transcription RNA products, Figure S18, S19, S20** 37
**In vitro transcription RNA products, Figure S20, S21** 38
***In vitro* transcription RNA products, Figure S21** 39
*REFERENCES* *40*

**SUPPLEMENTAL FIGURES**

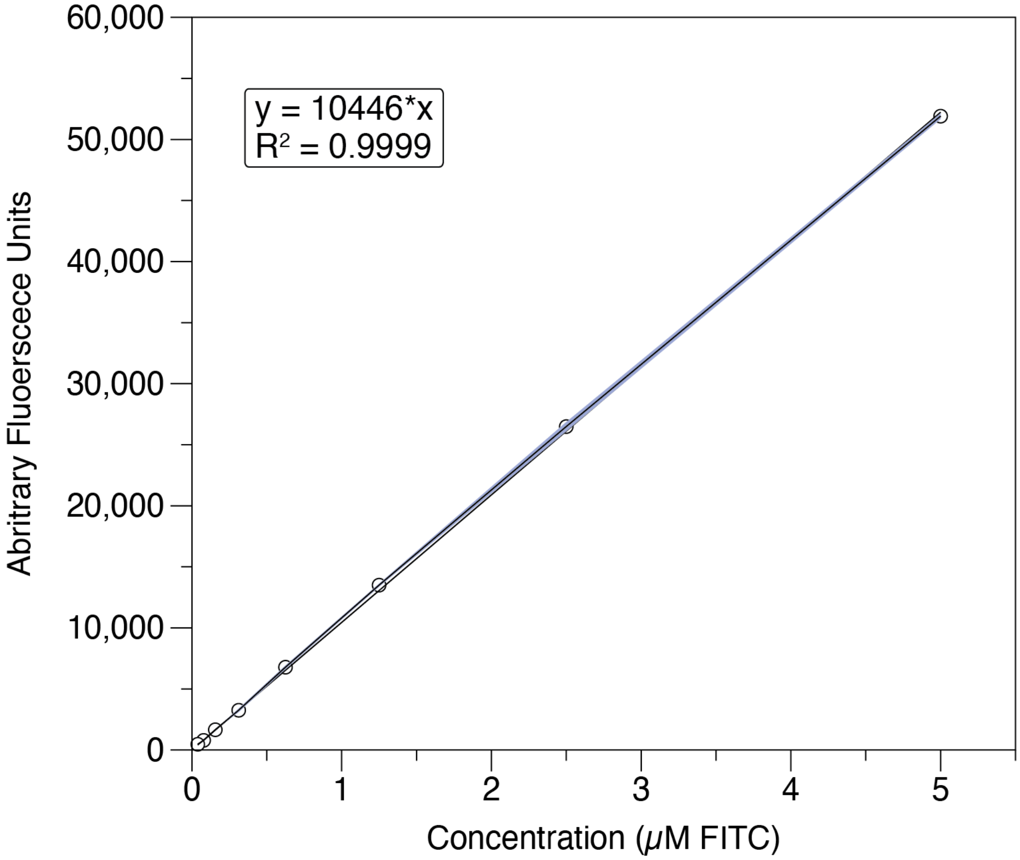

Figure S1. FITC calibration curve for fluorescence standardization. Fluorescence intensity (arbitrary units) was converted to micromolar fluorescein equivalents (µM FITC) using a NIST-traceable standard (see **Materials and Methods**). A series of two-fold serial dilutions starting at 5 µM FITC in buffer (100 mM sodium borate, pH 9.5), was prepared and measured using the same plate reader and its settings as for experimental samples (Ex 485 nm, Em 520 nm). A linear regression constrained to pass through the origin (0,0) was applied over the full dilution range, yielding strong linearity ($R^2 = 0.9999$). The shaded area around the fit line represents the standard deviation. Data points show mean three independent replicates.

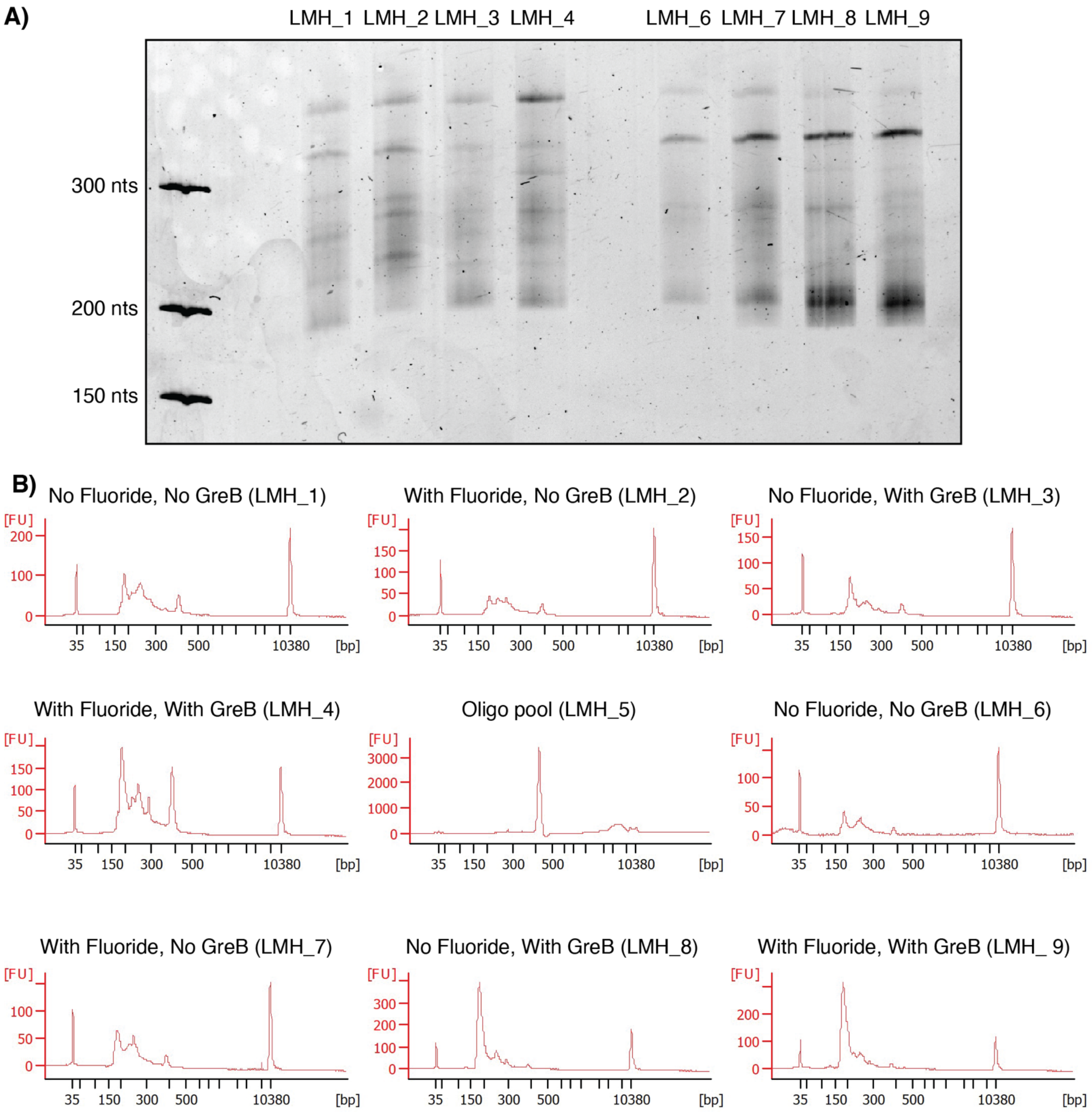

Figure S2. Quality controls for next-generation sequencing libraries. **(A)** An 8% denaturing gel run for 2 hrs at 18 W with the GeneRuler Ultra Low Range DNA Ladder (Thermo Scientific, SM1213). Raw gel image is provided in the "RAW GEL IMAGES" section. **(B)** Bioanalyzer high sensitivity DNA assays run on a 2100 bioanalyzer at the NUSeq core. The library names LMH_1 – 4 are replicate 1 and 6-9 are replicate 2. LMH_1 and 6 are 10 mM NaCl, LMH_2 and 7 are 10 mM NaF, LMH_3 and 8 are 10 mM NaCl with 1.2 µM GreB, and LMH_4 and 9 are 10 mM NaF with 1.2 µM GreB. LMH_5 is the library for directly sequencing the oligo pool.

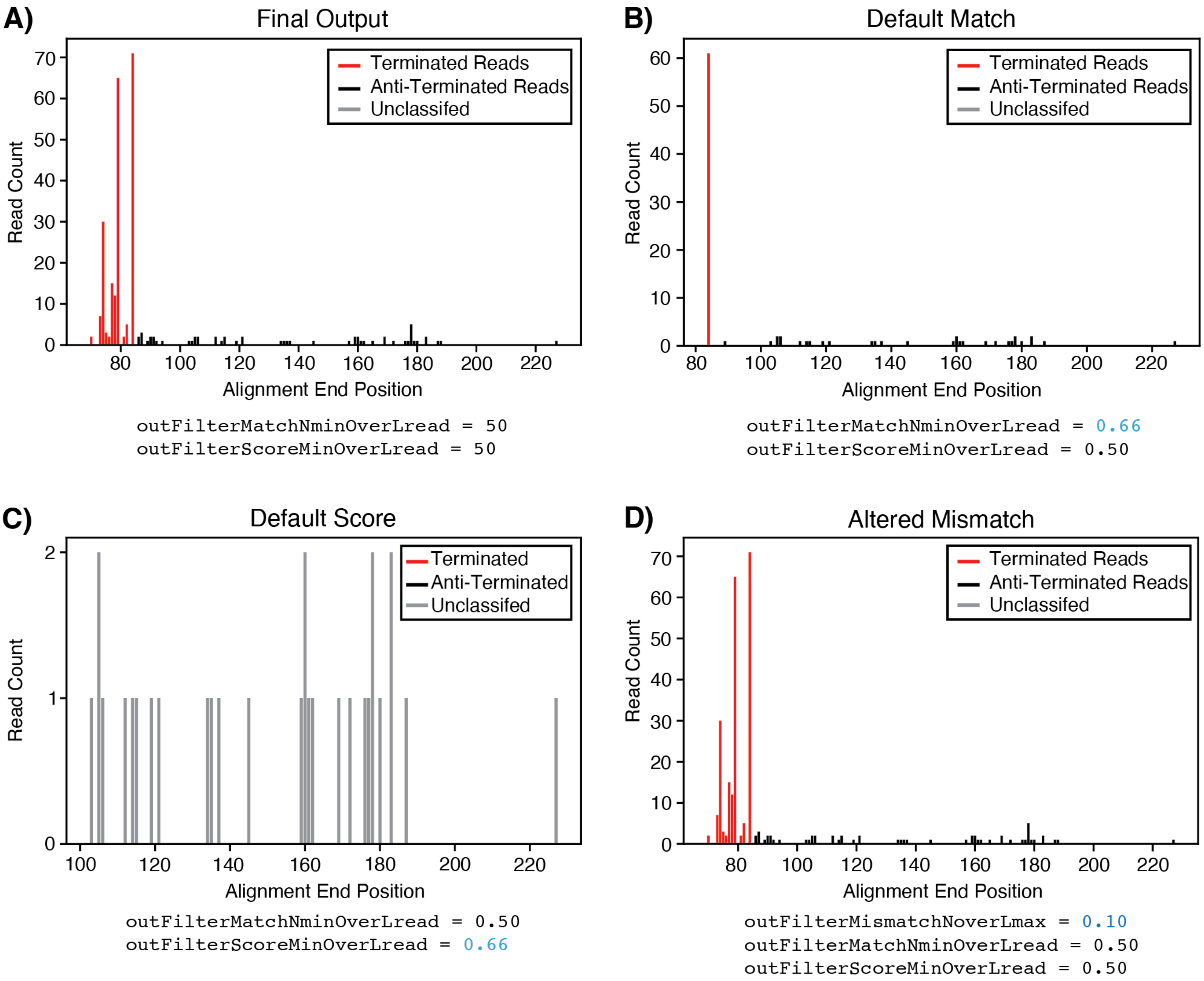

Figure S3. STAR parameter adjustments. Using LMH_1 and the positive control *Bacillus cereus* (CP000227.1/4763720-4763779), we altered several STAR alignment parameters. We adjusted the STAR alignment parameters since the reference genes are 235 nucleotides long, while the terminated B. ce reads are around 80 nucleotides. Without proper parameter adjustment, the shorter B. ce reads were being filtered out during alignment. A) STAR parameters used to align reads to reference genes. B) The same parameters as (A), but with the outFilterMatchNminOverLread set to its default value of 0.66 (blue text). Reads where the ratio of matched bases to reference bases is below this threshold are discarded. For instance, a 100 nt read with 70 matching bases is retained, but a read with only 60 matching bases is discarded. Without lowering this value, terminated reads were filtered out. C) The same parameters as (A), but with the outFilterScoreMinOverLread set to its default value of 0.66 (blue text). Reads with an alignment score normalized to the read length below this ratio are discarded. For example, a 100 nt read with an alignment score of 0.68 is retained, while a 100 nt read with a score of 0.54 is discarded. Without lowering this value, terminated reads were filtered out. D) Addition of the outFilterMismatchNoverLmax parameter lowered from the default of 0.3 to 0.1 (blue text). This filter discards reads where the ratio of mismatches to read length exceeds 0.3. For example, a 100 nt read with 32 mismatches would be discarded, while a 100 nt read with 28 mismatches would be retained. This filter was applied to account for potential overlap in aptamer sequences between fluoride riboswitch variants. Default parameters and detailed explanations are found in the STAR manual at:
https://physiology.med.cornell.edu/faculty/skrabanek/lab/angsd/lecture_notes/STARmanual.pdf

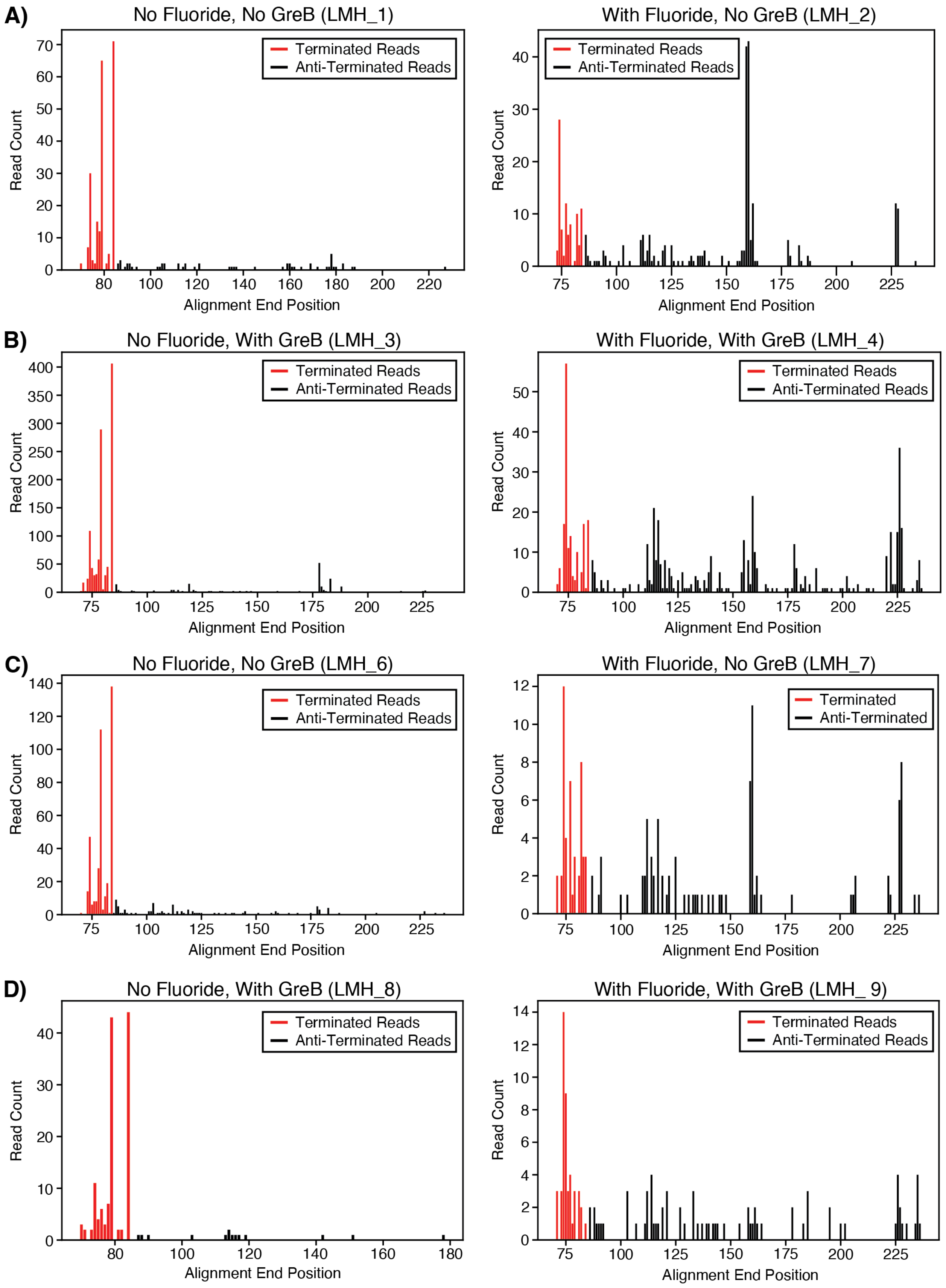

Figure S4. Histograms of read alignments to Bacillus cereus (CP000227.1/4763720-4763779) from all libraries. Reads were classified “Terminated” (red) if they ended from 2 nts before the pattern through 6 nts after the regular expression pattern: “'TTTTT|T[AGC]TTTT|TT[AGC]TTT|TTT[AGC]TT'”. Reads that came after the region were classified as “Anti-Terminated” (black). These counts were used to calculate the percent Anti-termination (%AT).

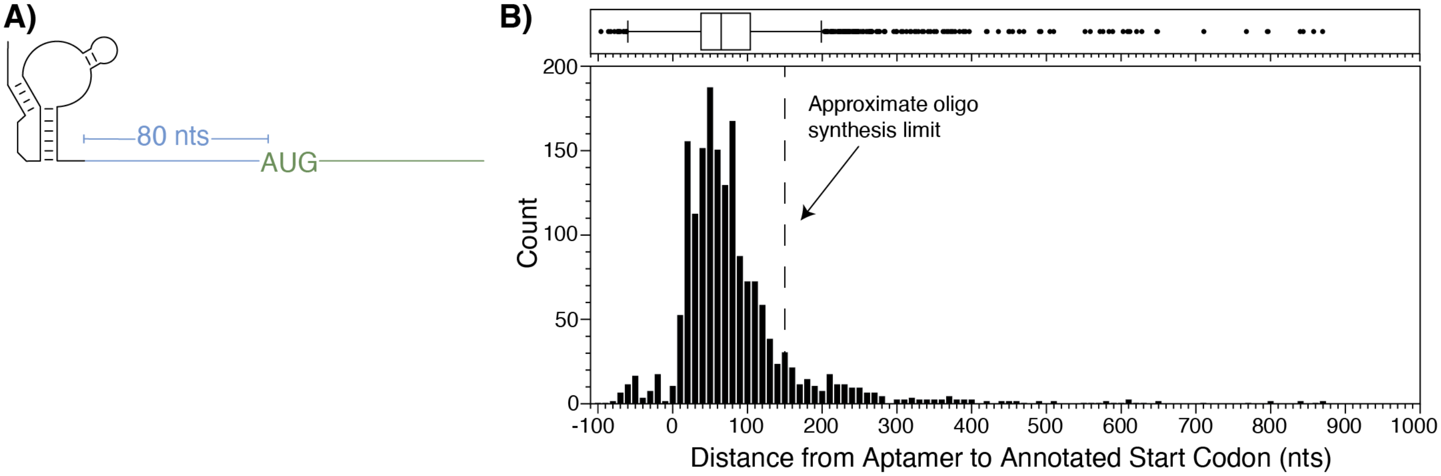

Figure S5. Histogram of distances from the end of each fluoride aptamer to the start codon of the nearest annotated genomic ORF. **(A)** Schematic describing the information gathered with the aptamer in black, expression platform in blue, and coding region in green. **(B)** Whisker plot (top) and histogram (bottom) of the distance between the aptamer end and start of the first downstream genomic ORF, i.e the blue region in the schematic in A. The dashed line at 150 nts marks the approximate sequence length added to each aptamer for oligo synthesis as determined by the technical synthesis length maximum of 300 nts: 300 nts = length(promoter)(60 nts) + length(aptamer)(~60 nts) + length(extended EP sequence)(~160 nts) + length(3' primer binding site)(20 nts).

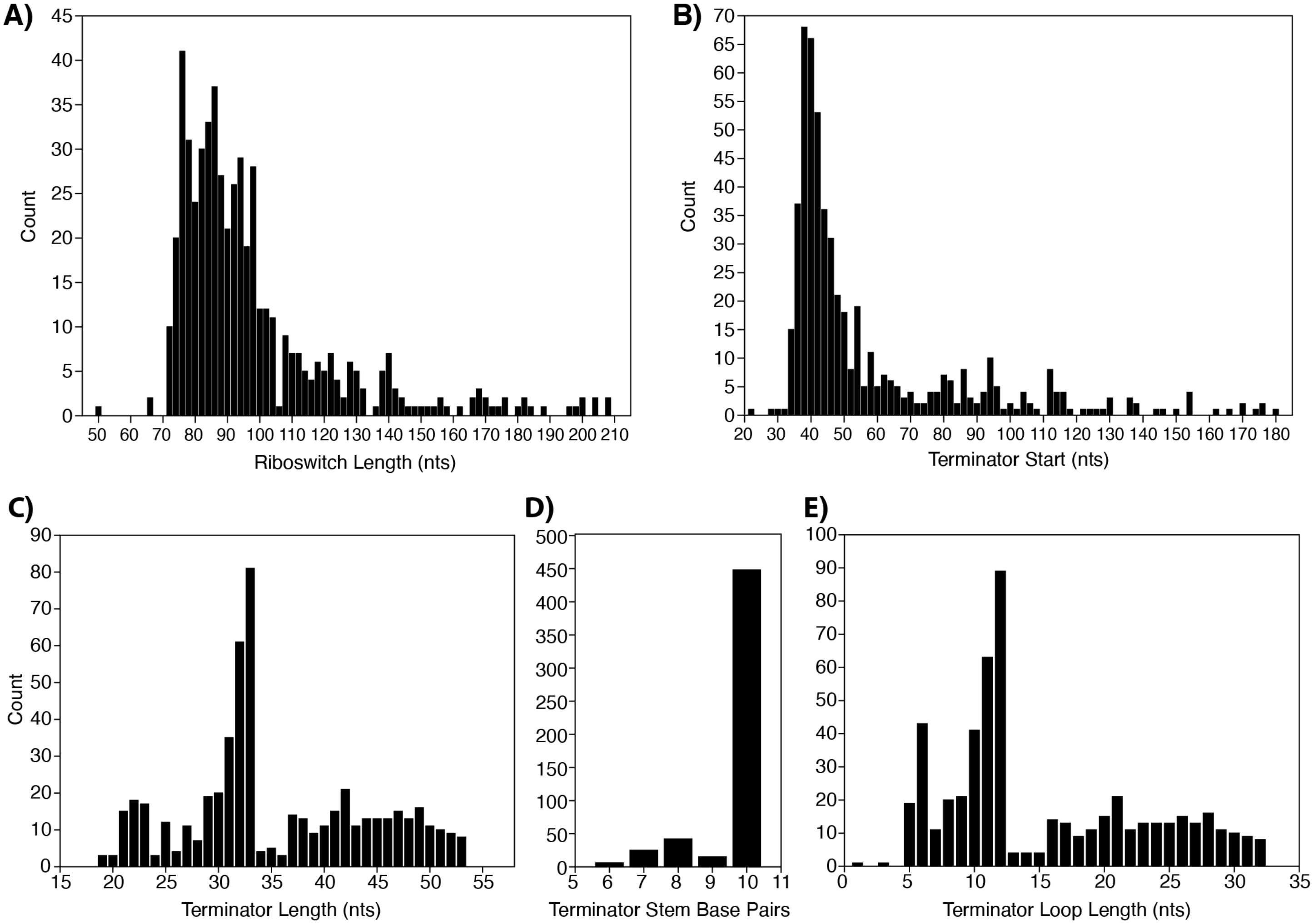

Figure S6. Terminator features predicted by ARNold over the 536 fluoride riboswitch variants predicted to regulate transcription. Histograms of: **(A)** number of nucleotides of the aptamer through the polyU of the predicted terminator, **(B)** the predicted 5' position of the terminator, **(C)** number of nucleotides in the predicted terminator, **(D)** number of base pairs in the terminator hairpin stem, and **(E)** number of nucleotides in the terminator loop. Data in Supplemental Document A.

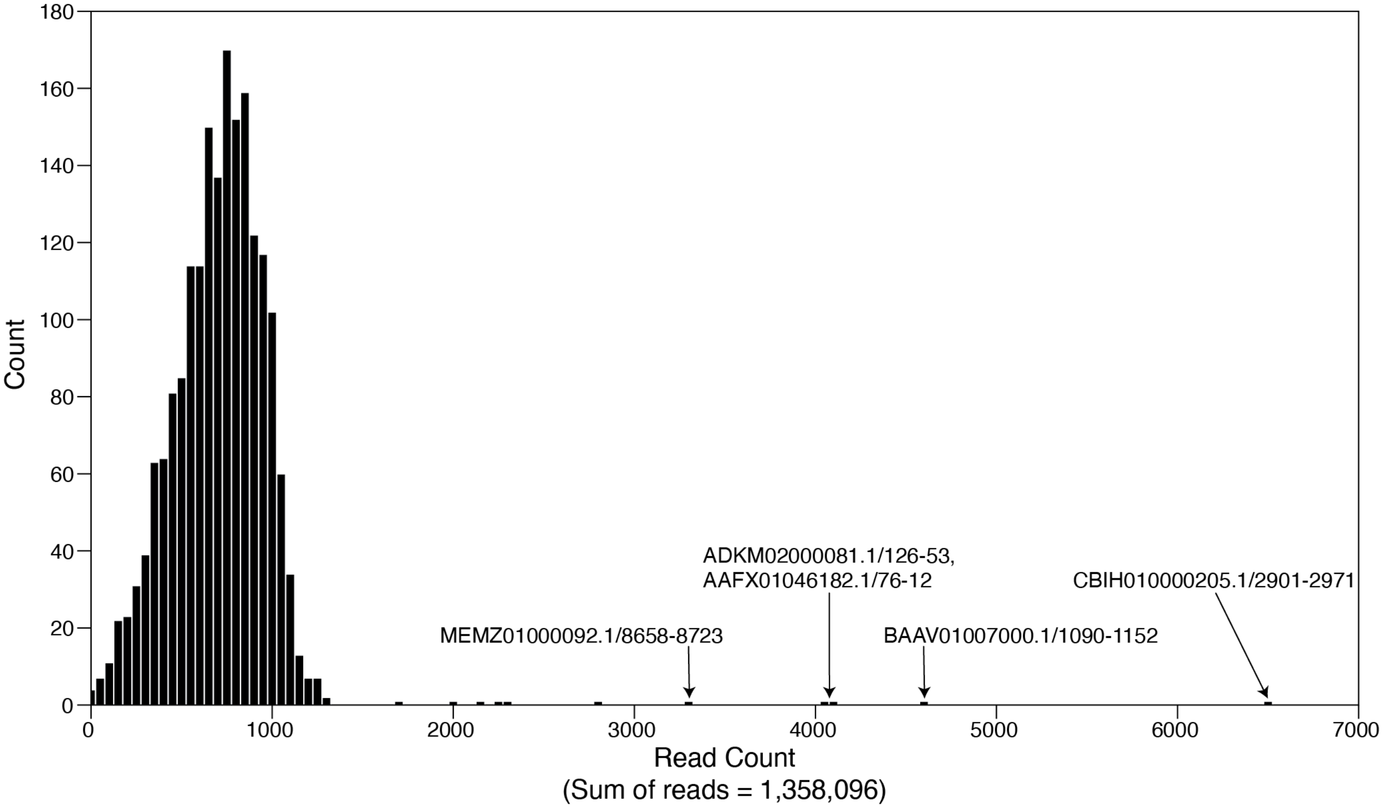

Figure S7. Histogram of read coverage of synthesized oligo pool. All 1901 sequences ordered were found in the oligo pool. Most sequences had fewer than 1000 reads (median reads = 726) with several labeled outliers over 3,000 reads: CBIH010000205.1/2901-2971 (6,520 reads), BAAV01007000.1/1090-1152 (4,622 reads), ADKM02000081.1/126-53 (4,120 reads), AAFX01046182.1/76-12 (4047 reads), MEMZ01000092.1/8658-8723 (3,303 reads). The over representation could be from oligo pool synthesis of the NGS PCR library prep. However, Twist technical support noted that their synthesis is biased towards shorter products and several of these overrepresented sequences are the shortest within the oligo pool. Data in Supplemental Document A.

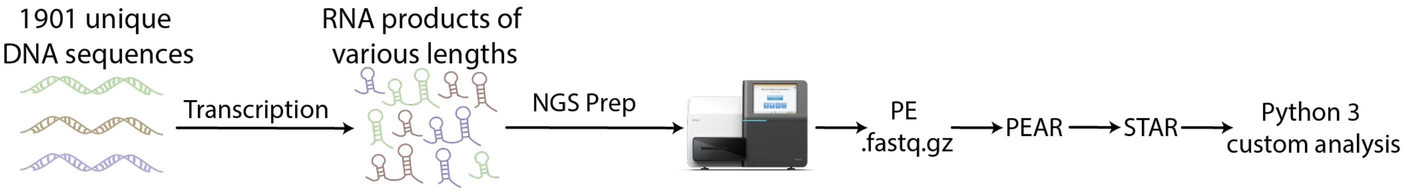

Figure S8. Overview of RNA sequencing and analysis pipeline. Schematic of the workflow that combines PEAR (v0.9.6) to combine pair-end (PE) reads, STAR (v2.7.9a) to align reads to the originating source, and Python 3 custom analysis scripts to quantify reads. Reads were mapped along the variant sequence to assess RNA transcript length. If a polyU motif was identified, then the reads in that region were classified as "Terminated" and the reads afterwards were "Anti-Terminated" (see Methods).

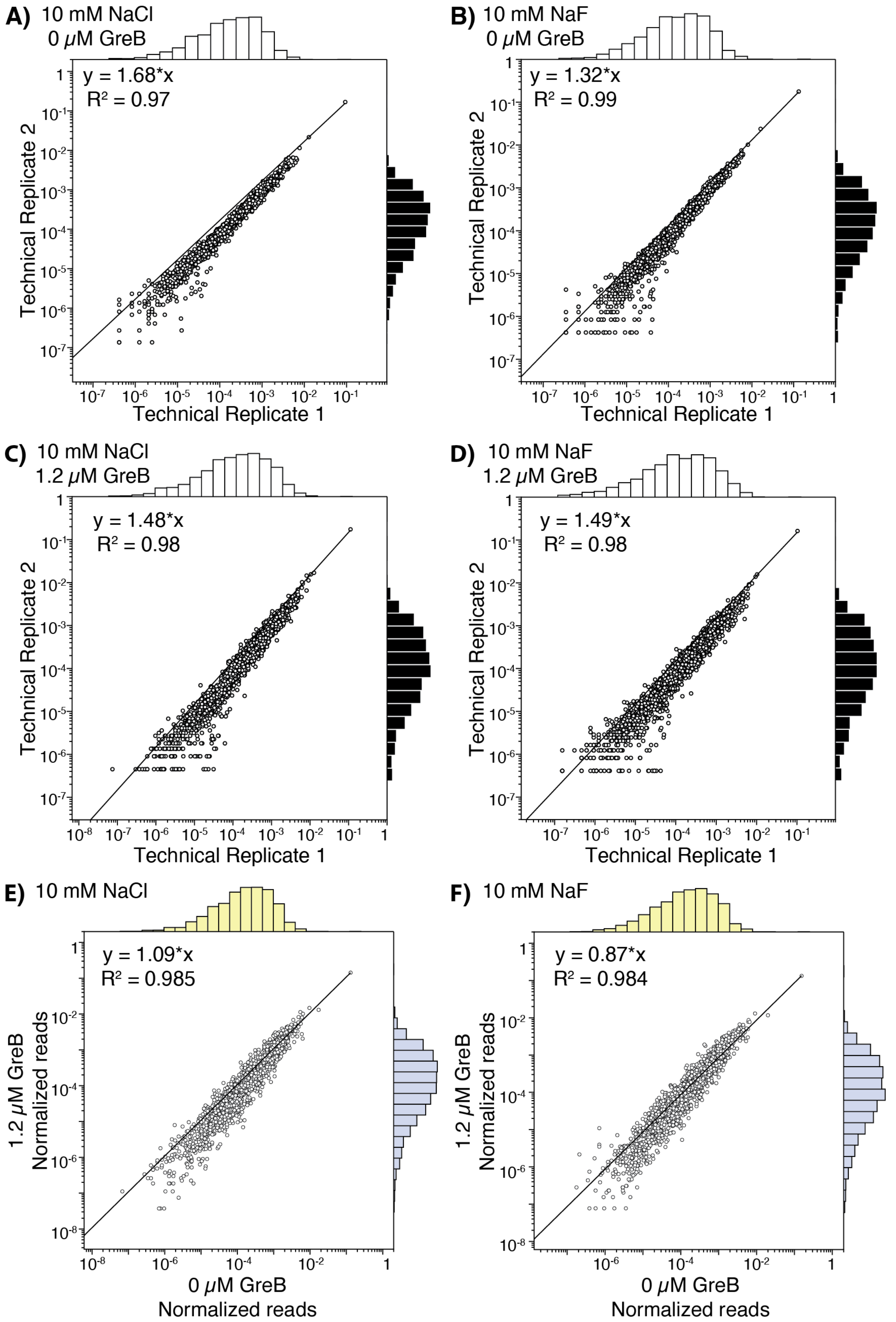

Figure S9. Correlation plots for the two technical NGS replicates across conditions. **(A-D)** Technical replicates with read counts ("Terminated" + "Anti-Terminated" variant read counts/ total sample technical reads) for all conditions: 10 mM NaCl, 0 µM GreB **(A)**, 10 mM NaF, 0 µM GreB **(B)**, 10 mM NaCl, 1.2 µM GreB **(C)**, 10 mM NaF, 1.2 µM GreB **(D)**. **(E-F)** The read counts for each variant are normalized by the total number of reads (both terminated and anti-terminated) for the sample, then averaged between the two technical replicates for 10 mM NaCl **(E)** and NaF **(F)** conditions. Colored distributions show distribution for each condition (yellow = 0 µM GreB, blue = 1.2 µM GreB).

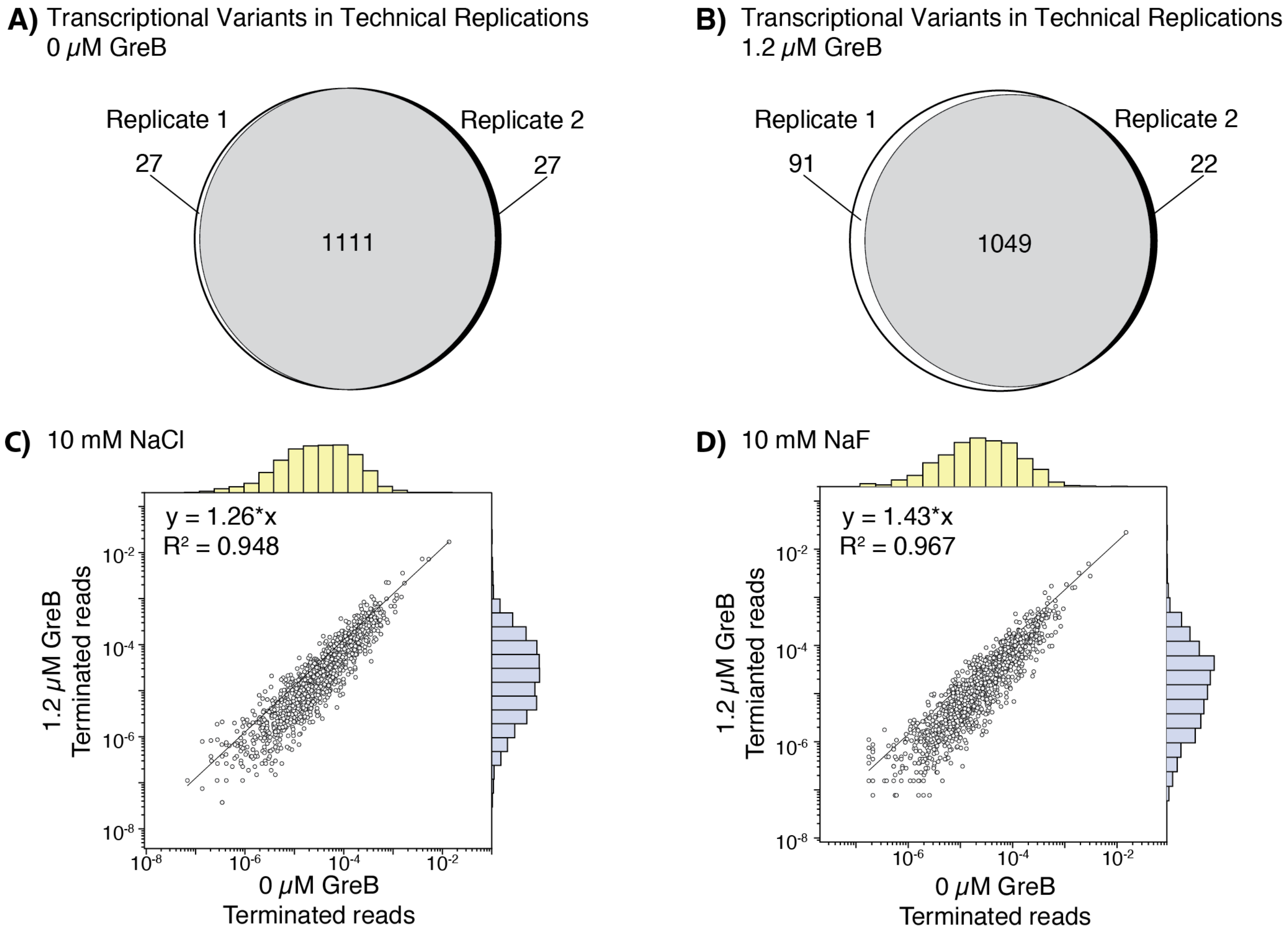

Figure S10. Assessing termination identification consistency across replicates. **(A-B)** Variants were classified in the two technical replicates as having a "terminated" read without **(A)** or with **(B)** GreB. **(C-D)** Correlation of normalized terminated reads between the two technical replicates in the without and with GreB for the variants in the NaCl (**C)** or NaF **(D)** conditions.

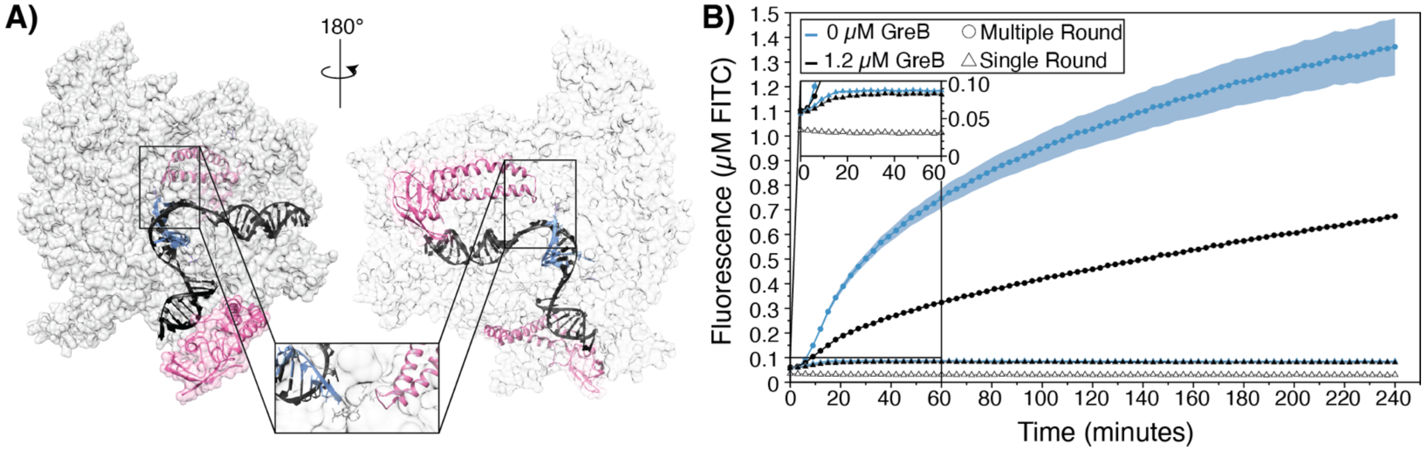

Figure S11. Time-course of fluorescent output assessing GreB transcriptional impact on IVT. **(A)** Structure of GreB (pink) on a transcriptional elongation complex; RNAP = grey, black = DNA, blue = RNA. PDB ID: 6RI7. **(B)** An IVT assay was used to transcribe a 3-way junction dimeric broccoli fluorescent aptamer in single (triangle) or multiple-round (circle) conditions with or without 1.2 µM GreB. Fluorescence was tracked for 2 hours and calibrated to µM FITC. A "No RNAP" control is plotted (white triangles). Points represent averages over three experimental replicates with two technical replicates for each sample (N = 6), and shading represents the standard deviation.  Data in Supplemental Document B.

**A)** Probability >= 99% | 90% > Probability >= 80% | 60% > Probability >= 50%
99% > Probability >= 95% | 80% > Probability >= 70% | 50% > Probability
95% > Probability >= 90% | 70% > Probability >= 60%

**B)** Terminators successfully predicted by ARNold

**C)** Terminators unsuccessfully predicted by ARNold

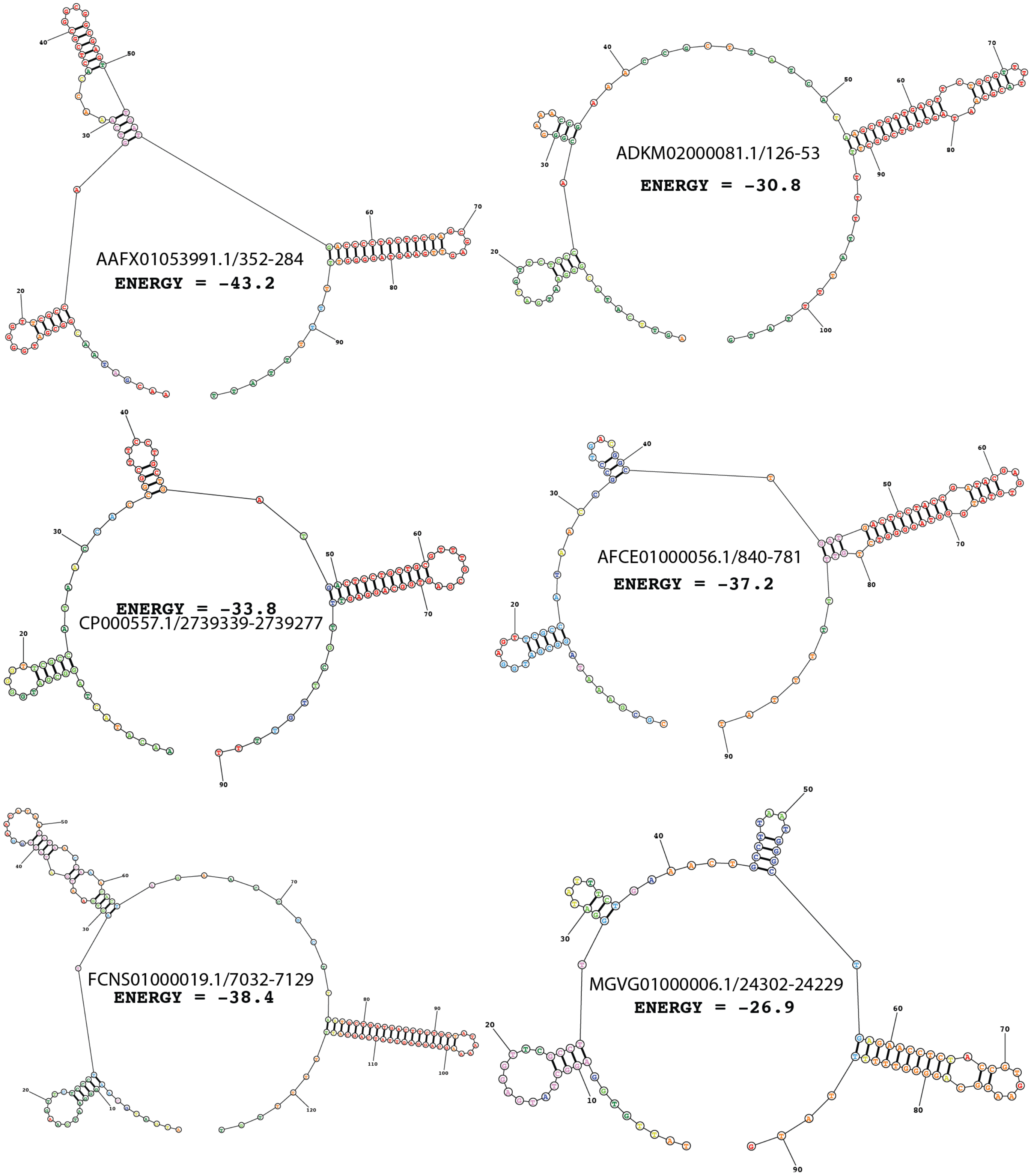

Figure S12. RNA structure analysis of successful and unsuccessful predicted terminators from RNAStructure. **(1) (A)** Color key code. **(B-C)** Structure predictions for three variants that were classified as "Terminated" in the NGS assay and either successfully **(B)** or unsuccessfully **(C)** predicted by ARNold. The species identifier and structure free energy is listed alongside the structure.

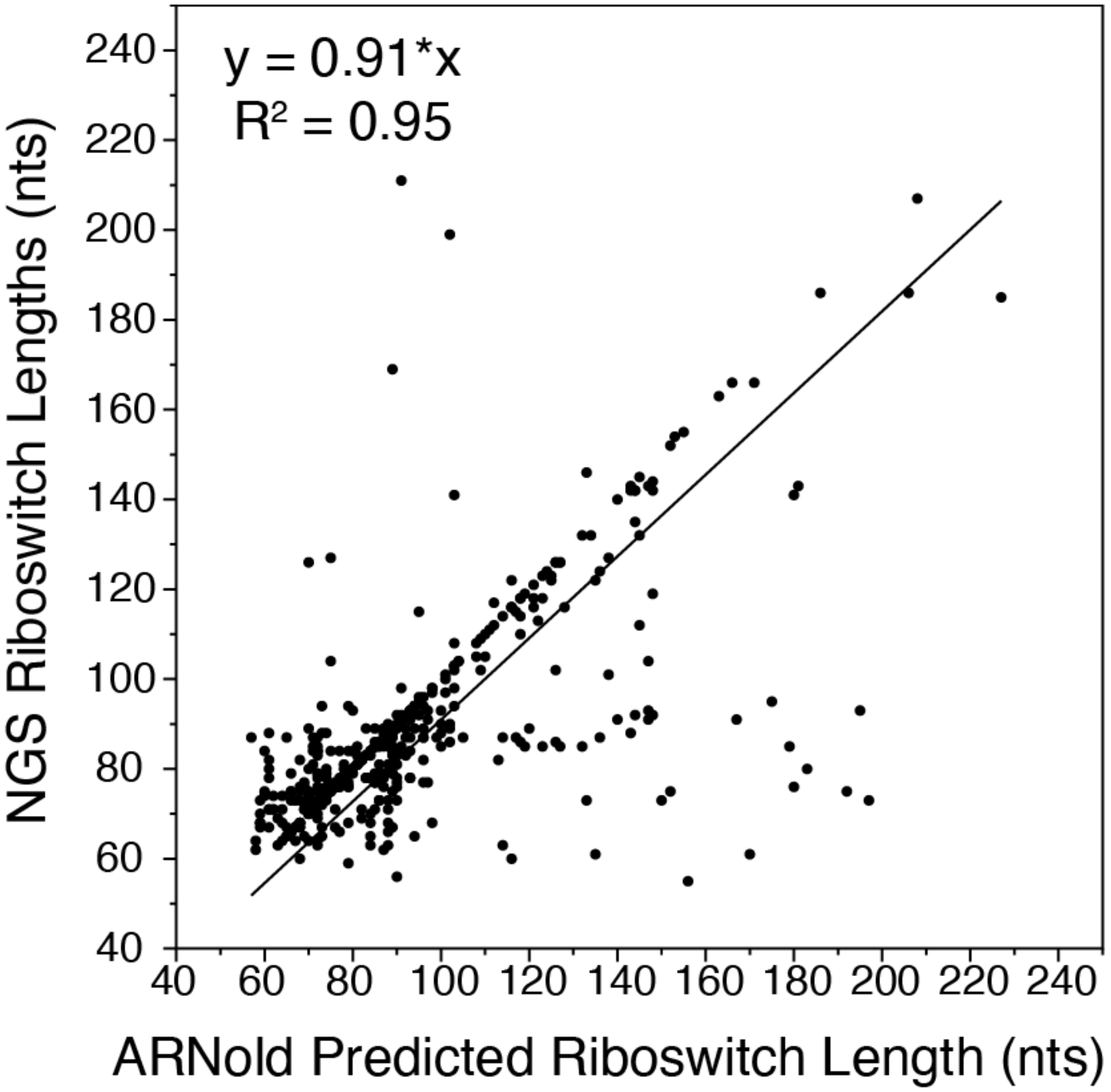

Figure S13. Correlation between the measured position of transcriptional termination (y-axis) and the predicted position using the ARNold webserver (x-axis). Riboswitch variants were plotted if they were both predicted and measured to undergo termination (N = 465). Measured riboswitch lengths were identified through bioinformatic analysis of sequencing reads (see Methods) and the ARNold predicted riboswitch lengths were calculated by adding the predicted terminator start site and length of the terminator. A linear fit showed an $R^2$ value of 0.95. Data in Supplemental Document A ('Predicted_Measured_Termination' sheet) and linear regression done in DataGraph (v4.5.1).

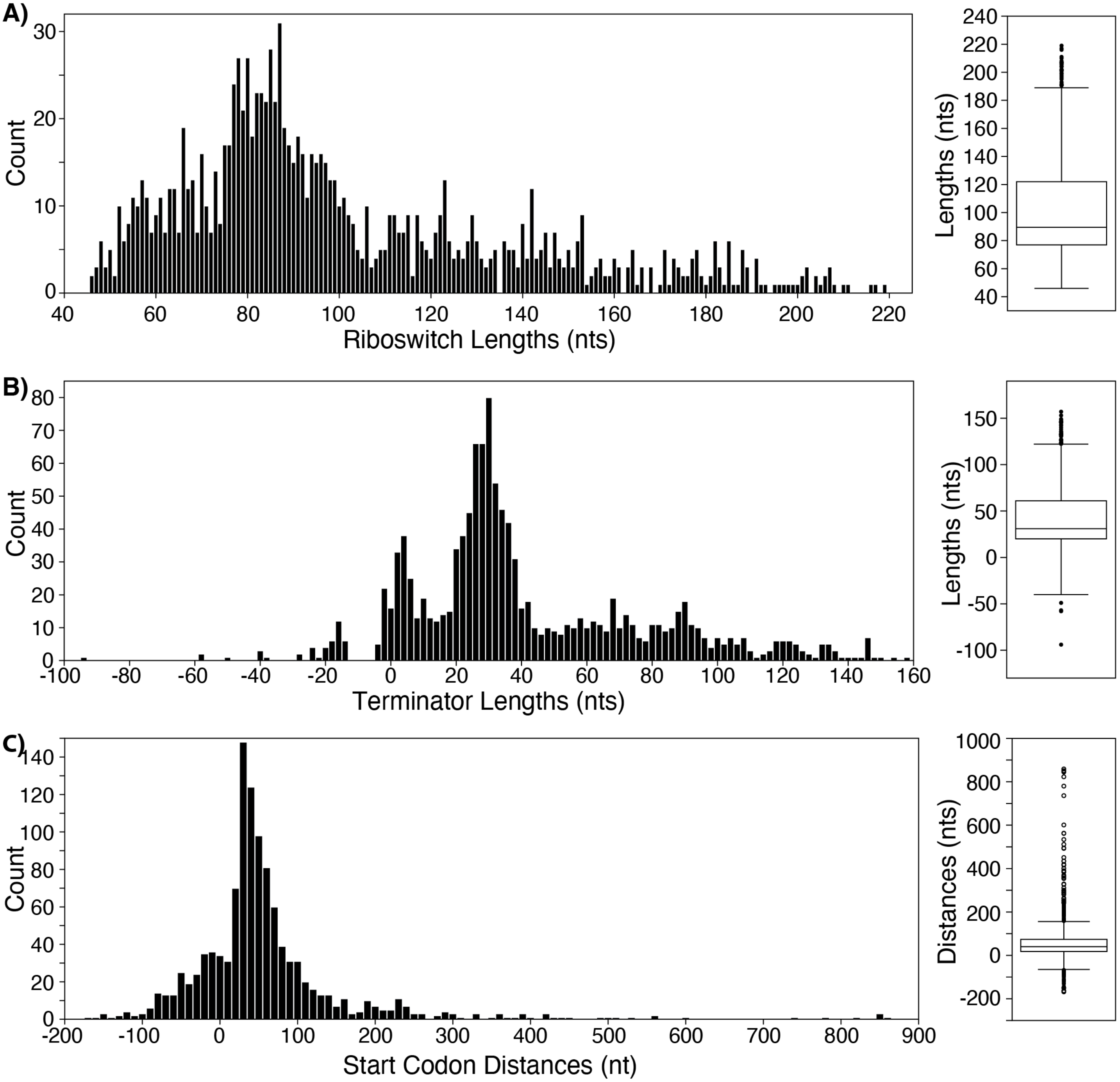

Figure S14. Histograms and whiskey plots of terminator features measured in the NGS assay: **(A)** Terminator length (nts) as calculated by subtracting the aptamer length from the length of each terminated read. Negative terminator lengths arise with long aptamers, like CP000975.1/1993796-1993891 which has an aptamer length of 97 nts and had a measured terminator site at 53 nts. Thus, these lengths are an approximation. **(B)** Length of the full riboswitch, including the aptamer, as chosen by the shortest terminated read length. **(C)** Distance between the end of the riboswitch and the start codon of the first downstream genomic ORF (Figure S5). Negative start codon distances reflect start codons that are within the aptamer. Data in Supplemental Document A.

**A)** CP000702.1/1794817-1795036 *Thermotoga petrophila RKU-1*

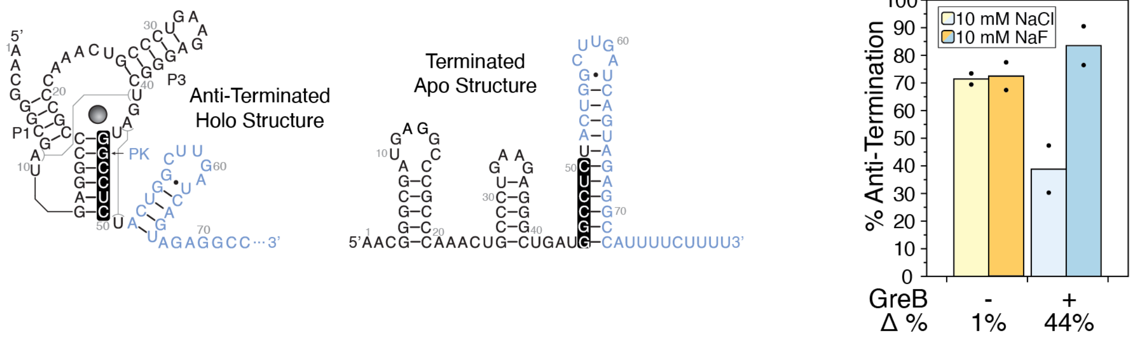

**B)** CP000227.1/4763720-4763939 *Bacillus cereus Q1*

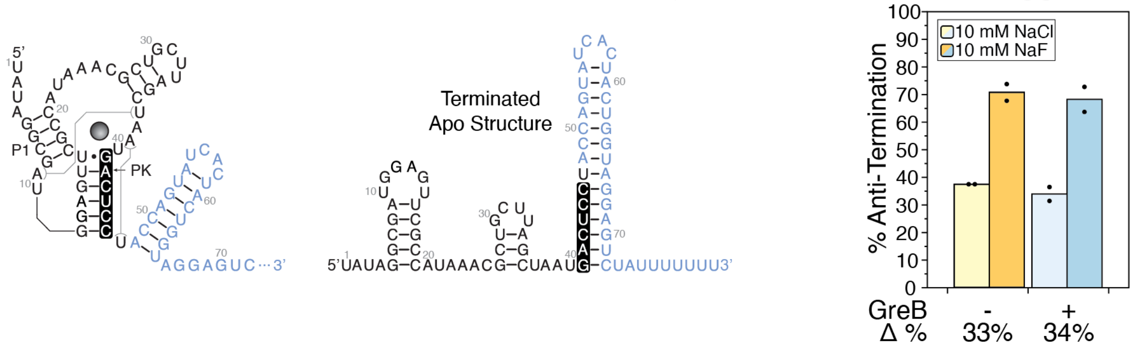

**C)** BX571966.1/2538786-2538786 *Burkholderia pseudomallei strain K96243*

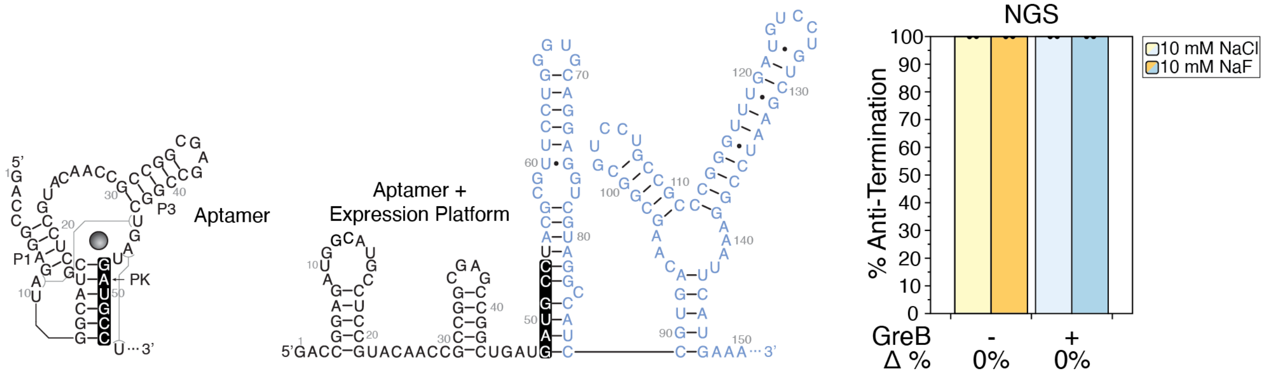

**D)** AE016853.1/5215490-5215709 *Pseudomonas syringae pv. tomato str. DC3000*

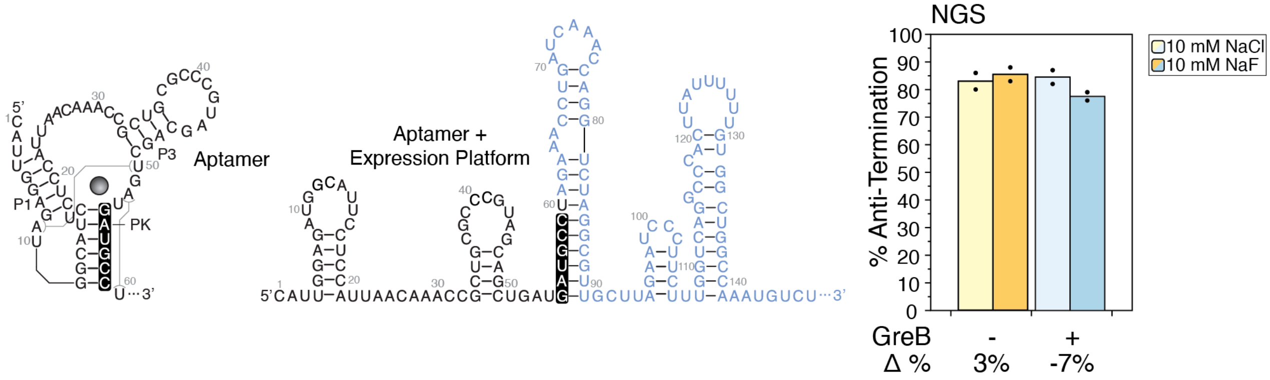

Figure S15. Fluoride riboswitch structures and NGS assay results without or with GreB for Figure 3. Hypothesized structures for the riboswitch variants. The structures were informed by the Rfam consensus structure and predictive modeling from RNAStructure. The bar graph depicts the percent (%) Anti-termination either without (0 µM, yellow) or with (1.2 µM, blue) the transcription elongation factor, GreB. The change (Δ) of % Anti-termination is written for each condition. **(A)** *Bacillus cereus* (accession ID: CP000227.1) **(B)** *Thermotoga petrophila* (accession ID: CP000702.1) **(C)** *Burkholderia pseudomallei* (accession ID: BX571966.1) **(D)** *Pseudomonas syringae* (accession ID: AE016853.1). Bars represent average % anti-termination with points plotted (N = 2). Data in Supplemental Document A.

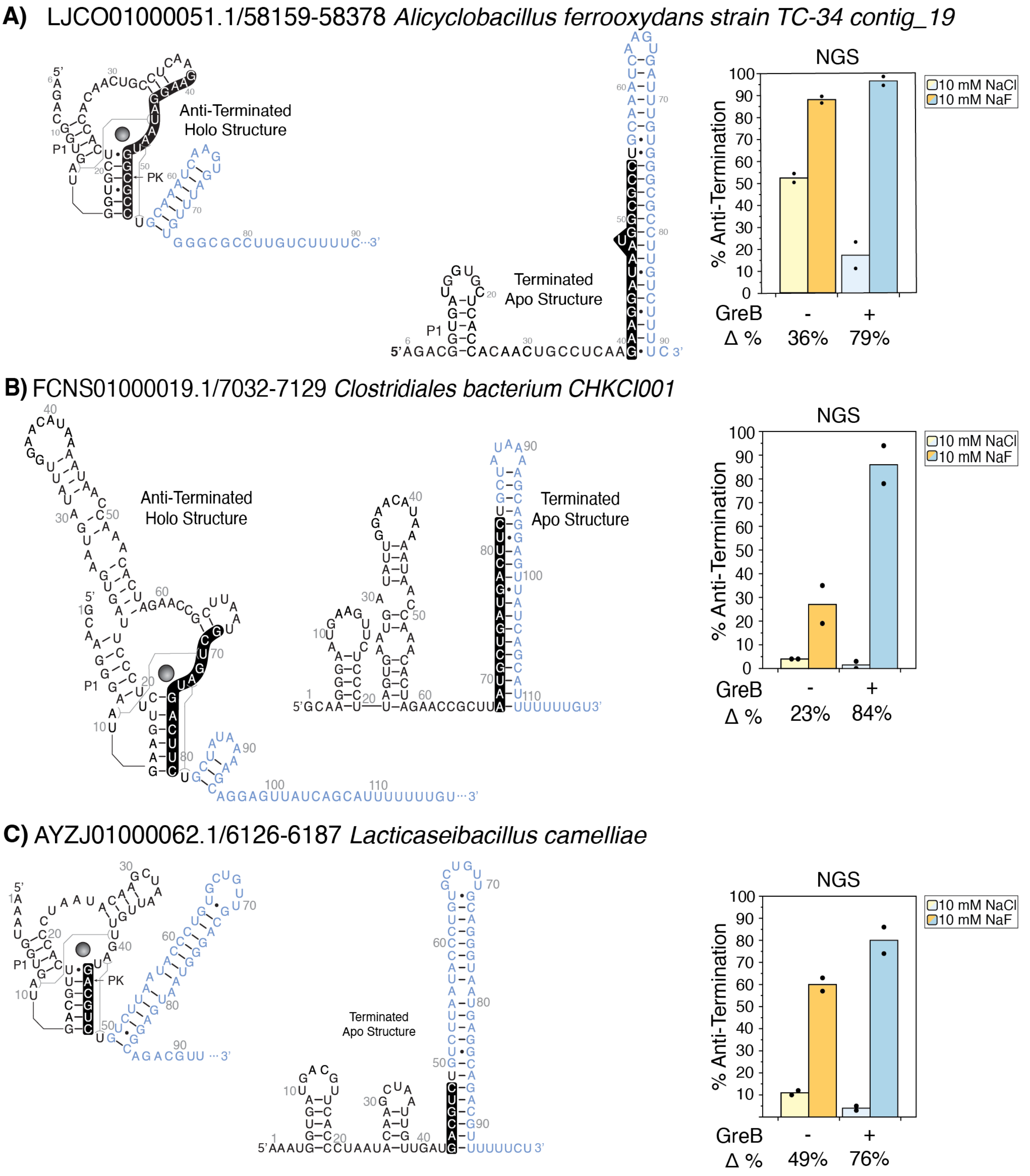

Figure S16. Fluoride riboswitch structures and NGS assay results without or with GreB for Figure 4. Hypothesized structures for the riboswitch variants informed by the RNACentral database and RNAStructure. The bar graph depicts the percent (%) Anti-termination either without (0 μM, yellow) or with (1.2 μM, blue) the transcription elongation factor, GreB. **(A)** *Alicyclobacillus ferrooxydans* (*A. fe*; LJCO01000051.1/58159-58221, **(B)** *Clostridiales bacterium CHKCI001* (*C. ba CHKCI001*; FCNS01000019.1/7032-7129, **(C)** *Lacticaseibacillus camelliae* (*L. ca*, AYZJ01000062.1/6126-6187). The change (Δ) of % Anti-termination is written for each condition. Bars represent average % anti-termination with points plotted (N = 2). Annotated gels in Figure S20. Data in Supplemental Document A.

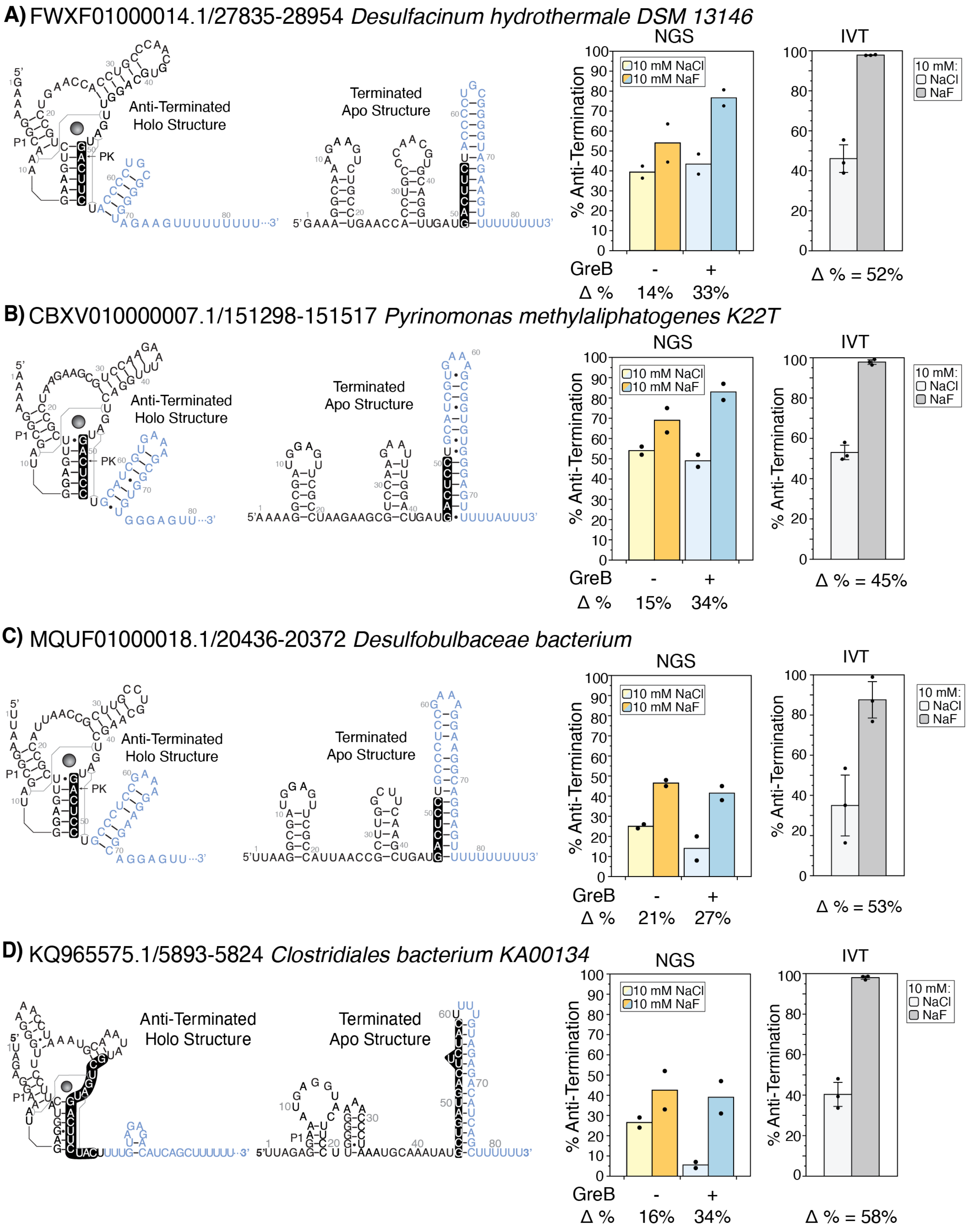

A) FWXF01000014.1/27835-28954 *Desulfacinum hydrothermale DSM 13146*
Anti-Terminated Holo Structure
Terminated Apo Structure
P1
PK
NGS
IVT
% Anti-Termination
10 mM NaCl
10 mM NaF
10 mM:
NaCl
NaF
GreB
-
+
Δ %
14%
33%
Δ % = 52%
B) CBXV010000007.1/151298-151517 *Pyrinomonas methylaliphatogenes K22T*
Anti-Terminated Holo Structure
Terminated Apo Structure
NGS
IVT
% Anti-Termination
10 mM NaCl
10 mM NaF
10 mM:
NaCl
NaF
GreB
-
+
Δ %
15%
34%
Δ % = 45%
C) MQUF01000018.1/20436-20372 *Desulfobulbaceae bacterium*
Anti-Terminated Holo Structure
Terminated Apo Structure
NGS
IVT
% Anti-Termination
10 mM NaCl
10 mM NaF
10 mM:
NaCl
NaF
GreB
-
+
Δ %
21%
27%
Δ % = 53%
D) KQ965575.1/5893-5824 *Clostridiales bacterium KA00134*
Anti-Terminated Holo Structure
Terminated Apo Structure
NGS
IVT
% Anti-Termination
10 mM NaCl
10 mM NaF
10 mM:
NaCl
NaF
GreB
-
+
Δ %
16%
34%
Δ % = 58%

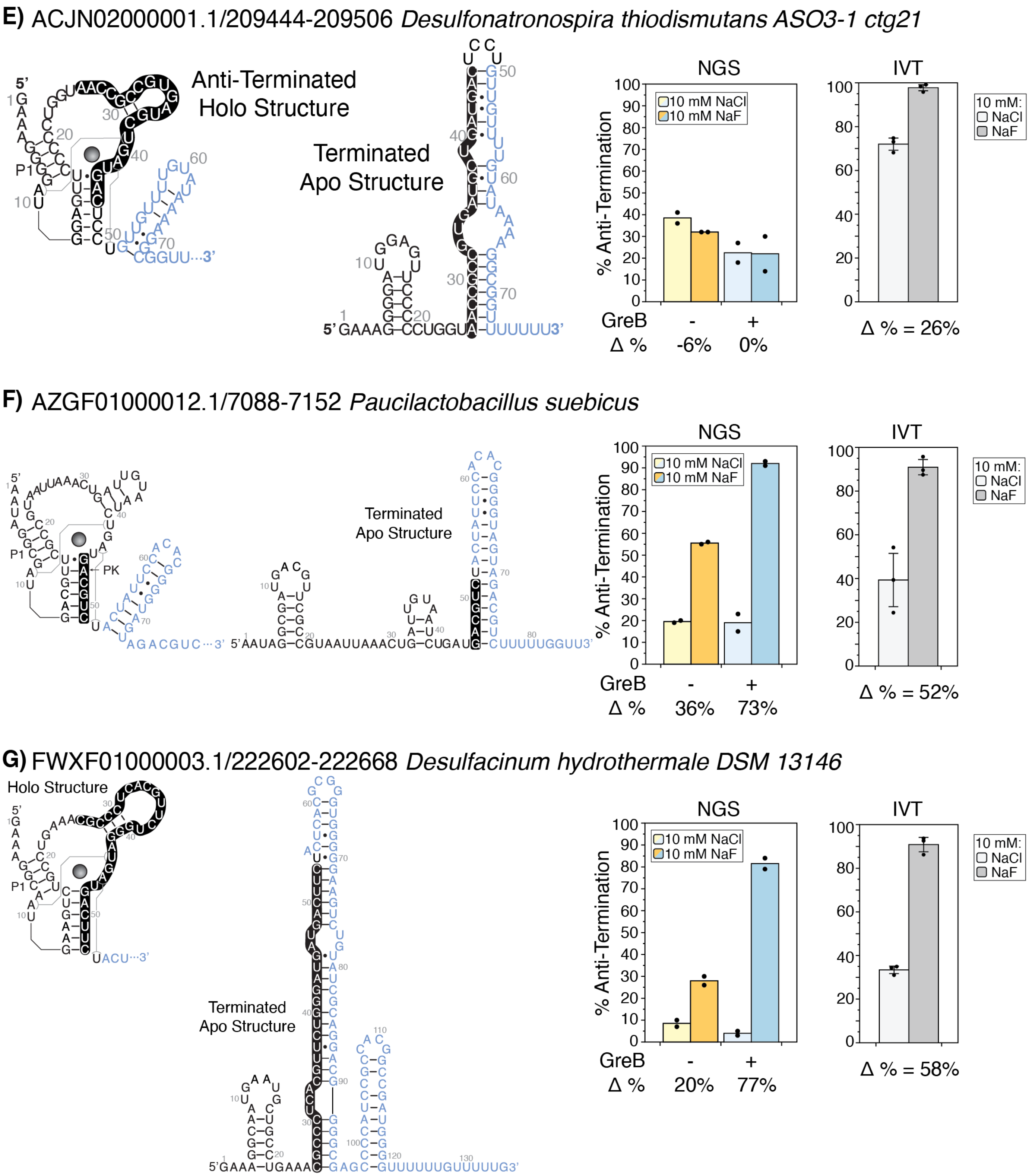

Figure S17. Fluoride riboswitch structures, NGS assay results without or with GreB, and IVT assay results with GreB for Figure 4E. Hypothesized structures for the riboswitch variants informed by the RNACentral database and RNAStructure. The blue bar graph depicts the percent (%) Anti-termination either without (0 µM) or with (1.2 µM) the transcription elongation factor, GreB. The change (Δ) of % Anti-termination is written for each condition. Bars represent average % anti-termination with error bars representing ± standard deviation (N = 2 for the high-throughput NGS assay, N = 3 for the low-throughput IVT assay). Annotated gels in Figure S21. Data in Supplemental Document A.

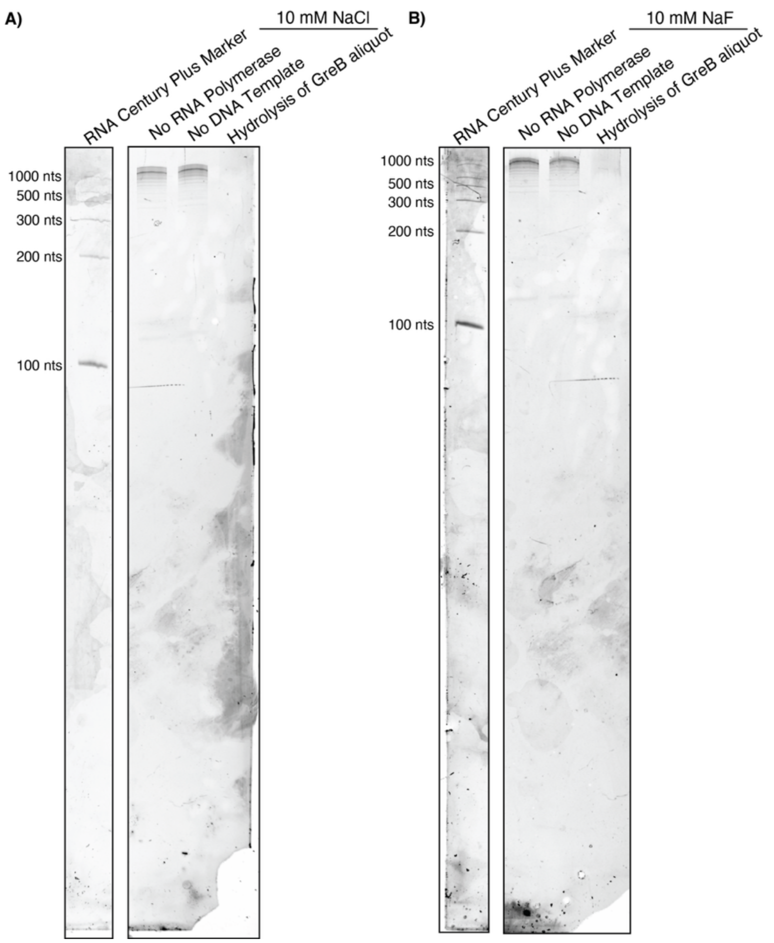

Figure S18. Annotated gels of GreB RNA co-precipitate. In the IVT assay gel output (Figure S19-S21), several bands can be seen that exceed the expected length for the input DNA template. We show these bands are an RNA co-precipitate that remained through the GreB purification process. The first lane, "No RNA Polymerase," underwent the full single-round IVT reaction but the RNAP was never added in the transcription reaction. The second lane, "No DNA Template," underwent the full single-round IVT reaction but no DNA template was added. The last lane, "Hydrolysis of GreB aliquot," was a protein sample input into the transcription reaction buffer with either 10 mM NaCl **(A)** or NaF **(B)** and 1 µL of 4 M NaOH added and boiled at 95 °C for 5 min. 2 µL of 1 M HCl was added to the sample and cleaned via ethanol precipitation to remove salts prior to gel run. The pellet was resuspended in 10 µL of water and then 10 µL of 2x RNA dye was added. The samples were run on a 10% denaturing gel and the raw gel images are provided in the "RAW GEL IMAGES" section.

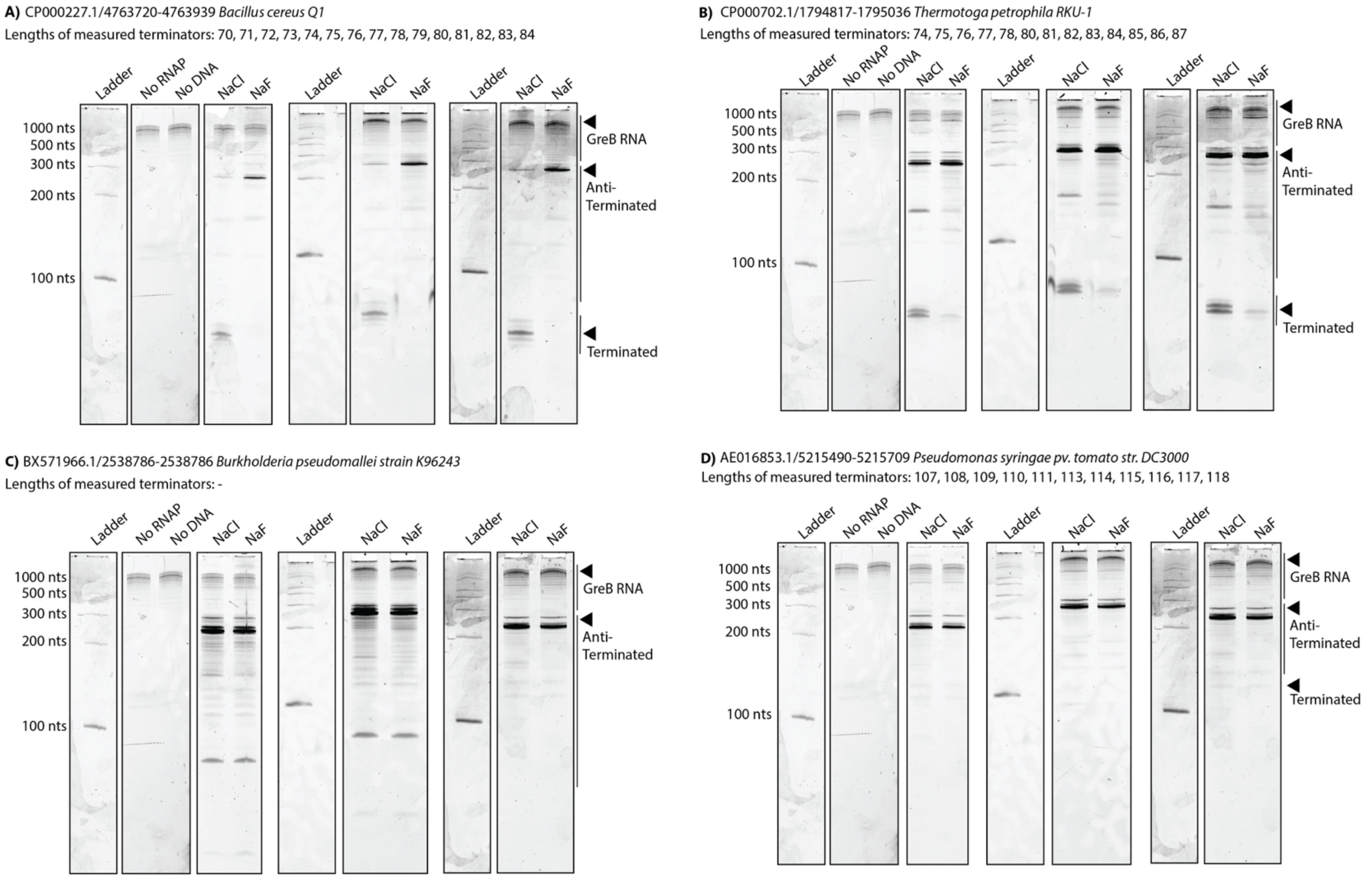

Figure S19. Annotated gel replicates of IVT data in Figure 3. RNA output from the low-throughput single-round IVT reaction with either 10 mM NaCl or NaF on a 10% denaturing gel in three biological replicates (N=3). IVT reactions were run for 5 min in the presence of 1.2 µM GreB, which resulted in excess bands from the RNA co-precipitant with the protein (Figure S18). The length of the terminated transcripts measured in the high-throughput assay are listed. The left most lane for each sample is the ladder (RNA Century Plus Marker) for each gel, then the GreB bands as a comparison, then the replicates, with the RNA band labeling on the right: **(A)** *Bacillus cereus* (accession ID: CP000227.1/4763720-4763779), **(B)** *Thermotoga petrophila* (accession ID: CP000702.1/1794817-1794880), **(C)** *Burkholderia pseudomallei* (accession ID: BX571966.1/2539005-2538939), and **(D)** *Pseudomonas syringae* (accession ID: AE016853.1/5215709-5215637). The raw gel images are provided in the “RAW GEL IMAGES” section.

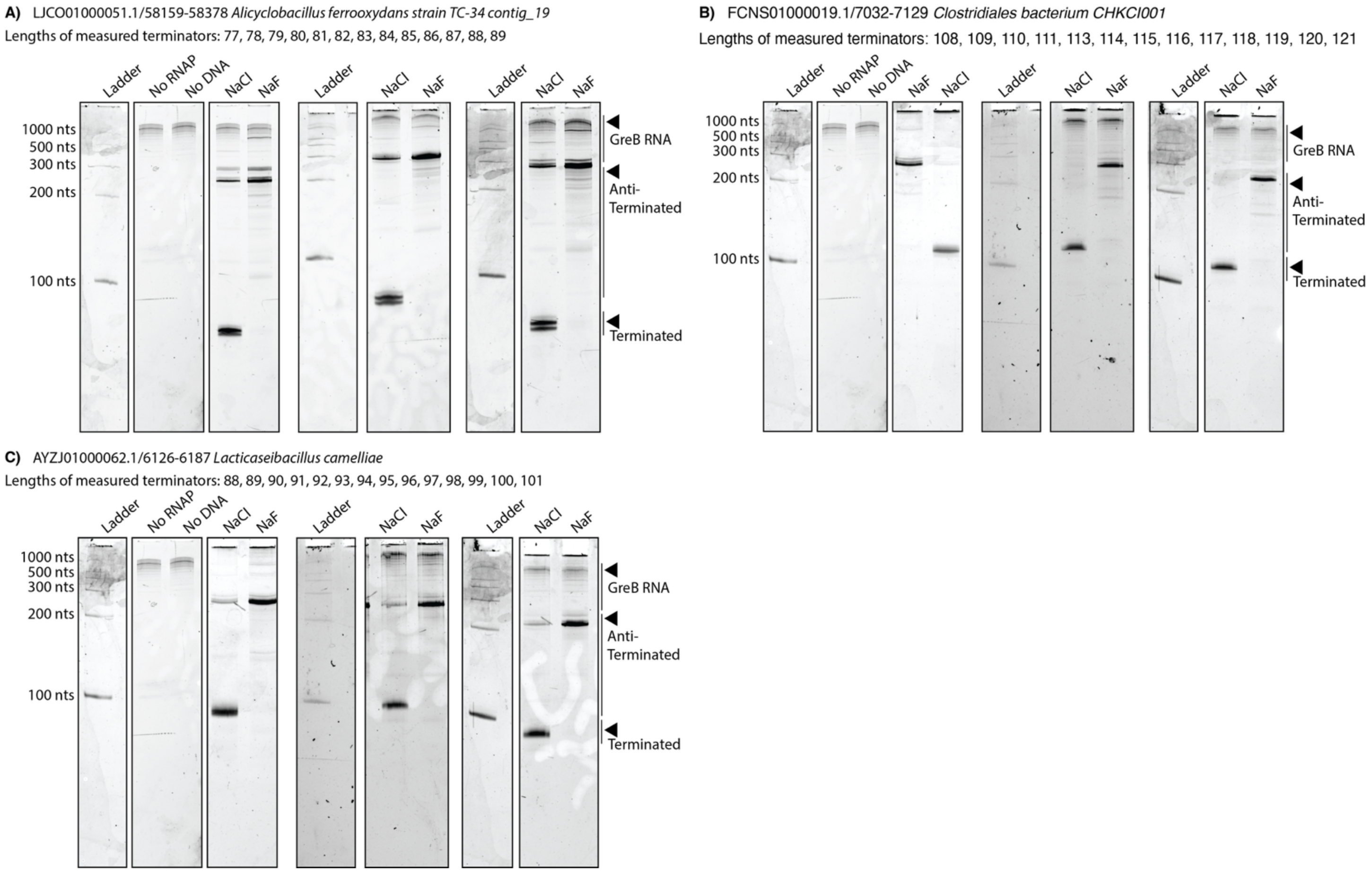

Figure S20. Annotated gel replicates of IVT data in Figure 4B-D. RNA output from the low-throughput single-round IVT reaction with either 10 mM NaCl or NaF on a 10% denaturing gel in three biological replicates (N=3). IVT reactions were run for 5 min in the presence of 1.2 µM GreB, which resulted in excess bands from the RNA co-precipitant with the protein (Figure S18). The length of the terminated transcripts measured in the high-throughput assay are listed. The left most lane for each sample is the ladder (RNA Century Plus Marker) for each gel, then the GreB bands as a comparison, then the replicates, with the RNA band labeling on the right. The raw gel images are provided in the "RAW GEL IMAGES" section.

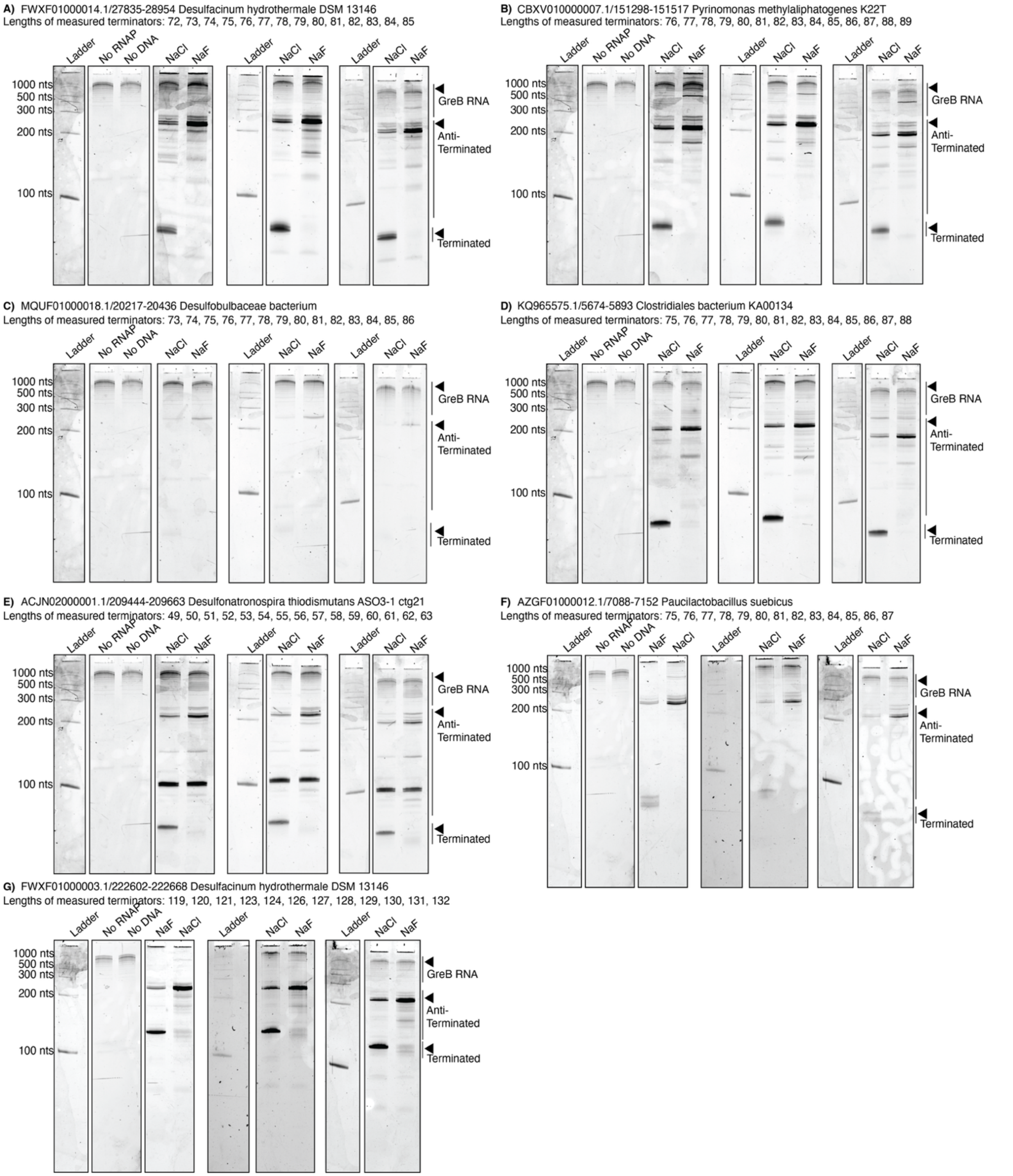

Figure S21. Annotated gel replicates of IVT data in Figure 4E. RNA output from the low-throughput single-round IVT reaction with either 10 mM NaCl or NaF on a 10% denaturing gel in three biological replicates (N=3). IVT reactions were run for 5 min in the presence of 1.2 µM GreB, which resulted in excess bands from the RNA co-precipitant with the protein (Figure S18). The length of the terminated transcripts measured in the high-throughput assay are listed. The left most lane for each sample is the ladder (RNA Century Plus Marker) for each gel, then the GreB bands as a comparison, then the replicates, with the RNA band labeling on the right. The raw gel images are provided in the “RAW GEL IMAGES” section.

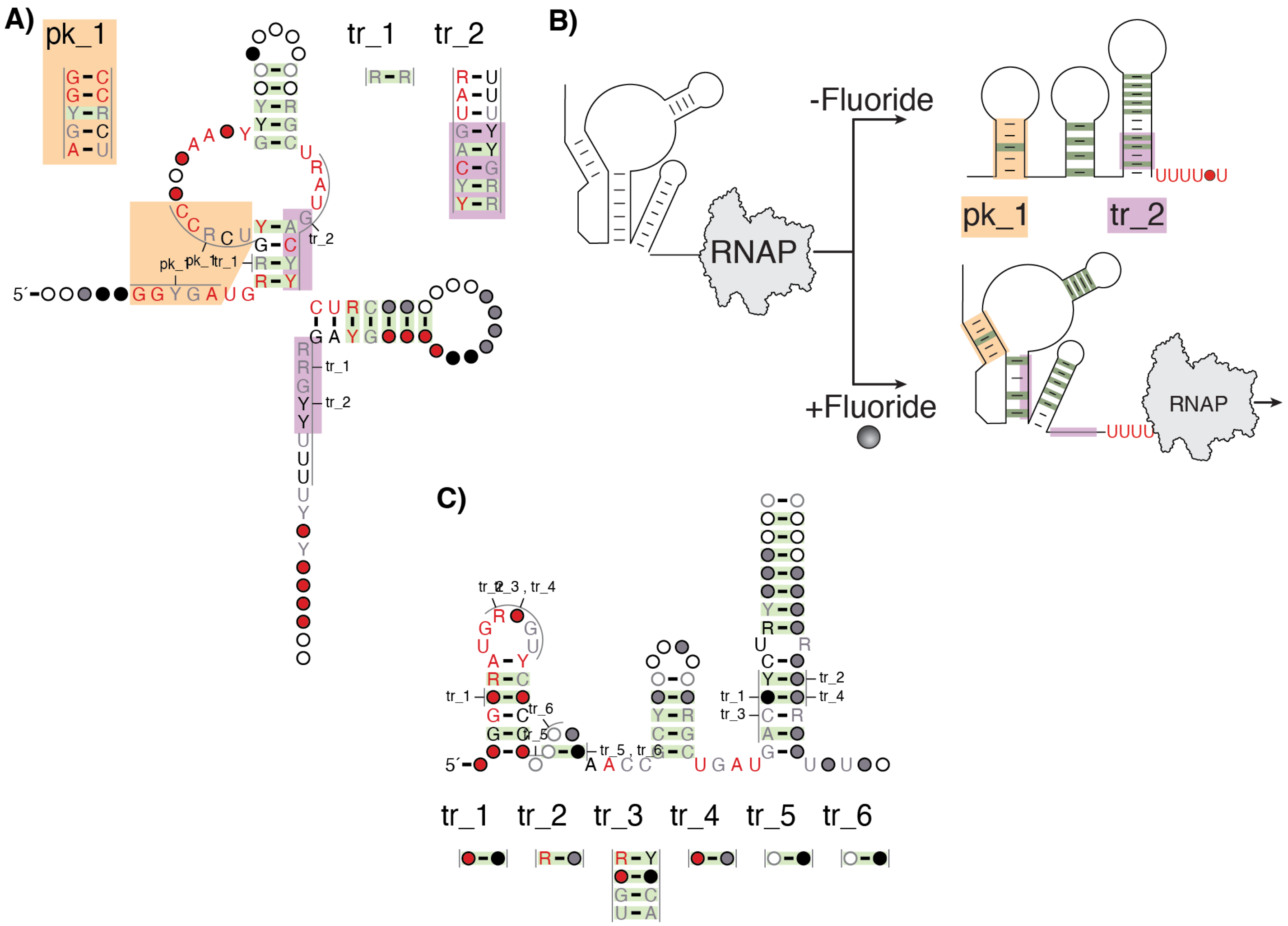

Figure S22. Raw R-scape outputs from running CaCoFold.**(2)** **(A)** Output from CaCoFold for fluoride riboswitch variants containing a predicted intrinsic terminator by ARNold. The P1 stem, labeled "pk_1", is highlighted in orange. The R-scape output shows the Holo alternative stems, PK, as part of the main structure and the Apo alternative stem, P4, as extended through "tr_2" (highlighted in purple). **(B)** Cartoon schematic of how (A) encodes both the apo (top) and holo (bottom) confirmations with key components highlighted (pk_1 as orange, tr_2 as purple). (**C)** Output from CaCoFold for variants measured to terminate with the NGS assay. The R-scape output shows the Apo alternative stem, P4, as part of the main structure and the Holo alternative stem, PK, as "tr_3". The raw output from CaCoFold is in Supplemental Document C.

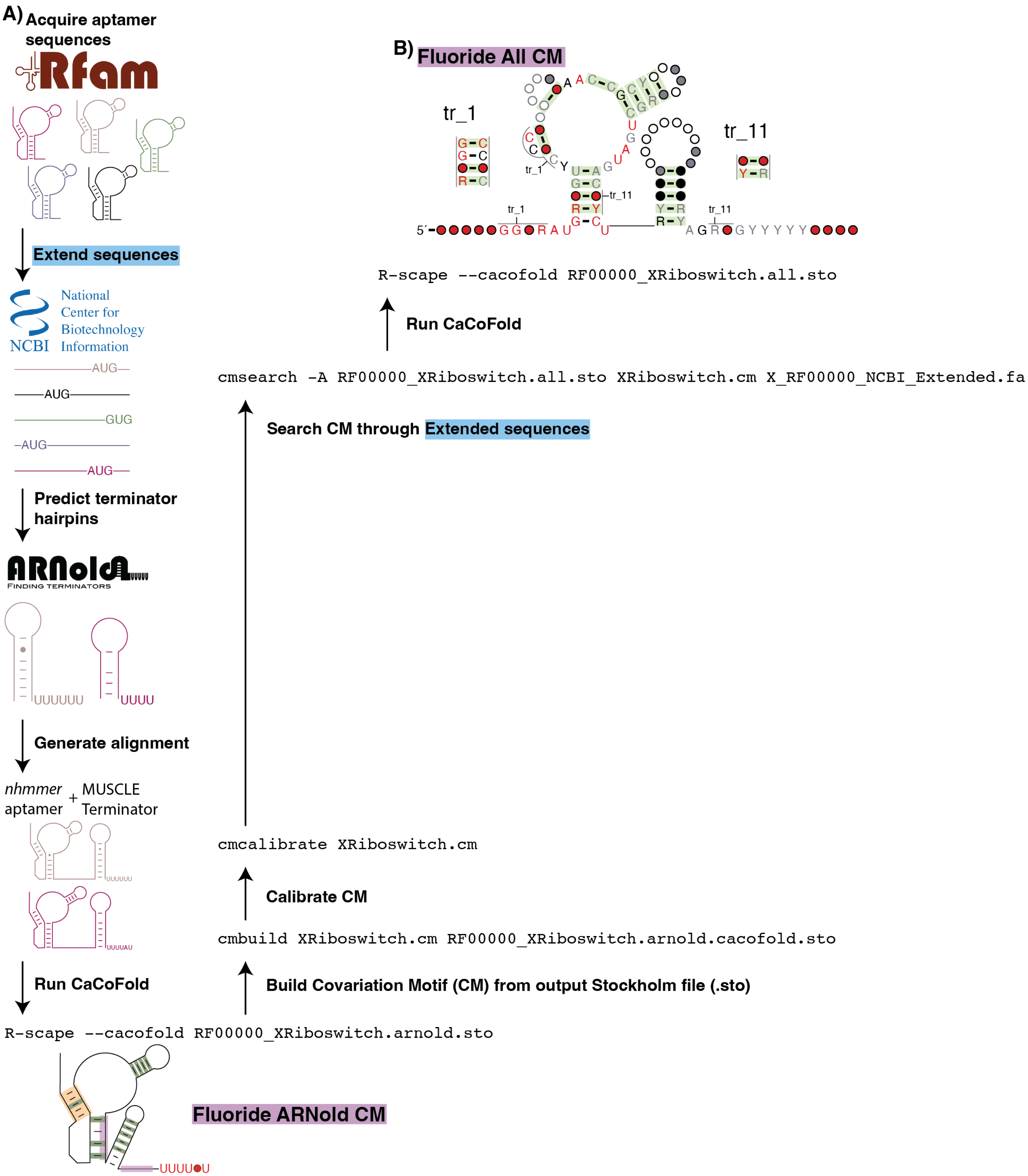

Figure S23. Searching all fluoride riboswitch variants for covariation motif analysis. **(A)** A schematic of the workflow described in the Methods and Materials: acquire deposited sequences on Rfam, extend the sequences through the NCBI, predict terminator hairpins, conduct alignment for the aptamer and terminator and combine, run CaCoFold, build the CaCoFold output into a Covariation Motif (CM), calibrate the CM, then search the sequences extended from the NCBI, re-run CaCoFold. **(B)** The CaCoFold structure output for the fluoride riboswitch after calibrating the predicted motif and searching all 1,901 sequences.

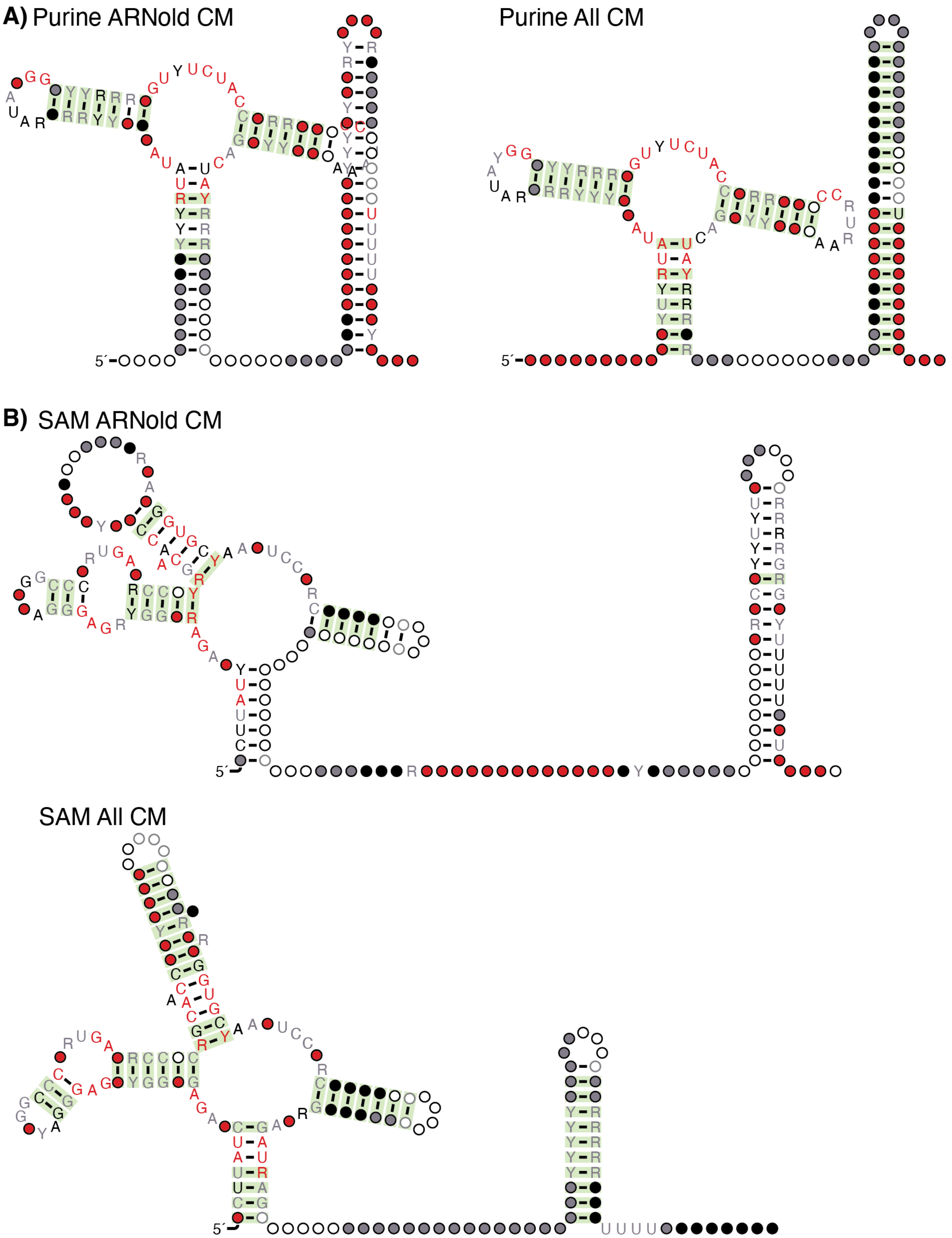

Figure S24. Covariation models of purine and SAM riboswitches. Covariation models were generated as in Figure 5 using the ARNold computational prediction to filter for transcriptional sequence variants. The models were then calibrated and searched for through all downloaded sequences of each riboswitch class (Figure S23). **(A)** The Purine (RF00167) riboswitch CaCoFold output after ARNold filtering (left) and after model calibration and searching all sequences (right). **(B)** The SAM (RF00162) riboswitch CaCoFold output after ARNold filtering (top) and after model calibration and searching all sequences (bottom). Green highlighting on base pairs denotes evolutionarily significance in base pair covariation, and nucleotides are denoted as circle or specific letters with colors to signify conservation: red = > 97%, black = 90-97%, grey = 75-90%, and white = 50-75%; R = purines: A or G. Y = pyrimidines: U or C; P1 = Pairing Element. The raw output from CaCoFold is in Supplemental Document C.

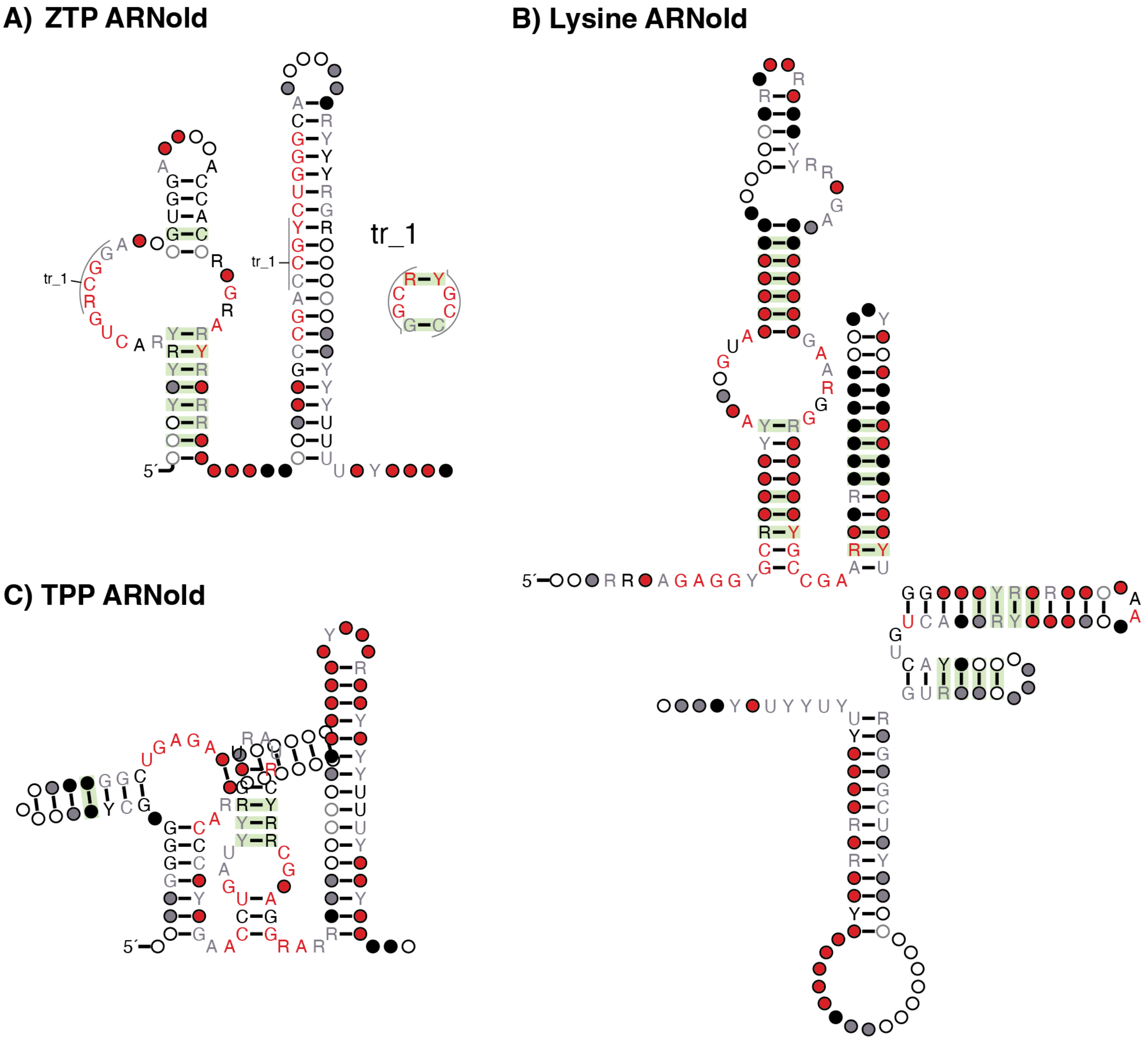

Figure S25. The CaCoFold outputs for the ZTP (RF01750), Lysine (RF00168), and TPP (RF00059) riboswitches at the stage of collecting terminating sequences from ARNold. Green highlighting on base pairs denotes evolutionarily significance in base pair covariation, and nucleotides are denoted as circle or specific letters with colors to signify conservation: red = > 97%, black = 90-97%, grey = 75-90%, and white = 50-75%; R = purines: A or G. Y = pyrimidines: U or C; P1 = Pairing Element. The raw output from CaCoFold is in Supplemental Document C.

**A)** glmS ARNold

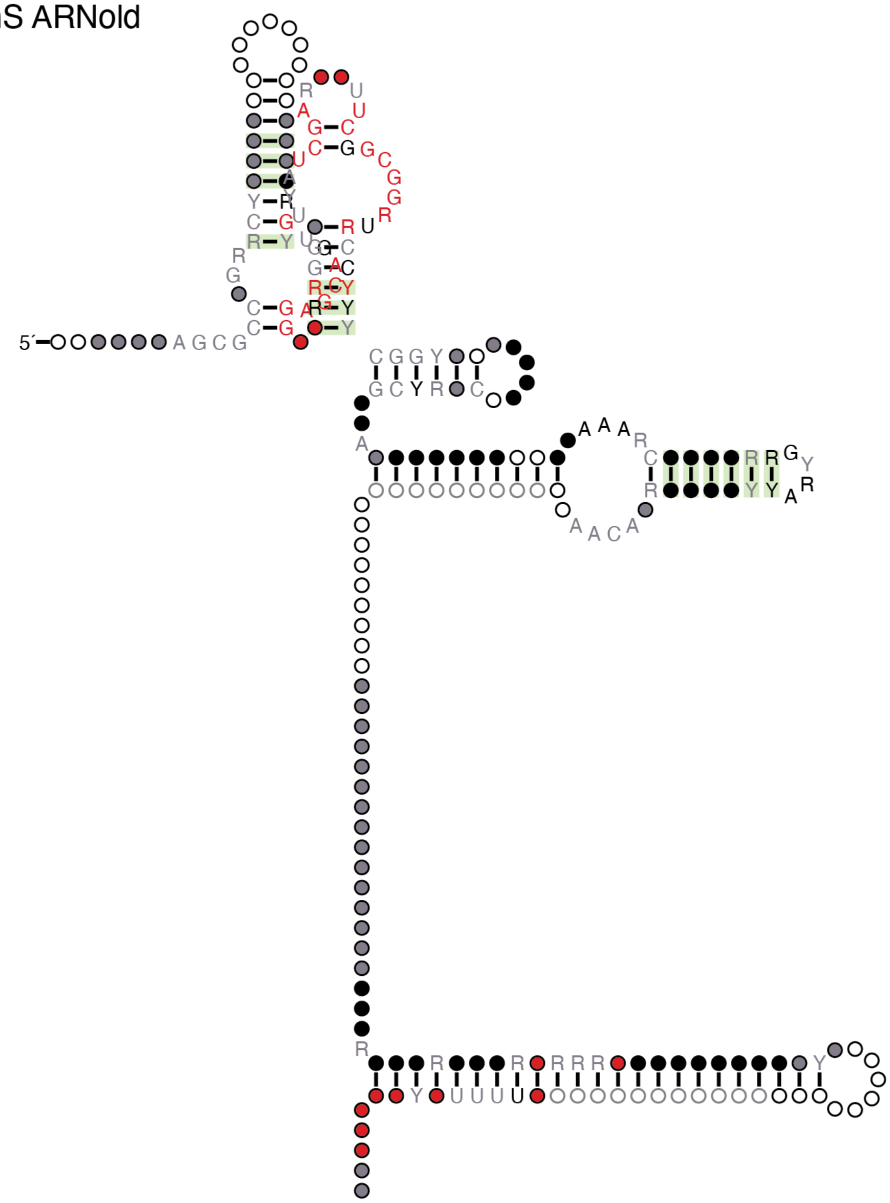

**B)** glmS CM

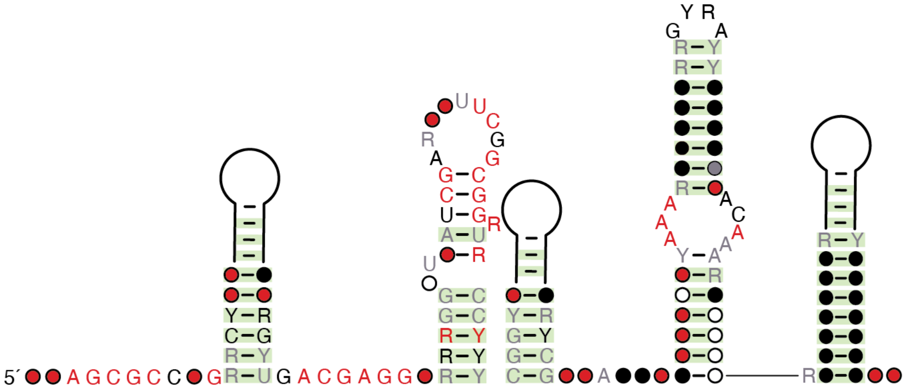

Figure S26. The CaCoFold outputs for the glmS (RF00083) riboswitch. Green highlighting on base pairs denotes evolutionarily significance in base pair covariation, and nucleotides are denoted as circle or specific letters with colors to signify conservation: red = > 97%, black = 90-97%, grey = 75-90%, and white = 50-75%; R = purines: A or G. Y = pyrimidines: U or C; P1 = Pairing Element.The raw output from CaCoFold is in Supplemental Document C.

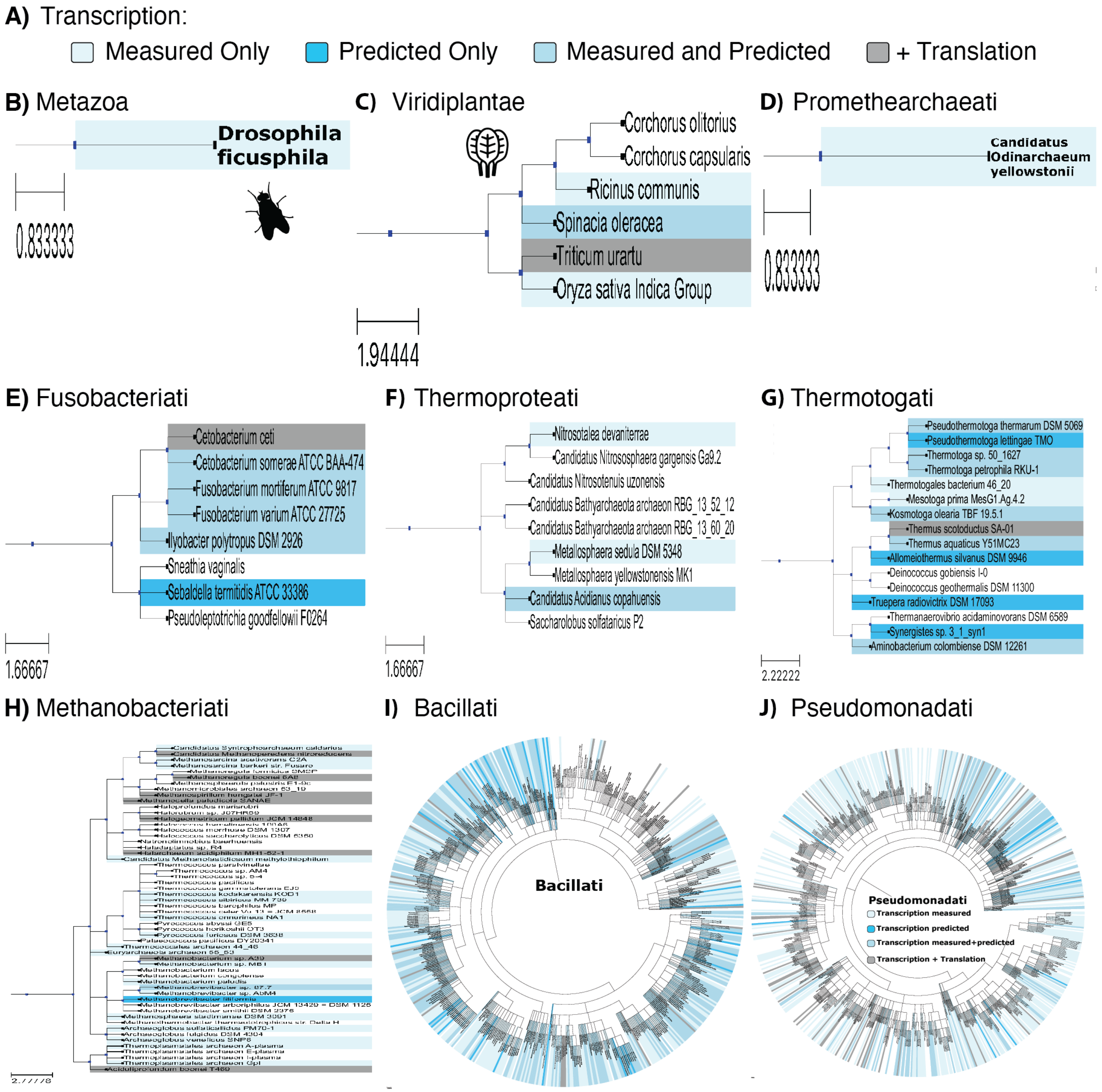

Figure S27. Kingdom phylogenetic trees. Phylogenetic trees of the taxa from the fluoride riboswitch variants split by kingdom. **(A)** Legend of the background color to designate the assay results for the variant in the species: transcription was only measured in the NGS assay (light blue), transcription was only predicted by ARNold (blue), transcription was both measured and predicted (sky blue), or a species had multiple variants that were either transcriptional or translational (grey). The phylogenetic trees made from this list of species include: Metazoa **(B)**, Vidiplantae **(C)** Promethearchaeati **(D)**, Fusobacteriati **(E)**, Thermoproteati **(F)**, Thermotogati **(G)**, Methanobacteriati **(H)**, Bacillati **(I)**, and Pseuomonadati **(J)**. Python packages used to generate these graphs: re, pandas, ete3 (NCBITaxa, Tree, TreeStyle, NodeStyle,TextFace), collections (defaultdict). Output trees in Supplemental Document D.

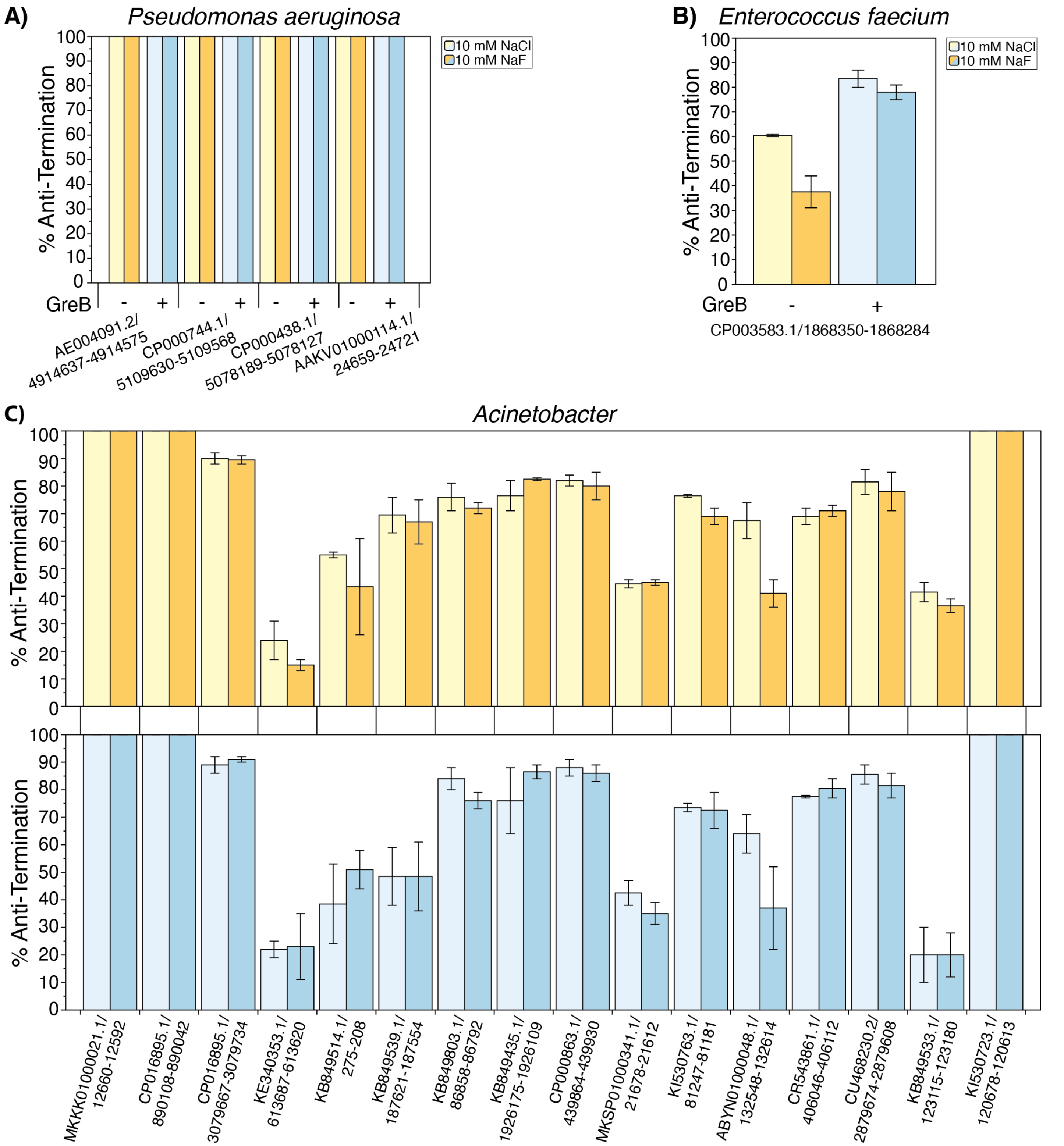

Figure S28. Antibiotic targets. **(3)** Results from high-throughput assay for fluoride riboswitch variants from bacterial species of antibiotic targeting interest: *Pseudomonas aeruginosa* **(A)**, *Enterococcus faecium*, considered a "serious threat" **(B)**, and *Acinetobacter* **(C)**. The data shown is from reactions without (yellow) or with 1.2 µM (blue) GreB and either 10 mM NaCl (light blue) or 10 mM NaF (blue).

### SUPPLEMENTAL TABLES

| Name | Use | Sequence |
|---|---|---|
| *E. coli* J23119 promoter | Transcription and oligo pool amplification | gcttccggcttgattctaaagatctttgacagctagctcagtcctaggtataatactagt |
| 3' primer site | Oligo pool amplification | gggcacaaattttctgtccg |

**Table 1: Sequences used in building the oligo pool.**

| Name | Use | Sequence | Ordering (IDT) |
|---|---|---|---|
| A | ssDNA library amplification (Forward) | gcttccggcttgattctaaagatc | PAGE purified |
| B | ssDNA library amplification (Reverse) | cggacagaaaatttgtgccc | PAGE purified |
| C | Sequence oligo pool (Forward) | AATGATACGGCGACCACCGAGATCTA CACTCTTTCCCTACACGACGCTCTTCC GATCTNNNNgcttccggcttgattctaaagatc | PAGE purified |
| D | Sequence oligo pool (Reverse) | CAAGCAGAAGACGGCATACGAGATagg aatGTGACTGGAGTTCAGACGTGTGCTC TTCCGATCTNNNNcggacagaaaatttgtgccc | PAGE purified |
| E | 3WJB amplification | CATTACTCGCATCCATTCTCAGGCTGTCTCGTCTCGTCTC | Standard Desalting |
| F | 3WJB amplification | GCTTGGATTCTGCGTTTGTTTCCGTCTACGAACTCCCAGC | Standard Desalting |
| G | Linker | /5Phos/rCrUrGrArCrUrCrGrGrGrCrArCrCrArArGrGrA/3ddC/ | Standard Desalting |
| H | RT Primer | /5BiosG/GTCCTTGGTGCCCGAGT | Standard Desalting |
| I | SS2.0 Dumbbell | /5Phos/TGAAGAGCCTAGTCGCTGTTCANNNNNNCTGCC CATAGAG/3SpC3/ | PAGE purified |
| J | Index | CAAGCAGAAGACGGCATACGAGAT[Table_3]GTGACTG GAGTTCAGACGTGTGCTCTTCCGATCTTGAACAGCGAC TAGGCTCTTCA | PAGE purified |
| K | Selection, RRRY (NaCl) | CTTTCCCTACACGACGCTCTTCCGATCTRRRYGTCCTT GGTGCCCGAG*T*C*A*G | Standard Desalting |
| L | Selection, YYYR (NaCl) | CTTTCCCTACACGACGCTCTTCCGATCTYYYRGTCCTT GGTGCCCGAG*T*C*A*G | Standard Desalting |
| M | TruSeq Universal Adapter | AATGATACGGCGACCACCGAGATCTACACTCTTTCCC TACACGACGCTCTTCCGATCT | Standard Desalting |

**Table 2: Oligos used for IVT dsDNA template generation** (A, B), NGS (C, D, J, K, L, M), GreB dsDNA template generation (E, F), and NGS library prep (G, H, I).

| Index Number | Sequence |
|---|---|
| i9 | CTGATC |
| i10 | AAGCTA |
| i22 | CGTACG |
| i23 | CCACTC |
| i1 | CGTGAT |
| i2 | ACATCG |
| i3 | TGACAT |
| i4 | GGACGG |

**Table 3: Index list.**

| Parameter | Average |
|---|---|
| ΔG | -10.18 kcal/mol |
| Terminator Length | 41 nts |
| Stem Lenght | 10 nts |
| Loop Length | 15 nts |
| Riboswitch Length | 97 nts |
| Terminator Start Position | 57 nts |

**Table 4: ARNold averages for the terminator results.**

| Samples | Sample Codes | r (Terminated Reads) | r (Anti-Terminated reads) |
|---|---|---|---|
| Rep 1 vs 2: NaCl, No GreB | LMH_1 vs LMH_6 | 0.97324 | 0.986 |
| Rep 1 vs 2: NaF, No GreB | LMH_2 vs LMH_7 | 0.98734 | 0.99762 |
| Rep 1 vs 2: NaCl, Yes GreB | LMH_3 vs LMH_8 | 0.956272 | 0.992258 |
| Rep 1 vs 2: NaF, Yes GreB | LMH_4 vs LMH_9 | 0.982996 | 0.989018 |

**Table 5: Pearson correlation coefficient (r) between samples as calculated in Excel (=PEARSON([array1],[array2])).**

| Code | Species | Accession ID | SI Figures |
|---|---|---|---|
| *A. fe* | *Alicyclobacillus ferrooxydans* | LJCO01000051.1/58159-58378 | Figure S16A |
| *B. ce* | *Bacillus cereus* | CP000227.1/4763720-4763779 | Figure S15B |
| *B. ps* | *Burkholderia pseudomallei* | BX571966.1/2539005-2538939 | Figure S15C |
| *C. ba* | *Clostridiales bacterium KA00134* | KQ965575.1/5893-5824 | Figure S17D |
| *C. ba CHKCI001* | *Clostridiales bacterium CHKCI001* | FCNS01000019.1/7032-7129 | Figure S16B |
| *D. ba* | *Desulfobulbaceae bacterium DB1* | MQUF01000018.1/20436-20372 | Figure S17C |
| *D. hy*[1] | *Desulfacinum hydrothermale* | FWXF01000014.1/27835-28954 | Figure S17A |
| *D. hy*[2] | *Desulfacinum hydrothermale* | FWXF01000003.1/222602-222668 | Figure S17G |
| *D. thio* | *Desulfonatronospira thiodismutans* | ACJN02000001.1/209444-209663 | Figure S17E |
| *L. ca* | *Lacticaseibacillus camelliae* | AYZJ01000062.1/6126-6187 | Figure S16C |
| *P. sy* | *Pseudomonas syringae* | AE016853.1/5215709-5215637 | Figure S15D |
| *P. me* | *Pyrinomonas methylaliphatogenes* | CBXV010000007.1/151298-151517 | Figure S17B |
| *P. su* | *Paucilactobacillus suebicus* | AZGF01000012.1/7088-7152 | Figure S17F |
| *T. pe* | *Thermotoga petrophila* | CP000702.1/1794817-1794880 | Figure S15A |

**Table 6: Key to Figure 4E.** Specie's three-letter code to their full name, NCBI accession ID and fluoride aptamer position, and the corresponding SI figure with predicted structure and complete NGS or IVT data.

**RAW GEL IMAGES**

The following show the uncrossed, unprocessed urea-PAGE gel images from Figures S2A, S18-S21.

Pre-sequencing gel image, Figure S2A

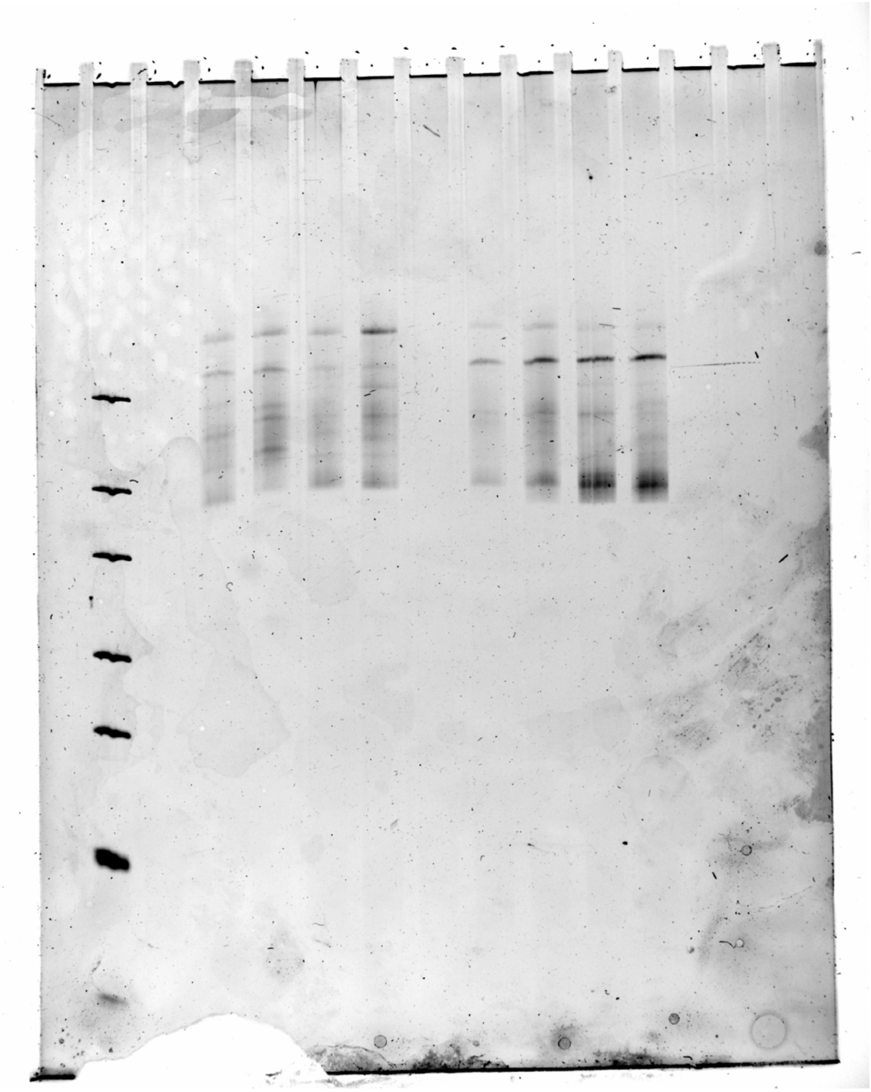

From left to right:
2. GeneRuler Ultra Low Range DNA Ladder (35, 50, 75, 100, 150, 200, and 300 bases) (Thermo Scientific, SM1213)
4. dsDNA products generated for Next-Generation Sequencing from an *in vitro* transcription reaction with a library of fluoride riboswitches in the presence of 10 mM NaCl (LMH_1)
5. dsDNA products generated for Next-Generation Sequencing from an *in vitro* transcription reaction with a library of fluoride riboswitches in the presence of 10 mM NaF (LMH_2)
6. dsDNA products generated for Next-Generation Sequencing from an *in vitro* transcription reaction with a library of fluoride riboswitches in the presence of 10 mM NaCl and 1.2 µM GreB (LMH_3)
7. dsDNA products generated for Next-Generation Sequencing from an *in vitro* transcription reaction with a library of fluoride riboswitches in the presence of 10 mM NaF and 1.2 µM GreB (LMH_4)
9. dsDNA products generated for Next-Generation Sequencing from an *in vitro* transcription reaction with a library of fluoride riboswitches in the presence of 10 mM NaCl (LMH_6)
10. dsDNA products generated for Next-Generation Sequencing from an *in vitro* transcription reaction with a library of fluoride riboswitches in the presence of 10 mM NaF (LMH_7)
11. dsDNA products generated for Next-Generation Sequencing from an *in vitro* transcription reaction with a library of fluoride riboswitches in the presence of 10 mM NaCl and 1.2 µM GreB (LMH_8)
12. dsDNA products generated for Next-Generation Sequencing from an *in vitro* transcription reaction with a library of fluoride riboswitches in the presence of 10 mM NaF and 1.2 µM GreB (LMH_9)

*In vitro* transcription RNA products, Figure S18, S19, S20

Replicate 1

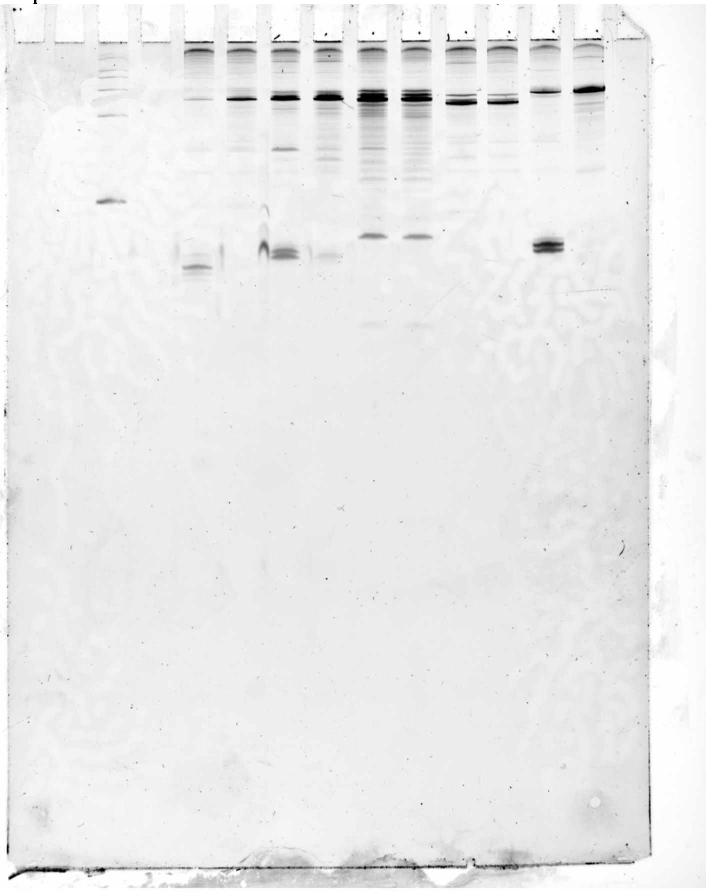

Replicate 2

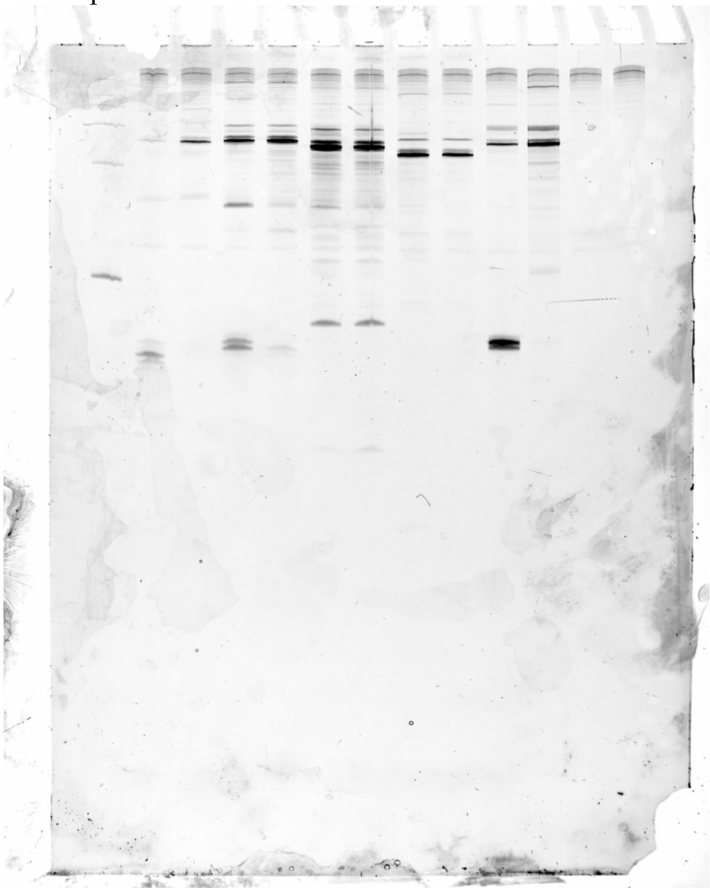

Replicate 3

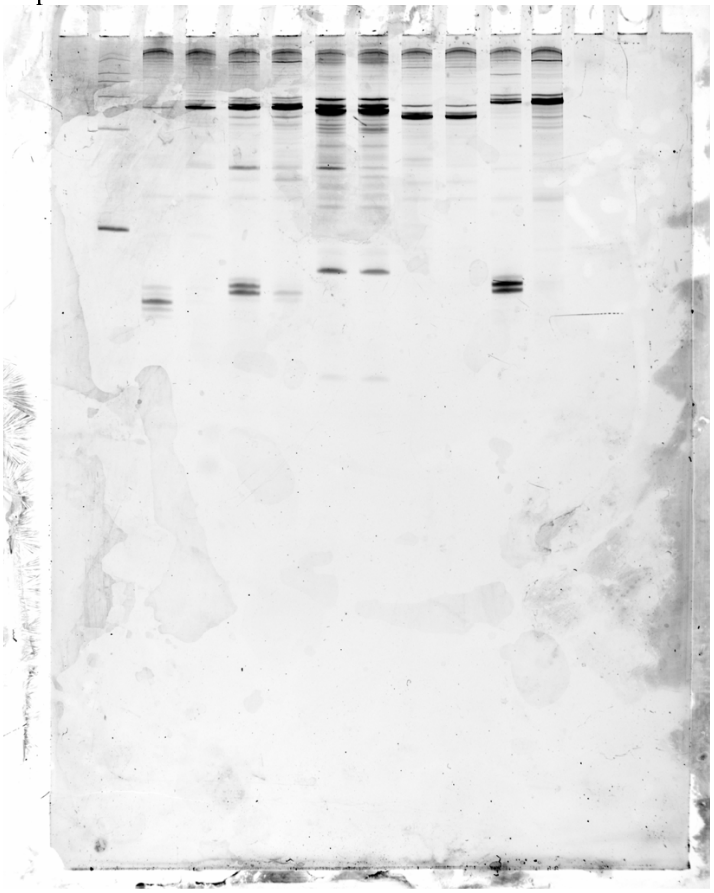

From left to right:
2. ssRNA Ladder (100, 200, 300, 400, 500, 750, and 1000 bases) (Invitrogen, cat. no. AM7145)
3-12. In vitro transcript products generated from E. coli RNAp during a single-round of transcription with 1.2 μM GreB and:
3. 10 mM NaCl, DNA template = CP000227.1/4763720-4763779
4. 10 mM NaF, DNA template = CP000227.1/4763720-4763779
5. 10 mM NaCl, DNA template = CP000702.1/1794817-1794880
6. 10 mM NaF, DNA template = CP000702.1/1794817-1794880
7. 10 mM NaCl, DNA template = BX571966.1/2539005-2538939
8. 10 mM NaF, DNA template = BX571966.1/2539005-2538939
9. 10 mM NaCl, DNA template = AE016853.1/5215709-5215637
10. 10 mM NaF, DNA template = AE016853.1/5215709-5215637
11. 10 mM NaCl, DNA template = LJCO01000051.1/58159-58221
12. 10 mM NaF, DNA template = LJCO01000051.1/58159-58221

Replicate 2:
13. No RNA Polymerase control with 10 mM NaCl
14. No DNA template control with 10 mM NaCl
15. GreB protein aliquot hydrolyzed with NaOH and boiled prior to gel loading with 10 mM NaCl

*In vitro* transcription RNA products, Figure S20, S21

Replicate 1

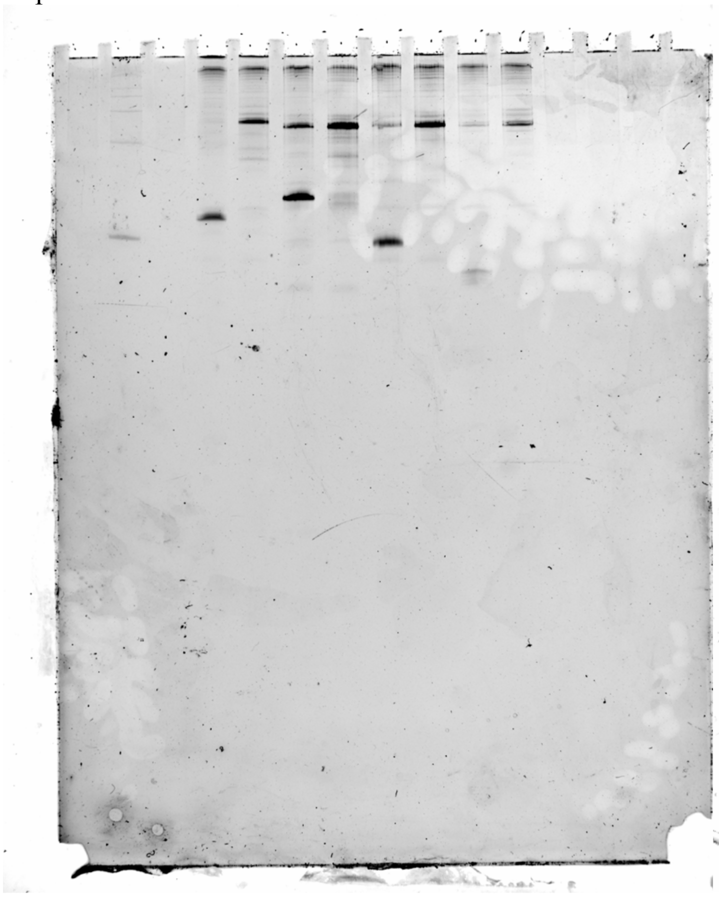

Replicate 2

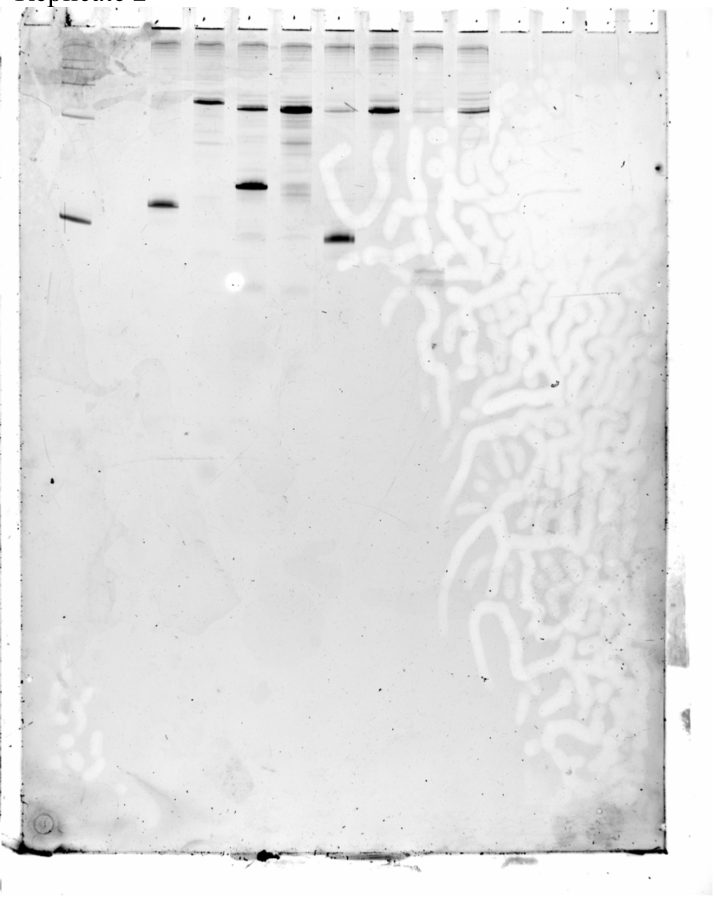

Replicate 3

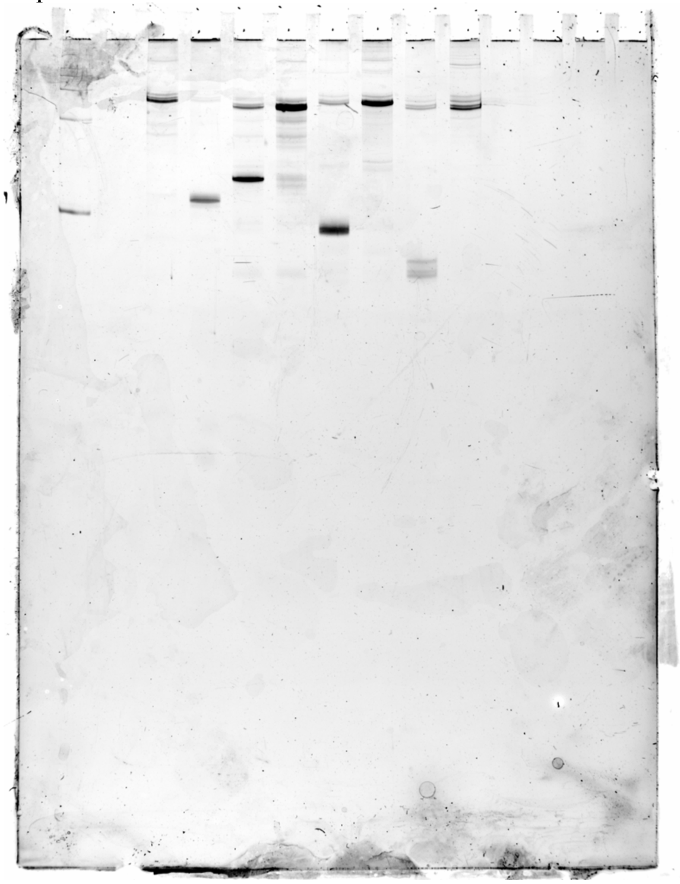

From left to right:

2. ssRNA Ladder (100, 200, 300, 400, 500, 750, and 1000 bases) (Invitrogen, cat. no. AM7145)

4-11. In vitro transcript products generated from E. coli RNAp during a single-round of transcription with 1.2 μM GreB and:

4. 10 mM NaCl, DNA template = FCNS01000019.1/7032-7129
5. 10 mM NaF, DNA template = FCNS01000019.1/7032-7129
6. 10 mM NaCl, DNA template = FWXF01000003.1/222602-222668
7. 10 mM NaF, DNA template = FWXF01000003.1/222602-222668
8. 10 mM NaCl, DNA template = AYZJ01000062.1/6126-6187
9. 10 mM NaF, DNA template = AYZJ01000062.1/6126-6187
10. 10 mM NaCl, DNA template = AZGF01000012.1/7088-7152
11. 10 mM NaF, DNA template = AZGF01000012.1/7088-7152

*Replicate 3:

4. 10 mM NaF, DNA template = FCNS01000019.1/7032-7129
5. 10 mM NaCl, DNA template = FCNS01000019.1/7032-7129

*In vitro* transcription RNA products, Figure S21

Replicate 1

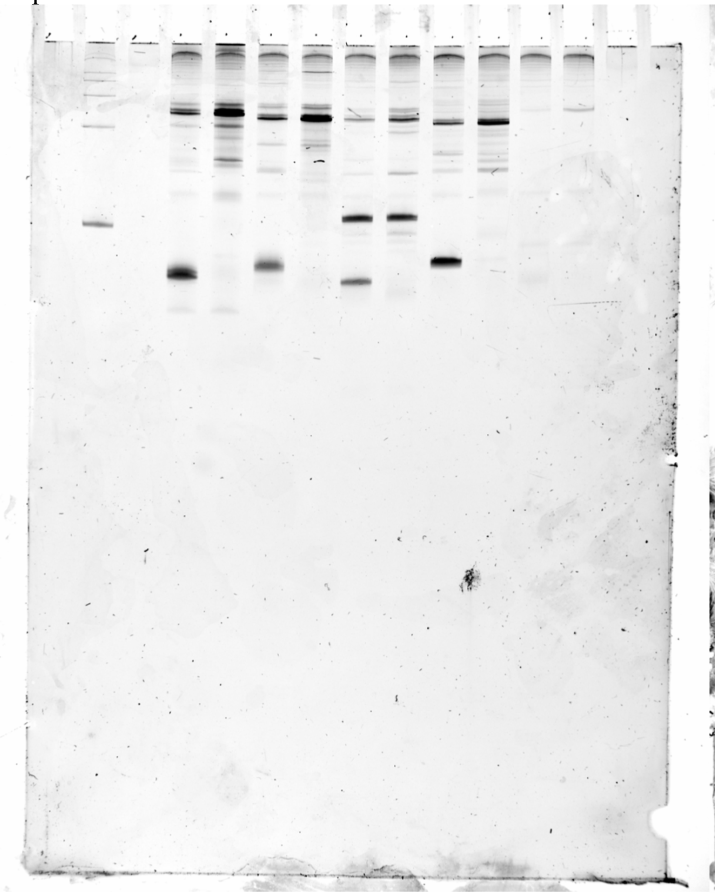

Replicate 2

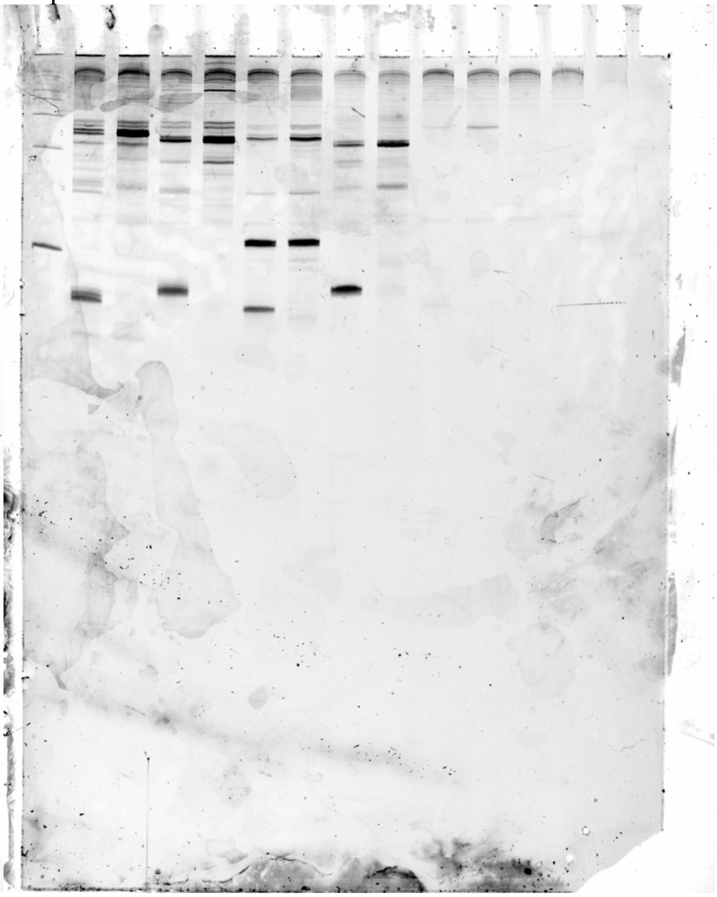

Replicate 3

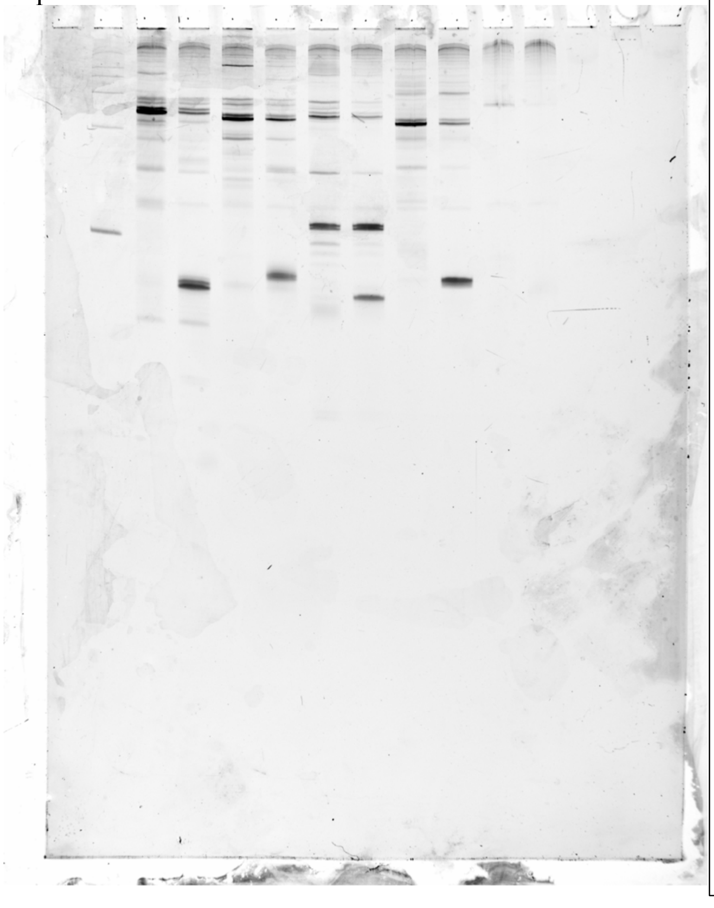

From left to right for Replicate 1 and 2:

2. ssRNA Ladder (100, 200, 300, 400, 500, 750, and 1000 bases) (Invitrogen, cat. no. AM7145)

3-12. In vitro transcript products generated from E. coli RNAp during a single-round of transcription with 1.2 µM GreB and:

3. 10 mM NaCl, DNA template = FWXF01000014.1/27835-27903
4. 10 mM NaF, DNA template = FWXF01000014.1/27835-27903
5. 10 mM NaCl, DNA template = CBXV010000007.1/151298-151365
6. 10 mM NaF, DNA template = CBXV010000007.1/151298-151365
7. 10 mM NaCl, DNA template = ACJN02000001.1/209444-209506
8. 10 mM NaF, DNA template = ACJN02000001.1/209444-209506
9. 10 mM NaCl, DNA template = KQ965575.1/5893-5824
10. 10 mM NaF, DNA template = KQ965575.1/5893-5824
11. 10 mM NaCl, DNA template = MQUF01000018.1/20436-20372
12. 10 mM NaF, DNA template = MQUF01000018.1/20436-20372

Replicate 2:

13. No RNA Polymerase control with 10 mM NaF
14. No DNA template control with 10 mM NaF
15. GreB protein aliquot hydrolyzed with NaOH and boiled prior to gel loading with 10 mM NaF

Replicate 3:

3. 10 mM NaF, DNA template = FWXF01000014.1/27835-27903
4. 10 mM NaCl, DNA template = FWXF01000014.1/27835-27903
5. 10 mM NaF, DNA template = CBXV010000007.1/151298-151365
6. 10 mM NaCl, DNA template = CBXV010000007.1/151298-151365
7. 10 mM NaF, DNA template = ACJN02000001.1/209444-209506
8. 10 mM NaCl, DNA template = ACJN02000001.1/209444-209506
9. 10 mM NaF, DNA template = KQ965575.1/5893-5824
10. 10 mM NaCl, DNA template = KQ965575.1/5893-5824
11. 10 mM NaF, DNA template = MQUF01000018.1/20436-20372
12. 10 mM NaCl, DNA template = MQUF01000018.1/20436-20372
