## Supplementary material for "High-throughput functional profiling and evolutionary covariation analysis of entire riboswitch sequences": Unedited output from R2R with Rscape.

### RF01750_ZTPRiboswitch.arnold.cacofold

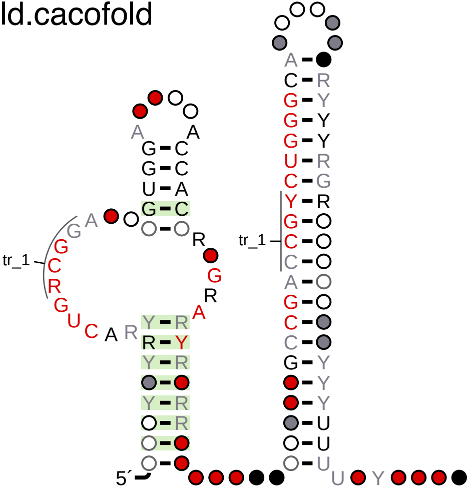


tr_1

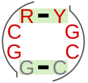

### RF01750_ZTPRiboswitch.all.cacofold

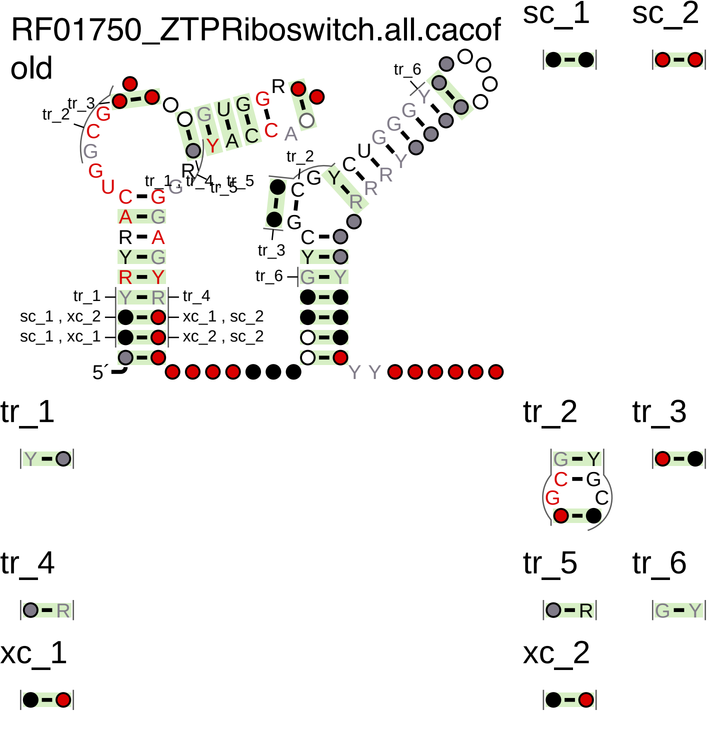

### RF00168_LysineRiboswitch.arnold.cacofold

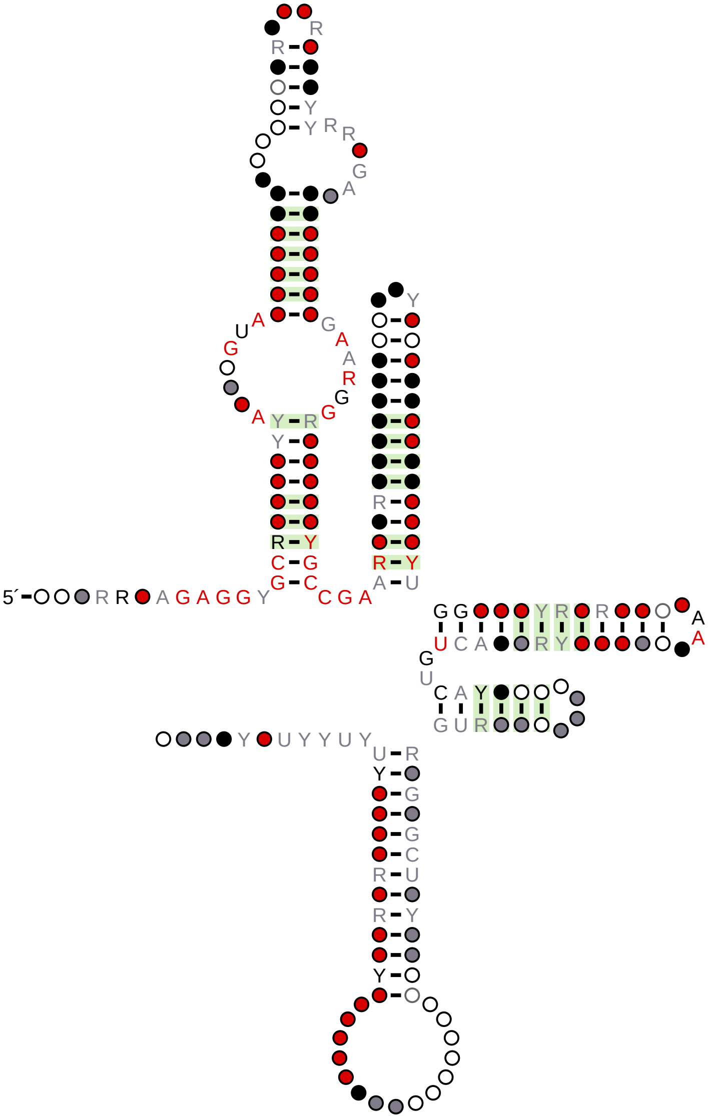

RF00168_LysineRiboswitch.all.cacofold

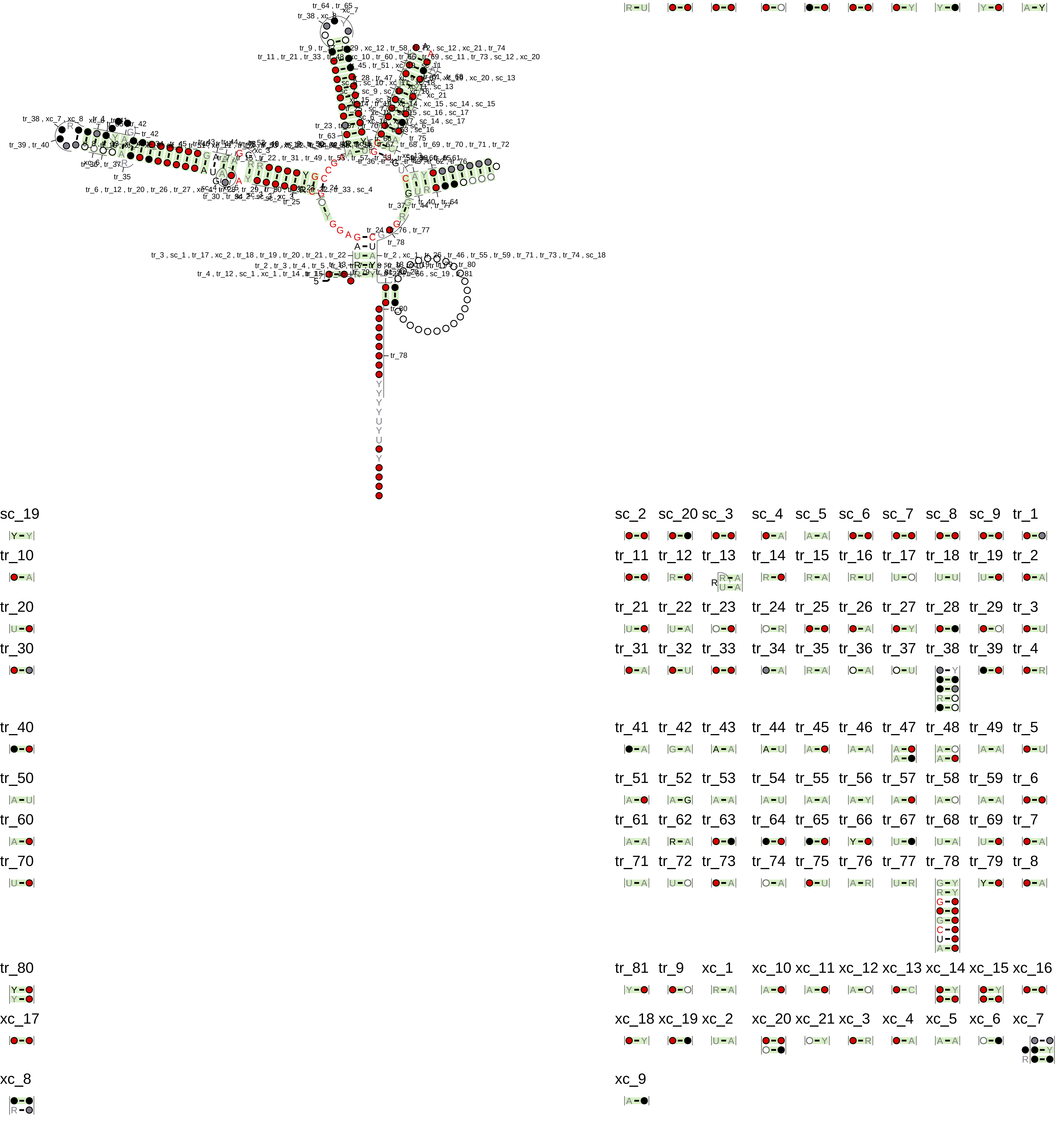

RF00059_TPPRiboswitch.arnold
.cacofold

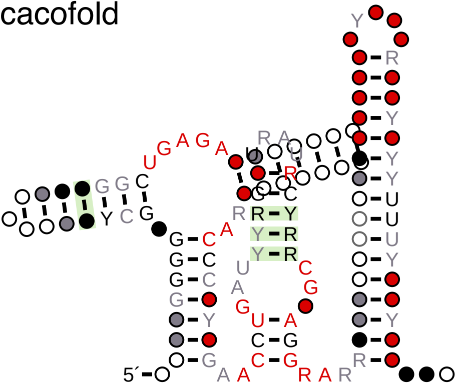

### RF00059_TPPRiboswitch.all.cacofold

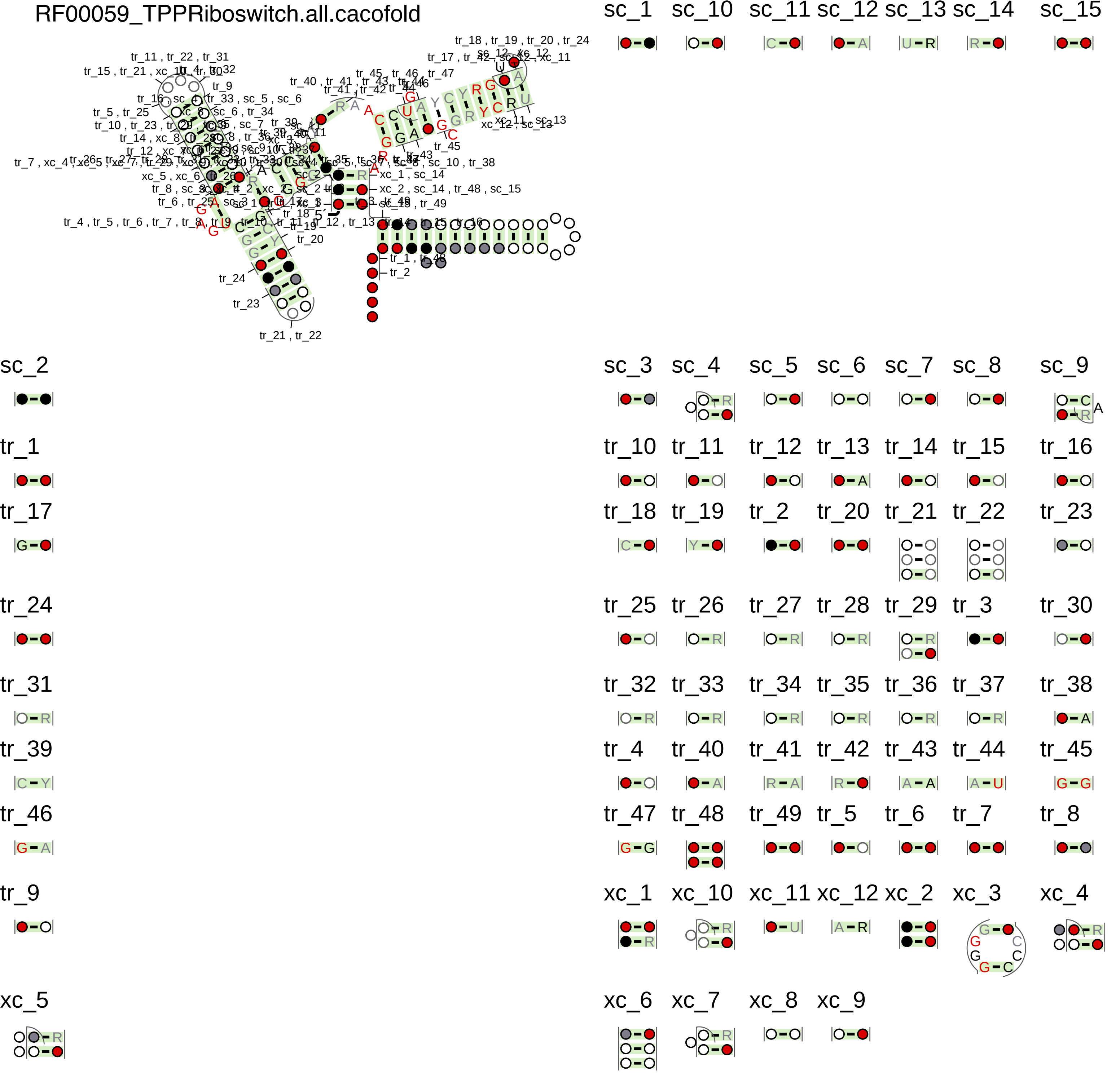

### RF00168_glmSRiboswitch.arnold.cacofold

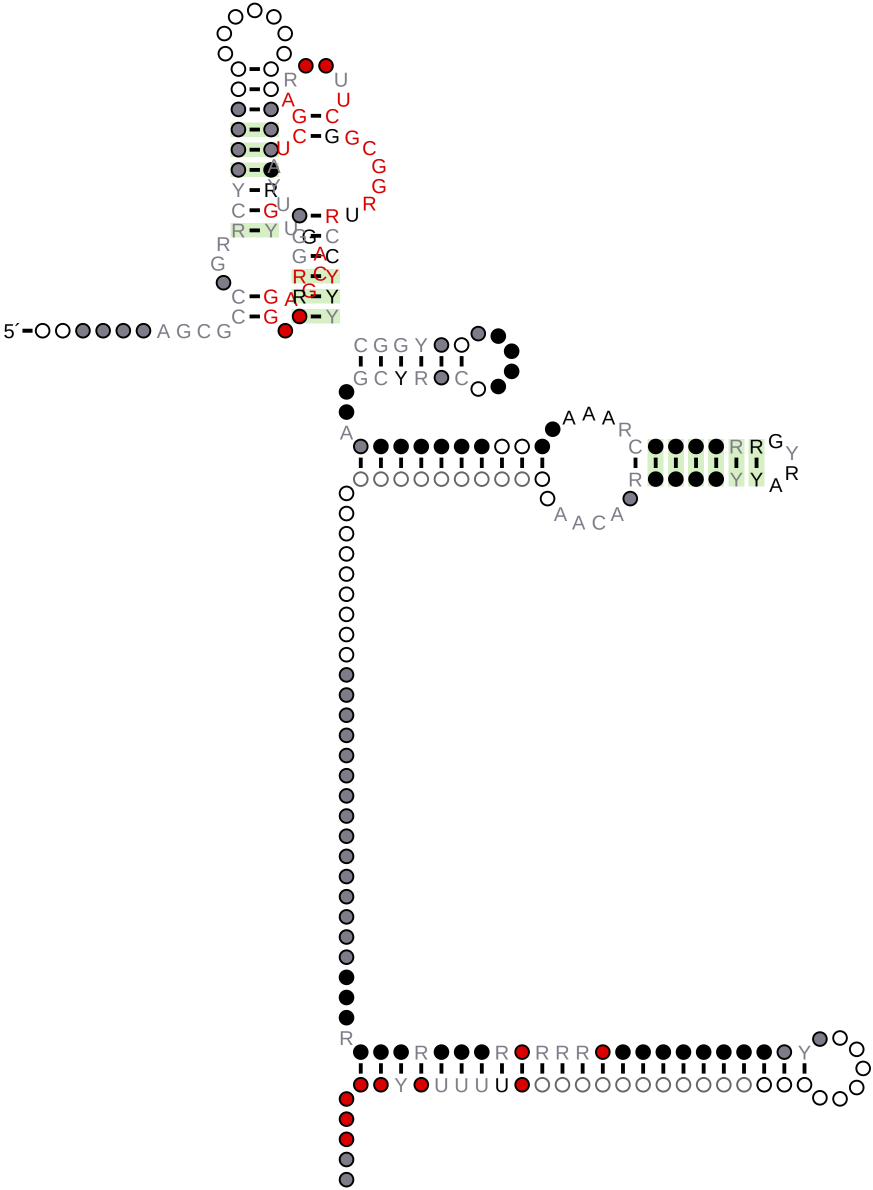

### RF00168_glmSRiboswitch.all.cacofold

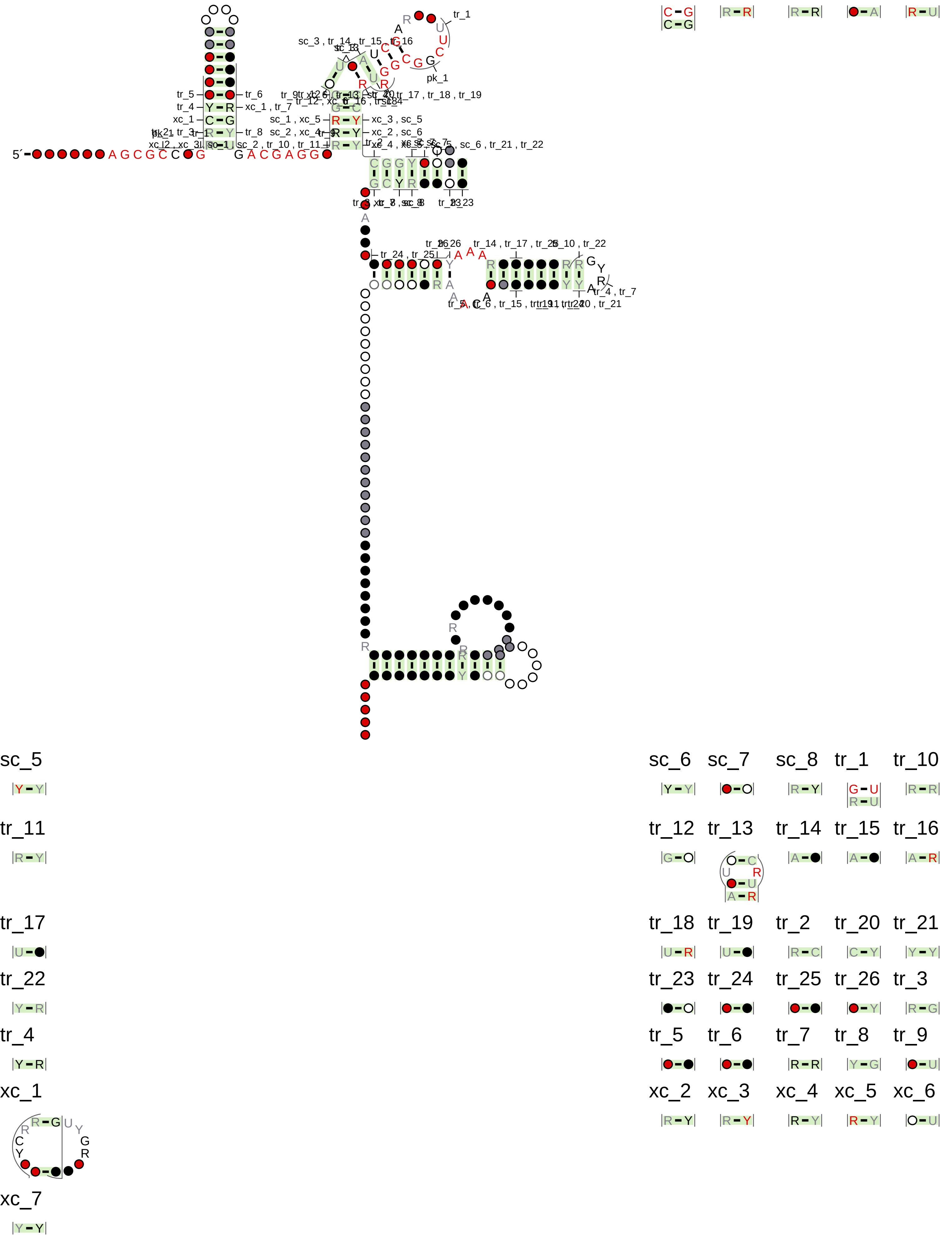
