## Supplementary material for "High-throughput functional profiling and evolutionary covariation analysis of entire riboswitch sequences": Unedited phylogenetic trees.

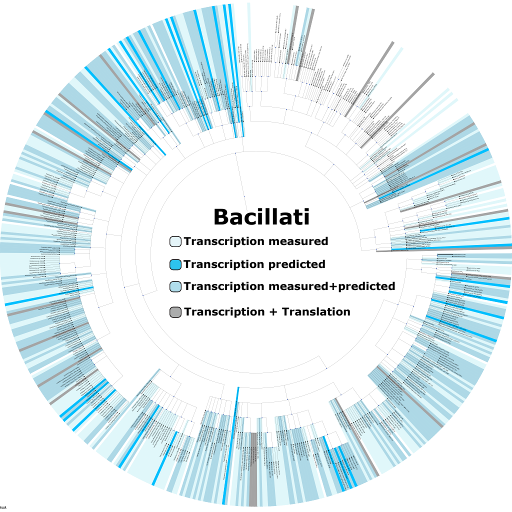

Bacillati
Transcription measured
Transcription predicted
Transcription measured+predicted
Transcription + Translation

### Fusobacteriati

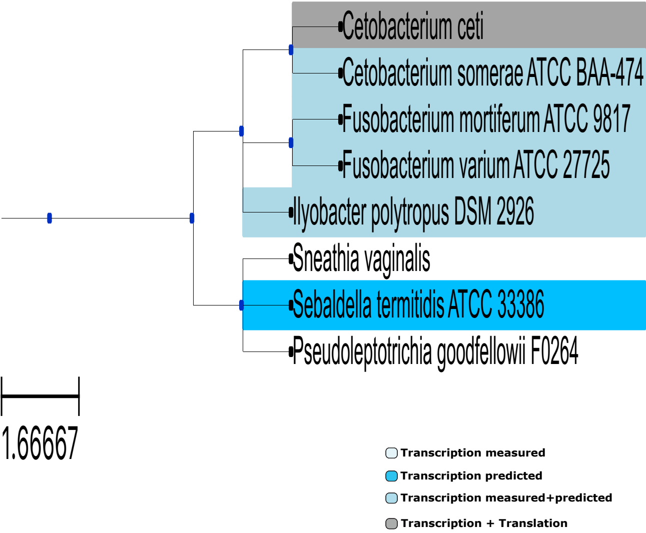

### Metazoa

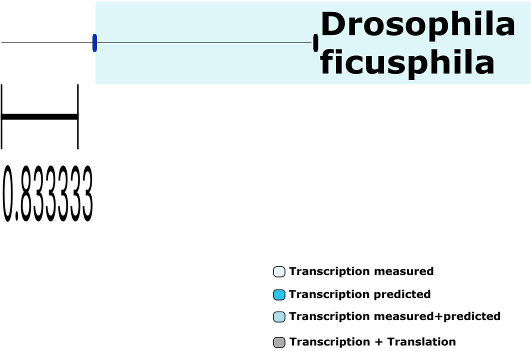

Drosophila ficusphila
0.833333
Transcription measured
Transcription predicted
Transcription measured+predicted
Transcription + Translation

### Methanobacteriati

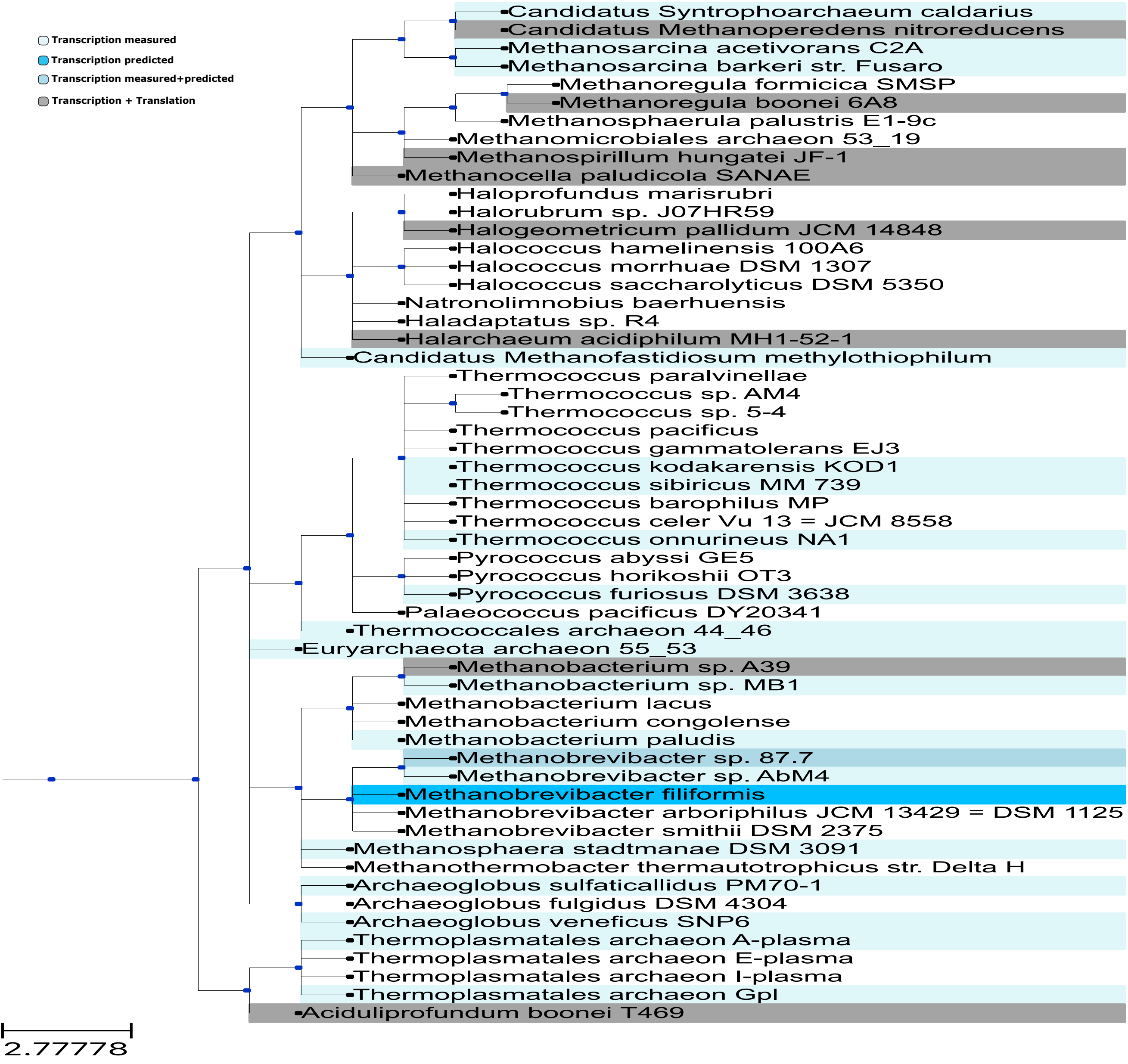

**Promethearchaeati**

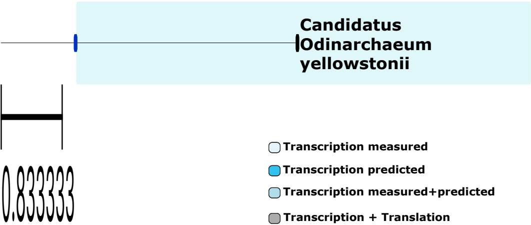

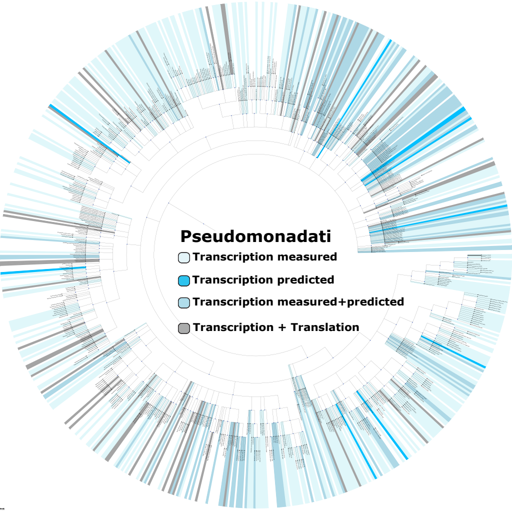
Pseudomonadati
Transcription measured
Transcription predicted
Transcription measured+predicted
Transcription + Translation

### Thermoproteati

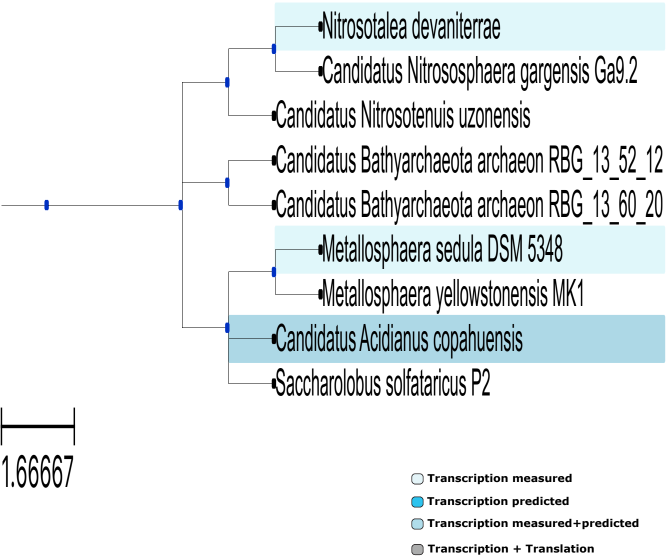

### Thermotogati

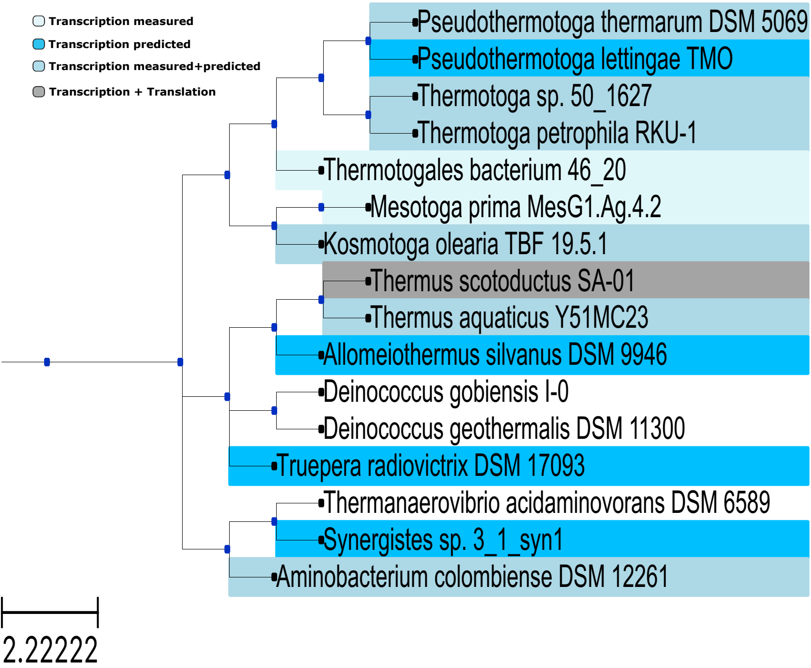

### Unknown

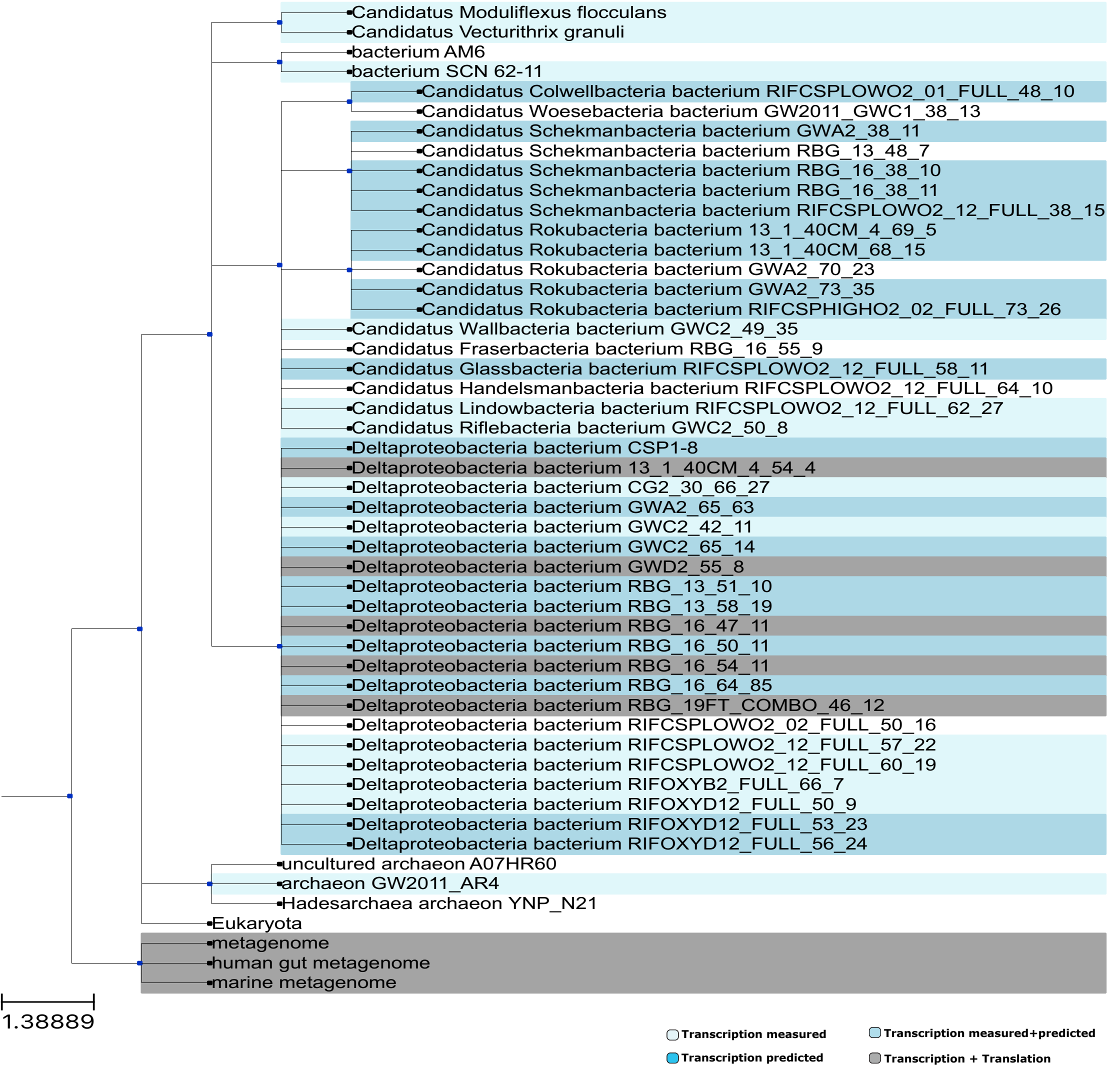

Candidatus Moduliflexus flocculans
Candidatus Vecturithrix granuli
bacterium AM6
bacterium SCN 62-11
Candidatus Colwellbacteria bacterium RIFCSPLOWO2_01_FULL_48_10
Candidatus Woesebacteria bacterium GW2011_GWC1_38_13
Candidatus Schekmanbacteria bacterium GWA2_38_11
Candidatus Schekmanbacteria bacterium RBG_13_48_7
Candidatus Schekmanbacteria bacterium RBG_16_38_10
Candidatus Schekmanbacteria bacterium RBG_16_38_11
Candidatus Schekmanbacteria bacterium RIFCSPLOWO2_12_FULL_38_15
Candidatus Rokubacteria bacterium 13_1_40CM_4_69_5
Candidatus Rokubacteria bacterium 13_1_40CM_68_15
Candidatus Rokubacteria bacterium GWA2_70_23
Candidatus Rokubacteria bacterium GWA2_73_35
Candidatus Rokubacteria bacterium RIFCSPHIGHO2_02_FULL_73_26
Candidatus Wallbacteria bacterium GWC2_49_35
Candidatus Fraserbacteria bacterium RBG_16_55_9
Candidatus Glassbacteria bacterium RIFCSPLOWO2_12_FULL_58_11
Candidatus Handelsmanbacteria bacterium RIFCSPLOWO2_12_FULL_64_10
Candidatus Lindowbacteria bacterium RIFCSPLOWO2_12_FULL_62_27
Candidatus Riflebacteria bacterium GWC2_50_8
Deltaproteobacteria bacterium CSP1-8
Deltaproteobacteria bacterium 13_1_40CM_4_54_4
Deltaproteobacteria bacterium CG2_30_66_27
Deltaproteobacteria bacterium GWA2_65_63
Deltaproteobacteria bacterium GWC2_42_11
Deltaproteobacteria bacterium GWC2_65_14
Deltaproteobacteria bacterium GWD2_55_8
Deltaproteobacteria bacterium RBG_13_51_10
Deltaproteobacteria bacterium RBG_13_58_19
Deltaproteobacteria bacterium RBG_16_47_11
Deltaproteobacteria bacterium RBG_16_50_11
Deltaproteobacteria bacterium RBG_16_54_11
Deltaproteobacteria bacterium RBG_16_64_85
Deltaproteobacteria bacterium RBG_19FT_COMBO_46_12
Deltaproteobacteria bacterium RIFCSPLOWO2_02_FULL_50_16
Deltaproteobacteria bacterium RIFCSPLOWO2_12_FULL_57_22
Deltaproteobacteria bacterium RIFCSPLOWO2_12_FULL_60_19
Deltaproteobacteria bacterium RIFOXYB2_FULL_66_7
Deltaproteobacteria bacterium RIFOXYD12_FULL_50_9
Deltaproteobacteria bacterium RIFOXYD12_FULL_53_23
Deltaproteobacteria bacterium RIFOXYD12_FULL_56_24
uncultured archaeon A07HR60
archaeon GW2011_AR4
Hadesarchaea archaeon YNP_N21
Eukaryota
metagenome
human gut metagenome
marine metagenome
1.38889
Transcription measured
Transcription measured+predicted
Transcription predicted
Transcription + Translation

### Viridiplantae

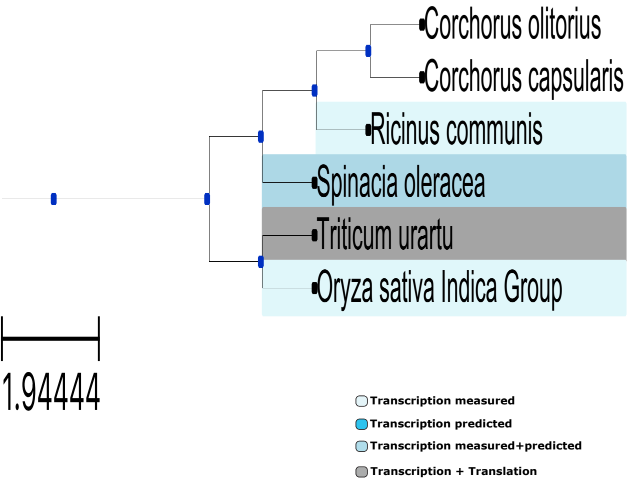
